# Defining RNA oligonucleotides that reverse deleterious phase transitions of RNA-binding proteins with prion-like domains

**DOI:** 10.1101/2023.09.04.555754

**Authors:** Lin Guo, Jacob R. Mann, Jocelyn C. Mauna, Katie E. Copley, Hejia Wang, Jack D. Rubien, Hana M. Odeh, JiaBei Lin, Bo Lim Lee, Laura Ganser, Emma Robinson, Kevin M. Kim, Anastasia C. Murthy, Tapas Paul, Bede Portz, Amanda M. Gleixner, Zamia Diaz, Jenny L. Carey, Ashleigh Smirnov, George Padilla, Ellen Lavorando, Carolann Espy, Yulei Shang, Eric J. Huang, Alessandra Chesi, Nicolas L. Fawzi, Sua Myong, Christopher J. Donnelly, James Shorter

**Author notes:** Co-first authors (Lin Guo, Jacob R. Mann). Co-second authors (Jocelyn C. Mauna, Katie E. Copley, Hejia Wang, Jack D. Rubien).

## Abstract

RNA-binding proteins with prion-like domains, such as FUS and TDP-43, condense into functional liquids, which can transform into pathological fibrils that underpin fatal neurodegenerative disorders, including amyotrophic lateral sclerosis (ALS)/frontotemporal dementia (FTD). Here, we define short RNAs (24-48 nucleotides) that prevent FUS fibrillization by promoting liquid phases, and distinct short RNAs that prevent and, remarkably, reverse FUS condensation and fibrillization. These activities require interactions with multiple RNA-binding domains of FUS and are encoded by RNA sequence, length, and structure. Importantly, we define a short RNA that dissolves aberrant cytoplasmic FUS condensates, restores nuclear FUS, and mitigates FUS proteotoxicity in optogenetic models and human motor neurons. Another short RNA dissolves aberrant cytoplasmic TDP-43 condensates, restores nuclear TDP-43, and mitigates TDP-43 proteotoxicity. Since short RNAs can be effectively delivered to the human brain, these oligonucleotides could have therapeutic utility for ALS/FTD and related disorders.

## Introduction

There are no effective therapeutics for several devastating neurodegenerative disorders, including amyotrophic lateral sclerosis (ALS) and frontotemporal dementia (FTD). A common feature of these disorders is the cytoplasmic mislocalization and aggregation of nuclear RNA-binding proteins (RBPs) with prion-like domains (PrLDs), such as TDP-43 or FUS, in degenerating neurons.^1–7^ This cytoplasmic aggregation is driven by PrLDs, which are distinctive low-complexity domains with an amino-acid composition enriched for uncharged polar residues (especially glutamine, asparagine, tyrosine, and serine) and glycine similar to yeast prion domains.^1^ Yeast prion domains enable various yeast proteins (e.g. Sup35 and Ure2) to form prions.^1^ In the context of TDP-43 and FUS, the PrLD renders these RBPs intrinsically aggregation prone and enables the formation of self-templating, amyloid-like fibrils.^1,3,8–10^ However, the PrLD also enables TDP-43 and FUS to undergo phase separation (PS), where TDP-43 and FUS can spontaneously condense from dispersed states in solution to a separated liquid phase.^1^ PS enables TDP-43 and FUS to function in membraneless organelles inside the nucleus.^1,11–14^ TDP-43 and FUS can also undergo PS in the cytoplasm during recruitment to stress granules and during formation of alternative cytoplasmic condensates.^1,11,12,15,16^ If TDP-43 or FUS dwell in liquid states for too long, especially in the cytoplasm, then they can transition into amyloid-like fibrils.^11,15,16^ This switch from a liquid to pathological fibrils in the context of disease is termed an aberrant phase transition, which can be accelerated by ALS-linked mutations in TDP-43 or FUS.^11,15^ A deleterious event in ALS/FTD occurs when TDP-43 or FUS become depleted from the nucleus and trapped in cytoplasmic aggregates.^1,12,15^

A key therapeutic innovation for ALS/FTD would be to develop agents that reverse the aberrant cytoplasmic aggregation of TDP-43 and FUS, and return these proteins to native form and nuclear function.^1^ Such agents would simultaneously eliminate two malicious problems associated with cytoplasmic TDP-43 or FUS aggregation: **(1)** the toxic gain of function of cytoplasmic aggregated TDP-43 or FUS conformers; and **(2)** the loss of cytoplasmic and nuclear function of soluble TDP-43 or FUS due to sequestration in cytoplasmic aggregates. These two issues likely synergize in the etiology of various forms of ALS/FTD.^1^

A therapeutic disaggregase that dissolves cytoplasmic aggregates and restores TDP-43 or FUS back to the nucleus could eradicate these two deleterious phenotypes simultaneously.^17^ Previously, we have established that engineered versions of yeast Hsp104 and endogenous human nuclear-import receptors can reverse TDP-43 or FUS aggregation and restore the RBPs to the nucleus.^8,18,19^ However, despite exciting advances in AAV technology,^20^ delivering these agents into the degenerating neurons of ALS/FTD patients remains a significant challenge.

Here, we define more deliverable therapeutic agents: short RNA oligonucleotides (24-48 nucleotides [nts]) that antagonize FUS fibrillization. We uncover two distinct classes of RNA inhibitor. Weak RNA inhibitors prevent FUS fibrillization by promoting liquid states. By contrast, strong RNA inhibitors prevent and reverse FUS condensation and fibrillization. These activities require interactions with multiple RNA-binding domains of FUS and are encoded by RNA sequence, length, and structure. Importantly, we define a short RNA (25nts) that dissolves aberrant FUS condensates, restores FUS to the nucleus, and mitigates FUS proteotoxicity in optogenetic models and in human motor neurons. A distinct short RNA oligonucleotide (34nts) can prevent and reverse TDP-43 PS and fibrillization by engaging the TDP-43 RNA-recognition motifs (RRMs). This RNA dissolves aberrant cytoplasmic TDP-43 condensates, restores TDP-43 to the nucleus, and mitigates TDP-43 toxicity. Thus, we establish an important concept: specific short RNAs can prevent *and* reverse aberrant phase transitions of TDP-43 and FUS to restore nuclear localization and mitigate neurotoxicity. Since short RNA oligonucleotides can be readily delivered to the human brain,^21^ these agents could have therapeutic utility for ALS/FTD and related disorders.

## Results

### Identification of short RNAs that strongly or weakly antagonize FUS assembly

First, we aimed to identify short specific RNAs (24-48nts) that antagonize aberrant FUS assembly. We sought RNAs of this length as they can be readily delivered to the CNS of patients akin to antisense oligonucleotides (ASOs), which are typically 12-30nts.^22^ While extracting recombinant GST-FUS from *E. coli*, we found that purified GST-FUS was bound to RNA (Figure S1A). Specific removal of the GST tag with TEV protease elicits rapid assembly of tangled FUS fibrils within an hour.^8,10^ By contrast, GST-FUS remains soluble in the absence of TEV cleavage over this timeframe.^10^ Strikingly, treating GST-FUS with RNase A to remove RNA strongly increased turbidity in the absence of TEV cleavage (Figure S1B). Hence, RNA promotes GST-FUS solubility.

We sought to identify enriched RNA motifs in this FUS-bound RNA population as they might antagonize aberrant FUS assembly. The A_260_/A_280_ ratio of the GST-FUS purified from *E. coli* was ∼1.8, indicating ∼38% (w/w) nucleic acid in the sample.^23^ RNA extraction followed by electrophoresis revealed a population of RNAs with a size range of ∼50-100nts. From this RNA population, a cDNA library was constructed and sequenced. We identified 42 enriched motifs between 8-12nts in our library (Table S1). 14 RNA motifs were selected for further testing based on their enrichment and sequence diversity (Table S1, S2). RNA oligonucleotides (oligos) containing 2 or 4 repeats of individual enriched motifs were synthesized and assessed for their ability to inhibit FUS fibrillization (Figure 1A; Table S2). An RNA oligo (RNA C2) that does not bind FUS effectively was used as a negative control (Table S2).^24^ Indeed, RNA C2 had little effect on FUS fibrillization (Figure 1B). Remarkably, RNA S2 (containing 4 repeats of GAGGUGGCUAUG) diminished FUS fibrillization (Figure 1B). Thus, a short RNA bearing specific motifs that engage FUS can abrogate FUS fibrillization.

**Figure 1.**
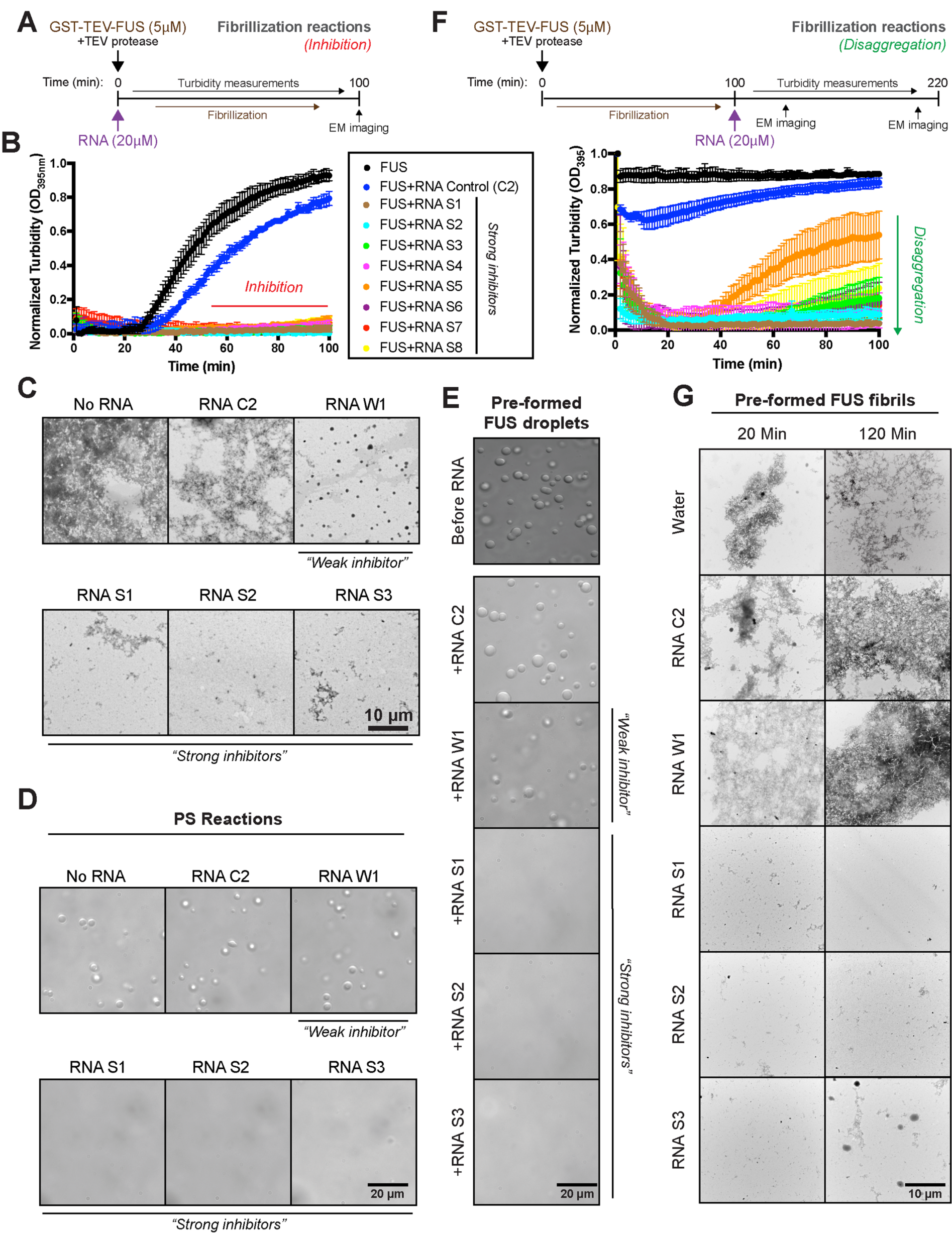
Strong RNA inhibitors inhibit and reverse FUS fibrillization and PS. **(A)** Schematic of experiments to test whether RNA oligos inhibit fibrillization. GST-FUS (5µM) was incubated with TEV protease in the presence or absence of RNA oligos (20µM) for 0–100min. Turbidity measurements were taken every minute to assess the extent of fibrillization. Samples were taken at the end of the reaction to visualize FUS structures via EM. **(B, C)** GST-FUS (5µM) was incubated with TEV protease in the presence or absence of RNA (20µM) for 0– 100min. Fibrillization was assessed by turbidity (B) or EM (C). Bar, 10μm. Data shown in (B) are means±SEM (n=3). **(D)** GST-FUS (10µM) was incubated for 4h in the presence or absence of the indicated RNA (40µM). Droplet formation was assessed by DIC microscopy. Bar, 20μm. **(E)** GST-FUS (10µM) droplets were incubated with the indicated RNA (40µM) for 10min. Droplet integrity was assessed by DIC microscopy. Bar, 20μm. **(F, G)** Schematic of experiments to test whether RNA oligos reverse FUS fibrillization. GST-FUS (5µM) was incubated with TEV protease for 100min to form fibrils. At this time, water, or RNA (20µM) was added. Disaggregation was assessed by turbidity (F). Data shown in (F) are means±SEM (n=3-4). Samples were taken after 20min and 120min to visualize FUS structures via EM **(G)**. Bar, 10μm. See also **Figure S1** and **S2**.

We found seven additional RNAs (RNA W1, W4-W9) that increased the lag time and reduced the extent of FUS assembly more than RNA C2 (Figure S1C, D, Table S2). By contrast, six RNAs (RNA N1-N6) based on enriched motifs from our library were ineffective at inhibiting FUS assembly and did not show a significant difference compared to RNA C2 (Figure S1E, F, Table S2). Thus, the presence of RNA in general is not sufficient to inhibit FUS fibrilization. Rather, specific RNAs likely have different abilities to antagonize FUS fibrillization.

We next tested whether RNAs that have been reported to bind FUS might also antagonize FUS fibrillization. Eight short RNA sequences that bind FUS (RNA S1, S3-S8, and W3) as well as a (UG)_6_-containing RNA (RNA W2) that binds TDP-43 were tested (Table S2).^25–28^ Within this group of RNAs, seven oligos (RNA S1, S3-S8) strongly inhibited FUS fibrillization (Figure 1B). By contrast, RNA W2 and the GGUG-containing RNA (RNA W3) mildly inhibited FUS assembly (Figure S1C, D). Based on these findings, we classified RNAs into three groups: strong inhibitors (RNA S1-S8) that reduced turbidity by more than 90%, weak inhibitors (RNA W1-W9) that reduced turbidity by less than 90% but significantly compared to control RNA C2, and non-effective RNAs (RNA N1-N6) that had no effect beyond control RNA C2 (Figure 1B, S1C-F, Table S2).

### Strong RNA inhibitors prevent FUS PS and fibrillization, whereas weak RNA inhibitors allow FUS PS but prevent fibrillization

Next, we assessed how strong and weak RNA inhibitors affected FUS assembly via sedimentation analysis. Strong RNA inhibitors (RNA S1-S3) promoted accumulation of FUS in the soluble fraction, whereas a weak RNA inhibitor (RNA W1) did not (Figure S1G). Thus, weak RNA inhibitors may allow FUS to assemble into structures that display reduced turbidity compared to large tangles of FUS fibrils. FUS can form liquid droplets that later convert into fibrils.^11,29^ Thus, we wondered whether strong and weak RNA inhibitors might antagonize different stages of this process. We utilized electron microscopy (EM) to visualize FUS assemblies formed in the presence or absence of strong and weak RNA inhibitors. In the absence of RNA, FUS forms large aggregates comprised of tangled fibrils (Figure 1C).^8,10^ FUS fibrillization was unaffected by the negative control RNA C2 (Figure 1C). By contrast, strong inhibitors (RNA S1-S5) greatly reduced the formation of any FUS assemblies (Figure 1C and S1H). Interestingly, a weak RNA inhibitor, RNA W1, inhibited the formation of large tangles of FUS fibrils, but allowed the formation of numerous spherical structures, indicative of phase-separated condensates (Figure 1C). Thus, weak RNA inhibitors may prevent FUS fibrillization by promoting liquid phases.

To explore this possibility further, we employed Differential Interference Contrast (DIC) microscopy to more closely study the formation and dynamics of FUS droplets in the presence or absence of strong and weak RNA inhibitors. If we do not remove the GST tag, then GST-FUS is initially soluble, but slowly condenses into liquid droplets after several hours.^8^ Indeed, without addition of RNA or in the presence of RNA C2, GST-FUS formed dynamic droplets that exhibited classic liquid-like behavior such as fusion and surface wetting (Figure 1D).^8,11^ Strong RNA inhibitors abolished formation of FUS droplets (Figure 1D and Figure S1I), whereas weak RNA inhibitors had no effect on FUS droplet formation (Figure 1D and Figure S1I, RNA W1 and W2). Thus, strong RNA inhibitors prevent FUS PS and fibrillization, whereas weak RNA inhibitors allow FUS PS but prevent fibrillization.

### Strong RNA inhibitors reverse FUS PS and fibrillization

Next, we assessed whether short RNAs could reverse the formation of preformed FUS droplets. The weak RNA inhibitors, RNA W1-W4, and control RNA C2 had no effect on preformed FUS droplets (Figure 1E, Movie S1). Remarkably, strong RNA inhibitors (RNA S1-S3) rapidly dissolved preformed FUS droplets (Figure 1E, Movie S2-S4). Thus, strong RNA inhibitors can reverse FUS PS, whereas weak RNA inhibitors cannot.

We next tested whether RNA inhibitors could disassemble preformed FUS fibrils (Figure 1F). Addition of solvent or RNA C2 had little effect on FUS fibrils (Figure 1F, G). By contrast, strong RNA inhibitors rapidly disassembled preformed FUS fibrils within 20 minutes (Figure 1F). Among the strong RNA inhibitors, RNA S2 disassembled FUS fibrils most rapidly, whereas RNA S1 showed the most complete disaggregation (Figure 1F). Thus, remarkably, strong RNA inhibitors can rapidly disassemble preformed FUS fibrils.

Interestingly, when other strong RNA inhibitors (RNA S3-S8) were added to FUS fibrils, we observed an initial rapid disassembly followed by slow recovery of turbidity upon further incubation (Figure 1F). EM revealed that 20 minutes after addition of RNAs, when turbidity is at the lowest point, all strong RNA inhibitors effectively disassembled FUS fibrils (Figure 1G and S2A). However, 2 hours after addition of RNAs, when turbidity increased again, dense FUS condensates were observed for samples treated with RNA S3, RNA S4, and RNA S5 (Figure 1G, S2A). These dense FUS condensates exhibited porous architecture resembling a hydrogel (Figure S2A).^8^ DIC microscopy revealed that these FUS condensates were spherical but did not fuse, indicating a gel-like phase (Figure S2B). Thus, a subset of strong RNA inhibitors disassembles FUS fibrils initially, but FUS then transforms into dense gel-like condensates. By contrast, RNA S1 and S2 are unusual in that they effectively dissolve FUS fibrils and do not transform FUS into another condensate.

Most weak RNA inhibitors (i.e., RNA W1, W4-W9) had no effect on preformed FUS fibrils beyond the negative control RNA C2 (Figure S2C). By contrast, RNA W2 and W3 initially reduced turbidity, but turbidity then returned to levels observed with RNA C2, indicating a lack of a sustained effect (Figure S2C). The non-effective RNAs (Table S2) did not have any effect on preformed FUS fibrils beyond RNA C2 (Figure S2D). Thus, the disassembly of FUS fibrils is due to binding to specific RNA sequences. Our findings suggest that RNA S1 and S2 possess an unusual ability to dissolve FUS liquids and fibrils. RNA S1 is a natural FUS-binding RNA (CUAGGAUGGAGGUGGGGAAUGGUAC) found in the 3’UTR of the BDNF gene.^26^ By contrast, RNA S2 is a synthetic RNA comprised of four repeats of an RNA motif (GAGGUGGCUAUG) found to engage FUS during purification from *E. coli*. Thus, both native and designed RNA sequences that engage FUS can reverse FUS PS and fibrillization.

### RNA S1 prevents and reverses assembly of ALS-linked FUS mutants

Mutations in FUS are an established cause of ALS.^30^ Thus, we next tested whether RNA S1 could also antagonize assembly of ALS-linked FUS variants, including FUS^P525L^, FUS^R521G^, FUS^R244C^, and FUS^R216C^. Importantly, RNA S1 inhibited (Figure S2E) and reversed fibrillization of these disease-linked FUS variants (Figure S2F-I). Thus, RNA S1 can prevent and reverse fibrillization of FUS as well as several ALS-linked FUS variants.

### RNA length, sequence, and structure determine ability to antagonize FUS fibrillization

We next investigated what features determined whether an RNA was a strong or weak inhibitor. Strong RNA inhibitors were typically longer than weak inhibitors (Table S2, Figure S2J). Most weak RNA inhibitors or non-effective RNAs were 24-28nts, whereas most strong inhibitors were 39-48nts (Figure S2J, Table S2). The two exceptions were RNA S1 and S7 (Table S2), which were 24-25nts (Table S2). To examine how RNA length affected the ability of RNA oligos to antagonize FUS assembly, we selected a strong inhibitor RNA S2, which is 4 repeats of the enriched motif GAGGUGGCUAUG, and synthesized RNA S2/2, which contains 2 repeats of the same enriched motif (Table S2, Figure 2A). Shortening the length of strong inhibitor RNA S2 reduced its ability to prevent FUS assembly (Figure 2B). This effect was more pronounced for reversing FUS fibrillization. Thus, RNA S2 effectively reversed FUS fibrillization, whereas RNA S2/2 had no effect beyond the negative control RNA C2 (Figure 2C). Thus, increasing RNA length from 24 to 48nts can enable more effective prevention and reversal of FUS fibrillization.

**Figure 2.**
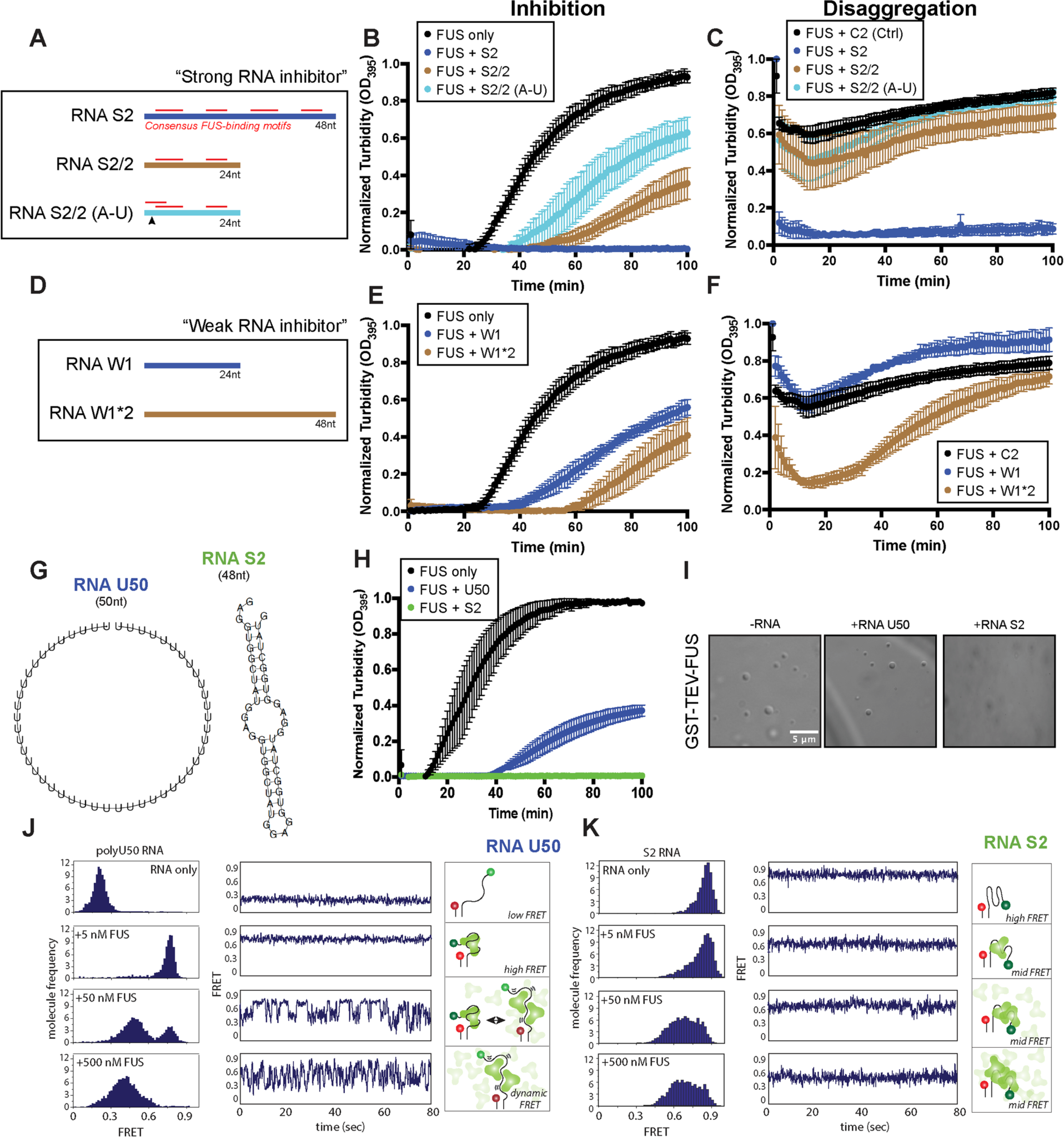
RNA oligo length, sequence, and structure determine ability to prevent and reverse FUS fibrillization. **(A)** Schematic of strong inhibitor RNA S2, which contains 4 repeats of the enriched motif GAGGUGGCUAUG, and RNA S2/2, which contains 2 repeats of the same enriched motif. An A to U mutation was introduced in RNAS 2/2 (arrowhead) to evaluate the effect of RNA sequence. The red bars represent the consensus FUS-binding motif, which is GGUG in RNA S2. The A to U mutation on RNA S2/2 creates overlapping GUGG FUS-binding motifs. **(B)** GST-FUS (5µM) was incubated with TEV protease in the presence or absence of indicated RNA (20µM) for 0–100min. Fibrillization was assessed by turbidity. Values are means±SEM (n=3-4). **(C)** GST-FUS (5µM) was incubated with TEV protease for 100min to form fibrils. At this time, the indicated RNA (20µM) was added. Disaggregation was assessed by turbidity. Values are means±SEM (n=3-4). **(D)** Schematic of weak inhibitor RNA W1, which contains two repeats of enriched motif UCAGAGACAUCA, and RNA W1*2, which doubles the length of RNA W1 and contains 4 repeats of the enriched motif. **(E)** GST-FUS (5µM) was incubated with TEV protease in the presence or absence of indicated RNA (20µM) for 0– 100min. FUS assembly was assessed by turbidity. The FUS only curve was plotted from the same data set as in (B), since experiments in (B) and (E) were run at the same time. Values are means±SEM (n=3-4). **(F)** GST-FUS (5µM) was incubated with TEV protease for 100min to form fibrils. At this time, the indicated RNA (20µM) was added. Disaggregation was assessed by turbidity. The FUS+C2 curve was plotted from the same data set as in (C), since experiments in (C) and (F) were run at the same time. Values are means±SEM (n=3-5). **(G)** Predicted secondary structure of U50 and RNA S2 by RNAfold.^80^ **(H)** GST-FUS (5µM) was incubated with TEV protease in the presence or absence of RNA U50 (blue) or RNAS2 (green) (20µM) for 0– 100min. FUS assembly was assessed by turbidity. Values are means±SEM (n=3). **(I)** GST-FUS (10µM) was incubated for 4 hours in the presence or absence of the indicated RNA (40µM). Droplet formation was assessed by DIC microscopy. Bar, 20μm. **(J, K)** smFRET histograms and representative traces for increasing FUS concentrations (0-500nM) (left) and schematic of the smFRET experiment in which Cy3 and Cy5 are attached to either end of RNA to report on the conformational changes induced by FUS binding (right) for RNA U50 (J) and RNA S2 (K).

To further determine the effect of RNA length on activity, we next focused on potentiating the weak inhibitor RNA W1, which contains two repeats of enriched motif UCAGAGACAUCA. We synthesized RNA W1*2, which doubles the length of RNA W1 and contains 4 repeats of the enriched motif (Figure 2D). Doubling the length of RNA W1 increased its ability to prevent FUS assembly (Figure 2E). RNA W1*2 was also more effective than RNA W1 in reversing FUS fibrillization in the initial 20min of the reaction (Figure 2F), but turbidity increased at later times (Figure 2F). Nonetheless, increasing RNA length from 24 to 48nts can enable more effective prevention and reversal of FUS assembly. Indeed, strong inhibitors were typically longer than weak inhibitors (Figure S2J).

RNA S2 and RNA W1*2 are the same length (48nt), but RNA S2 prevents and reverses FUS assembly more effectively than RNA W1*2 (Figure 2B, E). RNA S2 contains four consensus FUS-binding motifs, i.e., GGUG,^24^ whereas these FUS-binding motifs are absent from RNA W1*2 (Figure 2A). Thus, the precise RNA sequence is also important for activity. Strong RNA inhibitors tend to have more GGU and GG sequences than weak RNA inhibitors (Figure S2K, Table S2). Notably, the two strongest inhibitors RNA S1 and S2 have AUGGAGGUGG in their sequence. To further explore how sensitive sequence requirements might be, we introduced a single A to U mutation in RNA S2/2 (to yield RNA S2/2 (A-U)), which creates overlapping GUGG FUS-binding motifs (Table S2, Figure 2A).^28^ This single mutation reduced the ability of RNA S2/2 (A-U) to prevent FUS assembly (Figure 2A, B). RNA S2/2 and RNA S2/2 (A-U) have similar predicted secondary structures (Table S2). Thus, specific RNA sequences can encode more effective inhibition of FUS fibrillization.

We next considered whether RNA structure might also contribute to preventing and reversing FUS assembly. Thus, we employed single molecule Fӧrster Resonance Energy Transfer (smFRET) to study the conformation of RNA and the interaction between FUS and RNA.^31^ Here, we examined an unstructured RNA (U50) and a strong RNA inhibitor with similar length (RNA S2). RNA U50 is predicted to be unstructured, whereas RNA S2 is predicted to adopt a stem-loop structure with folding energy of −16.40 kcal/mol (Figure 2G). Unstructured RNA U50 was a weak RNA inhibitor that reduced FUS assembly (Figure 2H) but did not affect FUS PS (Figure 2I). For smFRET, RNA U50 or RNA S2 was immobilized onto a PEG-passivated quartz slide via an 18-bp duplex and biotin-neutravidin interaction (Figure 2J, K).^31,32^ Cy3 and Cy5 were attached to either end of each RNA to report on the conformational status of RNA and the change induced by FUS binding (Figure 2J, K). The FRET value for U50 in the absence of FUS is ∼0.2, consistent with an unstructured RNA.^31^ Conversely, the FRET value for RNA S2 is ∼0.8, indicating a stable, folded RNA conformation (Figure 2K), consistent with MFold predictions.

Addition of FUS to the RNA resulted in FRET changes which report on how FUS binding affects RNA conformation (Figure 2J, K). For U50, addition of low FUS concentration (5nM) immediately shifted the low FRET (∼0.2) to a single high FRET peak (∼0.8) with single molecule traces displaying a stable high FRET signal (Figure 2J).^31^ Thus, FUS induces a tight compaction of the long, unstructured U50 RNA (Figure 2J). As FUS concentration increased (50 and 500nM), the high FRET population diminished, and a broad mid FRET peak (∼0.5) emerged with smFRET traces showing increased fluctuations (Figure 2J).^31^ The mid FRET peak indicates an extended RNA structure, which allows dynamic interaction between FUS multimers and a single RNA (Figure 2J).^31^ This finding is consistent with U50 allowing FUS droplets to form (Figure 2I). Thus, the highly dynamic interaction between U50 and FUS (50 and 500nM) is consistent with the dynamic nature of FUS liquid droplets.

By contrast, addition of FUS to RNA S2 did not yield dynamic FRET fluctuations (Figure 2K), indicating formation of a static complex. FUS binding to RNA S2 induced a lower (∼0.6) FRET population, suggesting that FUS partially unfolds RNA S2 upon binding (Figure 2K). The FUS-bound peak is wider than the free RNA peak, indicating that the conformation of RNA S2 is more heterogeneous when bound to FUS (Figure 2K). Nonetheless, unlike U50, the structured nature of RNA S2 restricted FUS to a static complex that precluded the formation of dynamic FUS multimers (Figure 2K, right panel). Thus, RNA S2 eliminates FUS PS and fibrillization by restricting FUS to a static complex.

### Strong RNA inhibitors engage the FUS RRM to antagonize FUS assembly

We next assessed how the individual RNA-binding domains of FUS enable short RNAs to exert their effects. The FUS RRM, Zinc finger domain, and RGG domains can all contribute to RNA binding.^33^ We selected the three RNAs with the strongest *in vitro* activities (i.e., RNA S1, RNA S2, and RNA S3) and one weak inhibitor (RNA W1) for further analysis. Strong RNA inhibitors bind to FUS tightly (RNA S1: *K_D_*∼40.8nM; RNA S2: *K_D_*∼105nM; RNA S3: *K_D_*∼102nM) (Figure 3A), whereas RNA W1 binds to FUS, but with a *K_D_* greater than 3μM (value could not be determined via fluorescence anisotropy). Thus, tighter binding may enable the activity of strong RNA inhibitors.

**Figure 3.**
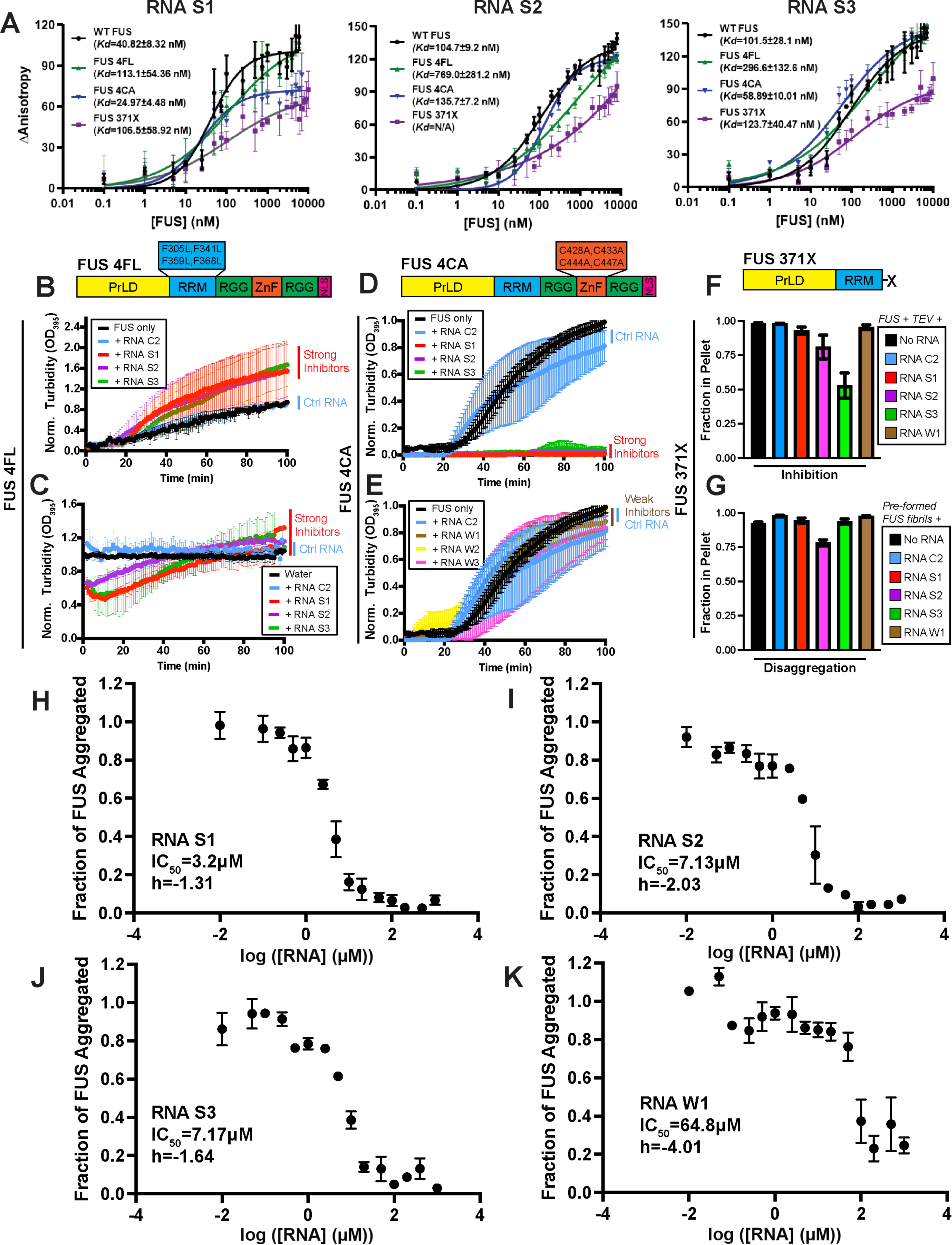
Strong and weak RNA inhibitors engage multiple RNA-binding domains of FUS to antagonize FUS fibrillization. **(A)** Change of anisotropy when the indicated fluorescein-labeled RNA (8nM) binds to GST-FUS, GST-FUS_4F-L_, GST-FUS_4C-A_, or GST-FUS_371X_ at the indicated concentrations. Values represent means±SEM (n = 3). Binding curves were fitted by Prism. Solid line represents the fit and the fitted *K_D_* is listed. **(B)** GST-FUS_4F-L_ (5µM) was incubated with TEV protease in the presence or absence of strong RNA inhibitors S1, S2, or S3 or the control C2 RNA (20µM) for 0–100min. Fibrillization was assessed via turbidity. Values represent means±SEM (n = 3). **(C)** FUS_4F-L_ fibrils (5µM monomer) were treated with water or the indicated RNA (20µM). Disaggregation was assessed by turbidity. Values represent means±SEM (n = 2-3). **(D)** GST-FUS_4C-A_ (5µM) was incubated with TEV protease in the presence or absence of strong RNA inhibitors S1, S2, and S3 or the control C2 RNA (20µM) for 0–100min. Fibrillization was assessed via turbidity. Values represent means±SEM (n = 3). **(E)** GST-FUS_4C-A_ (5µM) was incubated with TEV protease in the presence or absence of weak RNA inhibitors W1, W2, or W3 or the control C2 RNA (20µM) for 0–100 min. Fibrillization was assessed via turbidity. The FUS only and RNA C2 curves were plotted from the same data set as in (D), since experiments in (D) and (E) were run at the same time. Values represent means±SEM (n = 3). **(F)** GST-FUS_371X_ (10µM) was incubated with TEV protease in the presence or absence of the indicated RNA (40µM) at 25°C for 24h with agitation at 1200rpm. Aggregated FUS was quantified by sedimentation assay. Values represent means±SEM (n = 3). **(G)** FUS_371X_ fibrils (10µM monomer) were treated with water or indicated RNA (40µM) for 24h. Aggregated FUS was quantified by sedimentation assay. Values represent means±SEM (n = 3). **(H-K)** GST-FUS (5µM) was incubated with TEV protease in the presence or absence of **(H)** RNA S1, **(I)** RNA S2, **(J)** RNA S3, or **(K)** RNA W1 at indicated concentration for 0–100min. Fibrillization was assessed via turbidity. The dose response curves were fit by Prism using the log(inhibitor) vs. response -- Variable slope function. Values represent means±SEM (n = 3). See also **Figure S3**.

To assess the contribution of the FUS RRM to binding RNA S1, S2, and S3, we employed FUS_4F-L_ where four conserved phenylalanines (F305, F341, F359, and F368) in the RRM are mutated to leucine, which greatly reduces RNA binding.^34^ FUS_4F-L_ exhibited ∼2.8-7.3-fold reduced binding affinity to strong RNA inhibitors (RNA S1: *K_D_*∼113nM; RNA S2: *K_D_*∼769nM; RNA S3: *K_D_*∼297nM) (Figure 3A). Thus, FUS_4F-L_ can still bind RNA S1, S2, and S3, but with reduced affinity, indicating an important role for the FUS RRM in engaging these RNAs.

We next assessed whether strong (S1-S3) and weak (W1-W3) RNA inhibitors could prevent and reverse FUS_4F-L_ fibrillization. FUS_4F-L_ formed tangled fibrils, but these assembled more slowly than FUS (Figure 1B, 3B, S3A). None of the short RNAs tested here could prevent FUS_4F-L_ fibrillization (Figure 3B, S3A, B). Likewise, RNAs S1-S3 and W1-W3 were ineffective at reversing FUS_4F-L_ fibrillization (Figure 3C, S3C). Although turbidity was reduced in the first 20min by RNAs S1-S3, this effect was not sustained, and turbidity returned to initial levels (Figure 3C). Thus, strong RNA inhibitors must engage the FUS RRM to effectively prevent and reverse FUS fibrillization.

FUS_4F-L_ fibrillization could not be antagonized by RNAs S1 or S2. However, these RNAs could still bind to FUS_4F-L_, albeit with reduced affinity. To assess which other FUS domains might engage these RNAs, we used nuclear magnetic resonance (NMR) spectroscopy. Since the FUS PrLD does not bind to RNA,^35^ we employed FUS_269-454_, which lacks the N-terminal PrLD, but contains the RRM (residues 285-370), an RGG domain (residues 371-421), and the Zinc Finger (ZnF) domain (residues 422-453). FUS_269-454_ binds various RNAs robustly.^33^ We conducted 2D ^1^H,^15^N-HSQC experiments in the presence or absence of RNA S1, S2, W1, or C2. Addition of each RNA caused NMR chemical shifts in the RRM, RGG, and ZnF regions of FUS, consistent with RNA binding to all three domains (Figure S3D). Extensive NMR resonance broadening and low peak intensity in the spectra of FUS_269-454_ is observed in the presence of RNA S1 or S2 (Figure S3D). This effect is much more pronounced for RNA S1 and S2 than for RNA W1 and C2, particularly in the resonances of residues 290-360, which map to the RRM (Figure S3D). These observations suggest that FUS complexed with RNA S1 or RNA S2 exchange conformations (i.e., RNA binding/unbinding k_ex_) on the intermediate NMR chemical shift timescale or form higher order complexes. Either of these possibilities is consistent with higher affinities of RNA S1 and S2 for the RNA-binding domains than RNA W1 and C2. Thus, while the overall binding sites between FUS and the RNAs are similar, the affinities for the RNA-binding domains are likely different. Specifically, RNA S1 and S2 engage with higher affinity.

### Weak RNA inhibitors engage the FUS ZnF to antagonize FUS assembly

Since the RNA inhibitors engage multiple FUS domains (Figure S3D), we next investigated how RNA interactions with the ZnF domain might contribute to their ability to antagonize FUS assembly. Thus, we generated a FUS^C428A:C433A:C444A:C447A^ (FUS_4C-A_) mutant, which contains four cysteine to alanine substitutions that disrupt the C4-type Zinc coordination scheme, which enable RNA binding (Figure 3A, D).^36^ FUS_4C-A_ formed fibrils with similar kinetics to FUS (Figure 1B, 3D, E). Strong RNA inhibitors S1-S3 effectively prevented and reversed formation of FUS_4C-A_ fibrils (Figure 3D, S3E). Thus, disrupting the ZnF domain has little effect on the activity of strong RNA inhibitors. Consistent with this result, the binding affinities of strong RNA inhibitors to FUS_4C-A_ were not significantly different from their binding affinities to FUS (RNA S1: *K_D_*∼25nM; RNA S2: *K_D_*∼136nM; RNA S3: *K_D_*∼59nM) (Figure 3A). In striking contrast, weak RNA inhibitors could neither prevent nor reverse FUS_4C-A_ assembly (Figure 3E and S3F). Thus, the ZnF domain may play a critical role in enabling weak RNA inhibitors to antagonize FUS assembly but is less important for strong RNA inhibitors.

### FUS_371X_ is refractory to RNA inhibitors

In addition to the RRM and ZnF regions, our NMR studies revealed that a FUS RGG domain also interacted with various RNA inhibitors (Figure S3D). To assess how these interactions might contribute to RNA inhibitor activity, we created a FUS construct consisting of the N-terminal PrLD and RRM (FUS_371X_). As expected, after deleting the RGG domains that enable rapid FUS assembly,^10,37^ FUS_371X_ formed fibrils much more slowly than FUS, taking up to 24 hours to assemble (Figure S3G). Deletion of the C-terminal RGG domains and ZnF affected binding of RNA inhibitors to varying extents (RNA S1: *K_D_*∼107nM; RNA S2: *K_D_* could not be determined; RNA S3: *K_D_*∼124nM) (Figure 3A). For example, the most pronounced change was for RNA S2, where we could not saturate binding to determine a *K_D_*, though RNA binding still occurred (Figure 3A). The *K_D_* of RNA S1 increased from ∼41nM for FUS to 107nM for FUS_371X_. By contrast, the *K_D_* of RNA S3 for FUS and FUS_371X_ were similar.^27^ Importantly, FUS_371X_ fibrillization could not be inhibited or reversed with RNA S1, S2, or W1 (Figure 3F, G). By contrast, RNA S3 could inhibit FUS_371X_ assembly by ∼50% but was unable to reverse FUS_371X_ fibrillization (Figure 3F, G). These findings suggest that the C-terminal RNA-binding domains of FUS (RGG domains and ZnF) enable strong RNA inhibitors to exert their maximal effects in preventing and reversing FUS fibrillization.

### Weak RNA inhibitor W1 displays greater co-operativity than strong RNA inhibitors S1-S3

Next, we assessed the dose dependence of inhibition of FUS assembly by RNA S1, S2, S3, and W1. As expected, the strong RNAs were more effective inhibitors with half maximal inhibitory concentrations (IC_50_) ranging from ∼3-8µM, whereas the IC_50_ of RNA W1 was ∼65µM (Figure 3H-K). If RNA binding to multiple FUS domains were required to reduce assembly, one would expect to observe cooperativity in RNA inhibition. Indeed, both strong and weak RNA inhibitors exhibited cooperativity with Hill coefficients (h) ranging from ∼-1.3 to ∼-4.1 (Figure 3H-K). Strong RNA inhibitors S1, S2, and S3 had less steep dose-response slopes with h values from ∼-1.3 to ∼-2, whereas the weak RNA inhibitor W1 had a steeper dose-response slope with an h value of ∼-4.1. Thus, weak RNA inhibitor W1 displays greater cooperativity than strong RNA inhibitors S1-S3, which may reflect the requirement for a functional RRM and ZnF domain for RNA W1 to be effective. Overall, our findings suggest that short RNAs must engage multiple RNA-binding domains of FUS for maximal inhibition of assembly.

### The FUS RRM and ZnF domains cooperate to maintain FUS solubility in human cells

Injecting RNase into the nucleus causes FUS to aggregate, indicating that endogenous RNAs promote FUS solubility.^38^ However, whether specific, short RNAs can be introduced as agents to prevent and reverse aberrant phase separation of FUS in cells is unclear. To investigate how the RNA-binding domains of FUS might contribute to FUS solubility in human cells, we established an optogenetic system to control FUS phase separation in response to blue light. Thus, we adapted the Corelet system which has been used to map intracellular phase behavior of the FUS PrLD and other intrinsically disordered regions (IDRs) in response to blue light.^39^ Corelet is a two-module system that relies on the light-based dimerization between an improved light-induced dimer (iLID) domain on a ferritin heavy chain (FTH1) protein core (which forms 24mers) and an SspB domain on the molecule of interest, in this case FUS.^39^ We first generated FUS constructs (amino acids 1-453) containing wild-type RNA-binding regions, mutated RRM (4FL), mutated ZnF domain (4CA), or double RRM and ZnF mutants (4FL/4CA) with C-terminal SspB peptide tags (Figure 4A). We omitted the C-terminal RGG domain and PY-NLS to prevent spontaneous phase separation of full-length FUS proteins with mutated RNA-binding regions,^34^ and also to reduce interaction with the nuclear-import receptor, Karyopherin-β2, which can prevent FUS phase separation.^8,40–43^

**Figure 4.**
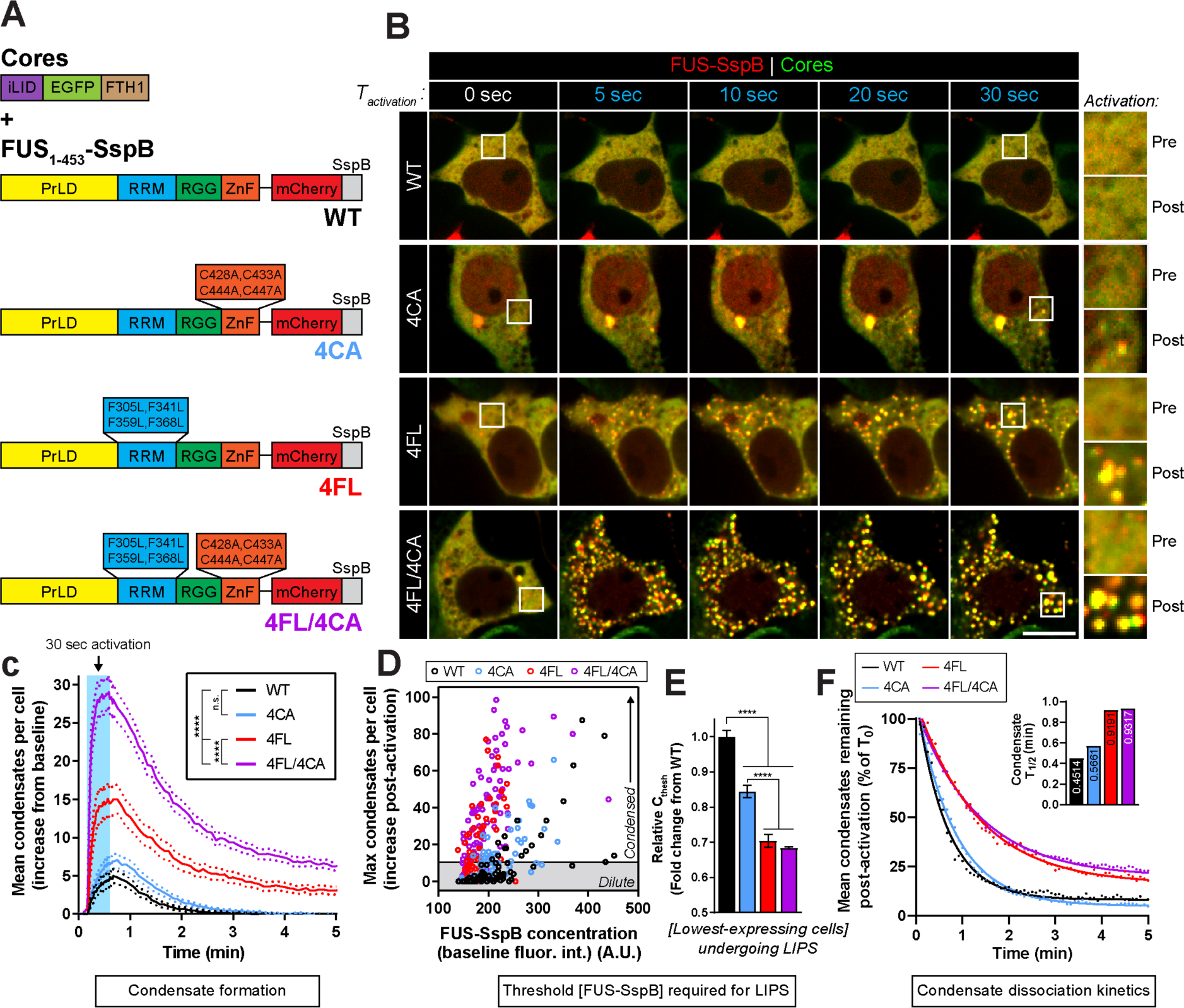
The FUS RRM and ZnF domains cooperate to maintain FUS solubility in human cells. **(A)** Schematic of iLID cores and FUS-SspB mutant constructs used in (B-F). **(B)** Representative images of HEK293 cells co-expressing iLID cores (green) and the indicated mutant FUS-SspB protein (red) prior to and during a 30s light activation protocol (488nm, 75% laser power). Insets show the boxed cytoplasmic area at baseline and following 30s activation. Bar, 10µm. **(C)** Quantification of the average number of FUS-SspB assemblies formed per cell during and following a 30s light activation period. *n*=68-91 cells per condition. Data are shown as mean (solid lines) ± SEM (dashed lines). Two-way ANOVA with Tukey’s post-hoc test was used to compare across groups; ****, *p*<0.0001. **(D)** Graph of maximal light response (number of condensates during the activation period) in (C) plotted against baseline FUS-SspB concentration. Data points represent individual cells. **(E)** Quantification of representative threshold concentrations required for cells to undergo LIPS. *n* = the lowest-expressing 5 cells with >10 condensates post-activation per condition. Values represent means±SEM. Fluorescence intensity values are normalized to WT and shown as fold-change. One-way ANOVA with Tukey’s post-hoc test was used to compare across groups; ****, *p*<0.0001. **(F)** Quantification of FUS condensate dissociation kinetics following conclusion of light activation. Number of condensates per cell were plotted over time as a percentage of condensates in the first frame following light removal (T_0_). One-phase exponential decay curves were fit and T½ was calculated for each condition and plotted in the inset (top right). *n*=20-76 cells per group.

We exposed cells co-expressing FUS-SspB constructs and photo-activatable seeds (iLID-EGFP-FTH1)^39^ to acute (30 second) blue light activation sequences and assessed FUS condensate formation and dissolution (Figure 4B). Interestingly, mutations within the RRM region (4FL, red trace) led to enhanced formation of light-induced FUS-SspB condensates compared to wild-type (WT) FUS_1-453_ (black trace), whereas mutations within the ZnF domain (4CA, blue trace) only mildly increased light-induced phase separation (LIPS) (Figure 4B, C). However, when ZnF mutations were combined with RRM mutations (4FL/4CA, purple trace), a further enhancement of FUS-SspB condensate formation was observed when compared to either RRM or ZnF mutations alone (Figure 4B, C). Thus, the FUS RRM and ZnF cooperate to prevent FUS condensation, with the RRM playing a larger role than the ZnF.

A similar pattern was observed when we next examined LIPS as a function of FUS-SspB expression level (Figure 4D, E). Here, ZnF mutations (4CA) slightly reduced the threshold protein concentration (C_thresh_) required for condensate formation, whereas RRM (4FL) and dual RRM and ZnF (4FL/4CA) mutations greatly reduced C_thresh_ (Figure 4D, E). Following light removal, RRM and dual RRM and ZnF mutations caused decelerated dissolution of light-induced condensates compared to WT and ZnF-only mutants (Figure 4F), which indicates increased stability of these condensates. Together, these results suggest that endogenous RNA contacts with the FUS RRM play a critical role in preventing aberrant phase transitions within the intracellular milieu, but the FUS ZnF domain also contributes.

### An optogenetic model of FUS proteinopathy recapitulates ALS-FUS phenotypes

Next, to determine whether short RNA oligonucleotides can prevent and reverse aberrant FUS condensation in human cells, we developed a light-inducible model of FUS proteinopathy based on a previous model developed to control TDP-43 aggregation.^15^ Specifically, we generated a doxycycline-inducible optogenetic Cry2-FUS (optoFUS) construct to selectively induce FUS proteinopathy under the spatiotemporal control of light stimulation (Figure S4A, B). We utilized Cry2olig as the tag, which is a variant of the Photolyase-Homologous Region (PHR) of the Cryptochrome 2 protein from *Arabidopsis thaliana* that undergoes reversible homo-oligomerization (within ∼5 min) in response to blue light.^44^

We first tested whether Cry2olig-mediated increases in focal intracellular concentrations of optoFUS protein can seed intracellular FUS proteinopathy upon chronic light exposure. Human cells treated with 10ng/mL doxycycline to express optoFUS protein were exposed to 8 hours of blue light (∼0.1-0.3mW/cm^2^, 465nm) or darkness, and were then examined by immunofluorescence (Figure S4B). Interestingly, cells expressing optoFUS that were exposed to blue light exhibited a significant depletion of nuclear optoFUS signal and enhanced formation of cytoplasmic inclusions relative to optoFUS-expressing cells kept in the dark (Figure S4A-E). Fluorescence recovery after photo-bleaching (FRAP) analysis of light-induced, cytoplasmic optoFUS inclusions revealed minimal recovery after photo-bleaching, indicating that optoFUS inclusions had solid-like properties with limited dynamics indicative of an aberrant phase transition (Figure S4F). Sedimentation analysis confirmed that light-induced optoFUS inclusions were detergent-insoluble and increased the amount of insoluble endogenous FUS relative to cells kept in darkness (Figure S4G). Thus, our optoFUS system recapitulates the cytoplasmic aggregation and nuclear depletion of FUS observed in ALS-FUS or FTD-FUS.

To determine whether optoFUS inclusions more closely resembled ALS-FUS or FTD-FUS pathology observed in postmortem patient tissues, we performed immunofluorescence analysis to assess common pathological hallmarks of ALS-FUS or FTD-FUS. OptoFUS inclusions did not colocalize with FET proteins EWSR1 and TAF15 (Figure S4H), two RBPs with PrLDs that typically co-deposit with FUS inclusions in FTD but not in ALS patients.^45^ Moreover, optoFUS inclusions were recognized by the 9G6 monoclonal antibody that recognizes methylated FUS, which is more consistent with ALS-FUS pathology (Figure S4I).^46–48^ In addition, optoFUS inclusions do not co-localize with stress granule marker G3BP1 or TDP-43 (Figure S4J, K). This immunocytochemical profile was also observed when optoFUS inclusions were induced in human ReNcell VM neurons (Figure S4L, M), indicating consistent results across human cell and neuronal models. Thus, light-activated optoFUS inclusions exhibit the hallmarks of FUS pathology observed in ALS.

### RNA S1 prevents aberrant phase transitions of FUS in human cells

We next tested whether the strong and weak RNA inhibitors isolated *in vitro* can prevent the formation of intracellular FUS inclusions in the optoFUS system (Figure 5A). We found that 5’-fluorescein-labeled RNA S1 accumulated predominantly in the cytoplasm of human cells ∼2 hours after transfection (Figure S5A-C). Importantly, introducing RNA S1 did not alter the nuclear localization of endogenous FUS (Figure S5D, E). Next, we pre-treated optoFUS-expressing human cells for two hours with strong RNA inhibitors RNA S1 or RNA S2, which can prevent and reverse FUS fibrillization *in vitro*, weak inhibitor RNA W1, or RNA C2, which is ineffective *in vitro* (Figure 5A). After the two-hour pretreatment, blue light was applied, and we monitored the formation of optoFUS inclusions. OptoFUS formed abundant cytoplasmic inclusions in cells treated with RNA C2 (Figure 5B, C, S5F). Likewise, RNA W1 was ineffective in preventing optoFUS inclusion formation (Figure S5F). Interestingly, despite being effective *in vitro*, RNA S2 only slightly prevented optoFUS inclusion formation in human cells (Figure S5F). Remarkably, however, pre-treatment with RNA S1 resulted in a dose-dependent reduction in optoFUS inclusion formation when compared to treatment with RNA C2 (Figure 5B, C). Thus, RNA S1, which is similar in length to therapeutic ASOs,^22^ can prevent aberrant phase transitions of FUS *in vitro* and in human cells.

**Figure 5.**
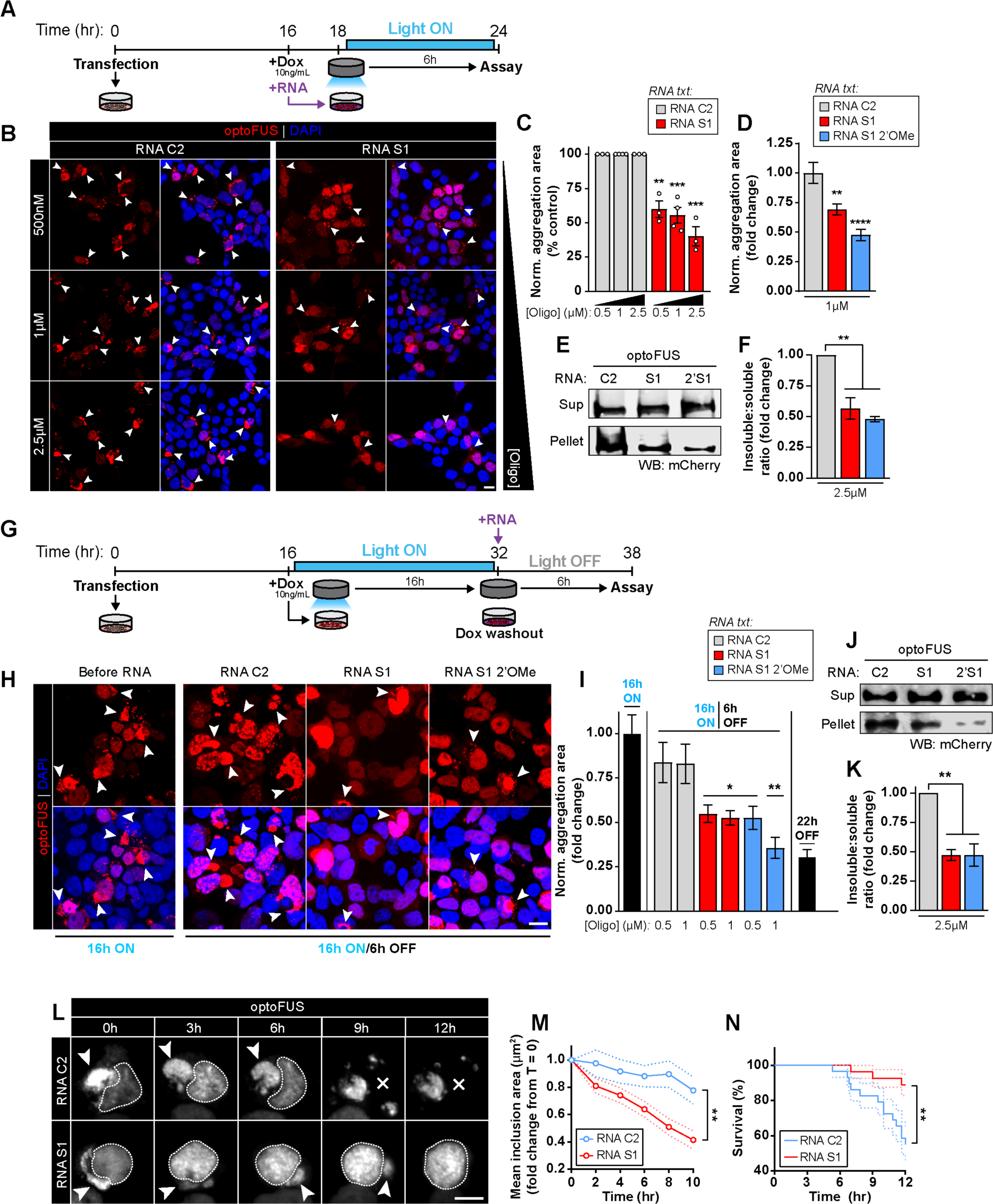
RNA S1 prevents and reverses aberrant phase transitions of FUS in human cells. **(A)** Schematic of light-activation paradigm used to assess whether RNA inhibitors can prevent optoFUS phase separation used in (B-F). **(B)** Representative images of optoFUS-expressing HEK293 cells pre-treated with control RNA C2 (Ctrl) or strong RNA inhibitor (S1) at concentrations ranging from 500nM-2.5µM for 2h prior to exposure to 6h of light activation. Bar, 10µm. Arrows indicate cytoplasmic optoFUS assemblies. **(C)** OptoFUS aggregation area in light-activated cells pre-treated with RNA C2 (Ctrl) or RNA S1 at the indicated concentrations. Data points represent individual experiments, *n*=3-4 individual experiments, 620-904 cells across 9 randomized fields-of-view per experiment. Values are normalized to control treatments within each treatment concentration group and presented as percentage of control per experiment. Values are means±SEM (n=3-4). Unpaired Student’s t-tests were used to compare RNA C2 and S1 conditions within each treatment concentration. **, *p*<0.01, ***, *p*<0.001. **(D)** OptoFUS aggregation area in light-activated cells pre-treated with 1µM RNA C2 (Ctrl), RNA S1, or 2’OMe-modified RNA S1 oligonucleotide. *n* = 9 randomized fields-of-view, 144-323 cells per field. Values are normalized to control treatments and presented as fold-change from control. Values are means±SEM (n=3 independent experiments). One-way ANOVA with Tukey’s post hoc test was used to compare across groups. **, *p*<0.01, ****, *p*<0.0001. **(E)** Detergent-solubility fractionation of cells pre-treated with 2.5µM RNA C2 (Ctrl), RNA S1, or 2’OMe-modifed RNA S1 prior to light-activation. **(F)** Quantification of ratios of detergent-insoluble to detergent-soluble band intensities in each treatment group described in (E). *n*=3 biological replicates per condition. Values are means±SEM (n=3 independent experiments). One-way ANOVA with Tukey’s post hoc test was used to compare across groups; **, *p*<0.01. **(G)** Schematic of light-activation paradigm used to assess whether RNA inhibitors can reverse optoFUS phase separation used in (H-K). **(H)** Representative images of optoFUS-expressing HEK293 cells exposed to the light-induction protocol outlined in (G) before addition of RNA (left panel) and following a 6h treatment with 1µM RNA C2 (Ctrl), RNA S1 or 2’OMe-modified RNA S1 (right panels) in the absence of further light stimulation. Bar, 10µm. Arrows indicate cytoplasmic optoFUS assemblies. **(I)** optoFUS aggregation area prior to (left black bar) and following treatment (middle bars) with the indicated RNA. Aggregation values from cells kept in darkness throughout the experiment (22h OFF, black bar) are included for reference. Values are normalized to groups fixed immediately following light activation and prior to RNA treatment. *n*=9 randomized fields-of-view, 79-275 cells per field. Comparisons shown are between control and targeting RNA treatments. Values are means±SEM (n=3 independent experiments). One-way ANOVA with Tukey’s post hoc test was used to compare across groups. *, *p*<0.05, **, *p*<0.01. **(J)** Detergent-solubility fractionation of cells treated with 2.5µM of the indicated RNA for 6h in the absence of light following pre-formation of light-induced optoFUS aggregates as in (G). **(K)** Quantification of ratios of detergent-insoluble to detergent-soluble band intensities in each treatment group described in (J). *n*=3 biological replicates per condition. Data shown are means±SEM. One-way ANOVA with Tukey’s post hoc test was used to compare across groups; **, *p*<0.01. **(L)** Representative live images of HEK293 cells expressing optoFUS pre-exposed to 10h of blue light stimulation following 2µM treatment with the indicated oligonucleotides as in (H-K). Arrows indicate inclusions and X indicates cell death. Cell nuclei are circled. Bar, 10µm. **(M)** Quantification of mean optoFUS inclusion size over time following treatment with the indicated oligonucleotides. *n* = 26-29 inclusions per treatment. Data are presented as mean (solid lines) ± SEM (dashed lines). Two-way mixed design ANOVA with Sidak’s correction was used to compare across groups; **, *p*<0.01. **(N)** Survival curves of cells containing optoFUS inclusions at the onset of imaging treated with the indicated oligonucleotides. *n*=27-29 cells per treatment. Kaplan-Meier estimates were used to generate survival curves (dashed lines represent standard error) and Gehan-Breslow-Wilcoxon tests were used to compare across groups, **, *p*<0.01. See also **Figure S4** and **S5**.

RNA oligos can be quickly digested by ribonucleases in cells. Thus, we also designed RNA analogues with greater intracellular stability to test in our optoFUS model. Using RNA S1 as a template, we designed RNA analogues with 2’OMe modifications to test both *in vitro* and in cells (Table S1). 2’OMe-modified RNA S1 and unmodified RNA S1 exhibited similar ability to prevent and reverse FUS fibrillization *in vitro* (Figure S5G, H). Importantly, 2’OMe-modified RNA S1 exhibited slightly enhanced inhibition of optoFUS inclusion formation compared to unmodified RNA S1 (Figure 5D-F). Thus, 2’OMe-modifications of RNA S1 could help stabilize the oligo in cells without impairing its ability to antagonize FUS aggregation.

### RNA S1 reverses aberrant phase transitions of FUS in human cells

We next determined whether treatment with RNA inhibitors could reverse formation of preformed optoFUS inclusions. Thus, optoFUS-expressing cells were first subjected to chronic light stimulation to induce optoFUS aggregates prior to RNA treatment and doxycycline washout to eliminate further optoFUS expression during a 6-hour dark “disassembly” period (Figure 5G). The control RNA C2 had little effect on preformed optoFUS inclusions (Figure 5H). Remarkably, RNA S1 and 2’OMe-modified RNA S1 oligonucleotides significantly reduced optoFUS inclusion burden toward levels observed in optoFUS-expressing cells kept in darkness throughout the experiment (Figure 5H-I). This effect was confirmed by sedimentation analysis of optoFUS cell lysates collected following the same light induction and treatment paradigm (Figure 5J-K). Thus, RNA S1 and 2’OMe-modified RNA S1 can reverse aberrant FUS phase transitions in human cells.

Next, we explored the kinetics of optoFUS inclusion dissolution by RNA S1 (Figure 5G). Remarkably, RNA S1 reduced cytoplasmic optoFUS inclusion size within 2-3h of treatment, whereas RNA C2 had no effect (Figure 5L, M). Indeed, cells treated with RNA C2 displayed persistent cytoplasmic optoFUS inclusions and exhibited reduced survival after ∼6-12h (Figure 5L, N). By contrast, cells treated with RNA S1 cleared cytoplasmic optoFUS inclusions and restored nuclear FUS, which was accompanied by increased survival (Figure 5L, N). Thus, RNA S1 clears cytoplasmic FUS inclusions, restores nuclear FUS, and mitigates toxicity.

### RNA S1 prevents FUS phase separation and mitigates toxicity in iPSC-derived FUS^R521G^ motor neurons

Next, we tested whether RNA S1 can prevent and reverse aberrant FUS aggregation in human motor neurons. Thus, we employed iPSC-derived motor neurons (iMNs) harboring ALS-linked FUS^R521G^ (Figure 6A). RNA S1 effectively prevents and reverses FUS^R521G^ fibrillization (Figure S2E, I). Upon differentiation, iMNs harboring ALS-linked FUS^R521G^ exhibited increased FUS mislocalization to the cytoplasm, compared to control iMNs harboring WT FUS (Figure 6B, C). FUS^R521G^-iMNs treated with RNA S1 showed partial restoration of FUS^R521G^ nuclear localization, whereas control C2 RNA had no effect (Figure 6B, C). Moreover, upon sodium arsenite treatment, FUS^R521G^ iMNs exhibited formation of FUS-positive stress granules, whereas FUS-positive stress granules were less abundant in control iMNs (Figure 6D, E). Importantly, FUS^R521G^ iMNs treated with RNA S1 but not RNA C2 exhibited a reduction in FUS-positive stress granules, indicating that RNA S1 prevented FUS recruitment into these phase-separated structures (Figure 6E). Moreover, RNA S1 reduced stress granule number and area in FUS^R521G^ iMNs, indicating that FUS^R521G^ was likely driving stress granule assembly (Figure 6F, G). Treatment of control or FUS^R521G^ iMNs with a proteotoxic stressor, tunicamycin, reduced iMN viability (Figure 6H, I).^49^ Remarkably, RNA S1 but not RNA C2 mitigated toxicity in FUS^R521G^ iMNs, but not control iMNs (Figure 6H, I). Thus, RNA S1 is neuroprotective under proteotoxic conditions in iMNs expressing ALS-linked FUS^R521G^.

**Figure 6.**
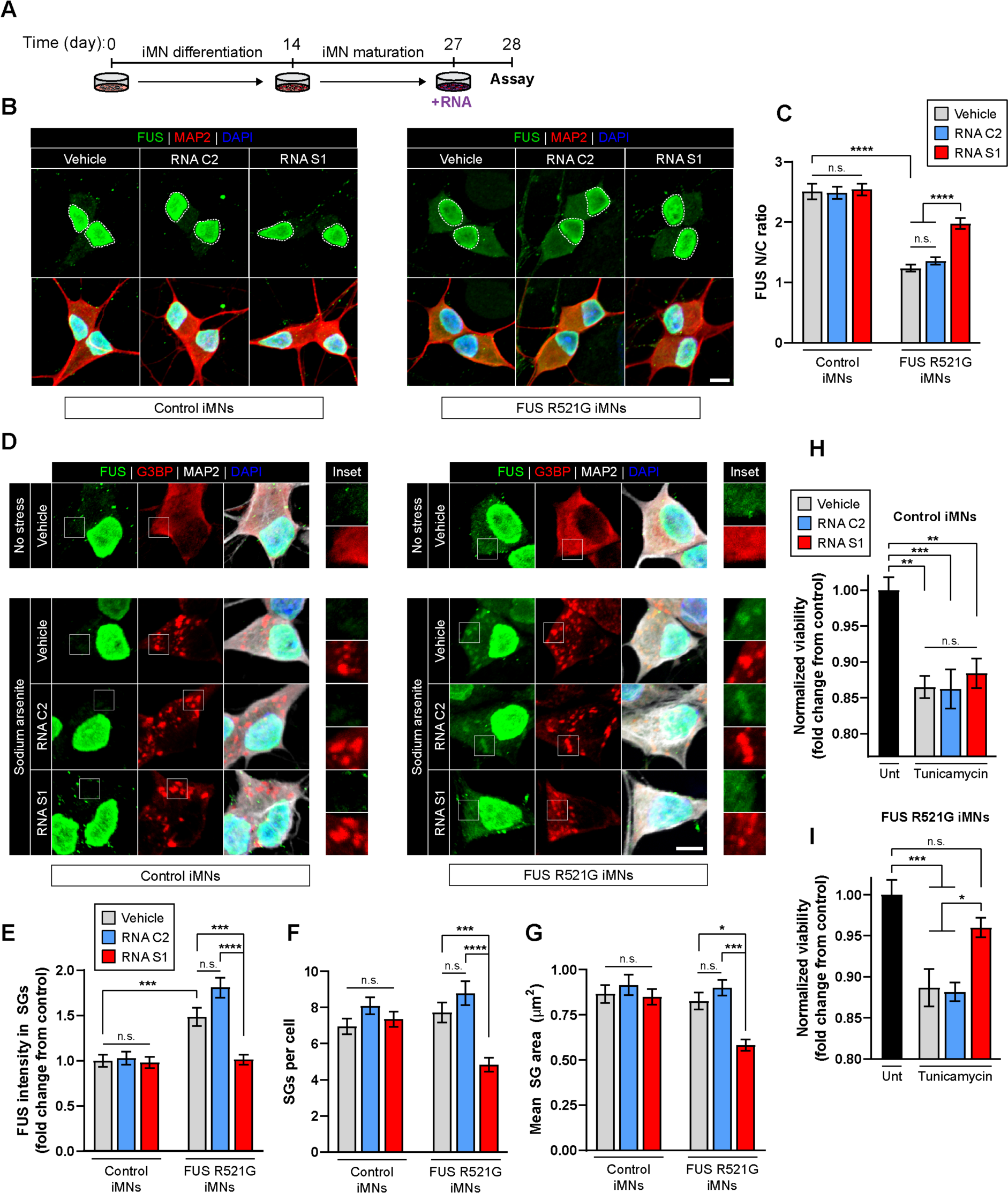
RNA S1 prevents FUS phase separation and mitigates toxicity in iPSC-derived FUSR^521G^ motor neurons. **(A)** Schematic of motor neuron differentiation and RNA oligonucleotide treatment paradigm used in (B-I). Control iPSC motor neurons (iMNs) or FUSR^521G^ ALS iMNs that harbor a single FUS^R521G^ mutation in the NLS were treated with a control or RNA S1 (500nM for 24h). **(B)** Representative images of immunostained control and FUS^R521G^ iMNs revealed enriched cytoplasmic FUS protein in FUS^R521G^ iMNs. **(C)** Graph depicts means±SEM of FUS nuclear/cytoplasmic (N/C) ratio in FUS^R521G^ ALS iMNs, indicating a reduced FUS N/C localization in vehicle and control oligonucleotide (RNA C2) treated iMNs compared to controls. RNA S1 enhanced FUS nuclear localization in FUS^R521G^ ALS iMNs but not control iMNs (n=81-87 iMNs over 3 differentiations; two-way ANOVA with Bonferroni correction: ****, p<0.0001). **(D)** Representative images of immunostained SGs (G3BP1; inset) induced with NaAsO_2_ (0.5mM for 45min) in control and FUS^R521G^ iMNs pre-treated with a vehicle, control oligonucleotide (RNA C2) or RNA S1 (500nM). **(E)** FUS intensity (pixel/µm^2^) colocalization with G3BP1 SGs was enhanced in FUS^R521G^ iMNs compared to controls and reduced by RNA S1 (means±SEM, n=363-509 SGs, over 3 differentiations; two-way ANOVA with Bonferroni correction: ***, p<0.001; ****, p<0.0001). **(F)** G3PB1+ SG number/cell and **(G)** mean G3PB1+ SG area were reduced in FUS^R521G^ iMNs upon S1 oligonucleotide treatment but did not affect control iMNs (means±SEM, n=363-509 SGs, 59-67 iMNs over 3 differentiations; two-way ANOVA with Bonferroni correction: ***, p<0.001; ****, p<0.0001). **(H-I)** Control and FUS^R521G^ iMNs exhibit reduced viability (as measured by intracellular ATP) following tunicamycin treatment (25µM for 24h) when compared to untreated iMNs (Utr). RNA S1 (500nM) treatment enhanced FUS^R521G^ iMN viability compared to vehicle and control oligonucleotide (RNA C2) treatment. (4 technical replicates per experiment, 3 differentiations per line; one-way ANOVA with Bonferroni correction: *, p<0.05; **, p<0.01; ***, p<0.001).

### A short, specific RNA, Clip34, directly prevents and reverses aberrant TDP-43 phase separation

In addition to FUS, cytoplasmic aggregation of other RBPs with PrLDs has been reported in patient postmortem tissue in ALS/FTD and related disorders, including TAF15, EWSR1,^45,50,51^ hnRNPA1, hnRNPA2,^52–54^ and TDP-43.^3,4,7^ Previously, we established that Clip34, a 34nt RNA derived from the 3’UTR of the *TARDBP* gene (Table S2), which binds to TDP-43,^13,55–58^ can prevent aberrant phase transitions of TDP-43 in optogenetic neuronal models and mitigate associated neurotoxicity.^15^ At physiological concentrations and buffer conditions, purified TDP-43 spontaneously phase separates.^13,59^ We now establish that Clip34 prevents (IC_50_∼0.31µM) and reverses (EC_50_∼0.6µM) TDP-43 PS directly in a dose-dependent manner, whereas a control RNA oligo, (AC)_17_, which does not bind TDP-43 has no effect on TDP-43 PS (Figure S6A-F). The ability of Clip34 to prevent and reverse TDP-43 PS required interaction with the TDP-43 RRMs, as PS by the TDP-43 mutant, TDP-43^5FL^, which bears F147L, F149L, F194L, F229L, and F231L mutations in the RRMs that impair RNA binding,^25^ was unaffected by Clip34 (Figure S6A-F).

Purified TDP-43 can also rapidly assemble in fibrillar structures.^8,9,12,15^ Importantly, Clip34 also prevented TDP-43 fibrillization (IC_50_∼0.62µM), whereas (AC)_17_ was ineffective (Figure S6G-J). By contrast, Clip34 was unable to prevent or reverse TDP-43^5FL^ fibrillization (Figure S6K-M). Remarkably, Clip34 but not (AC)_17_, could partially reverse aggregation of TDP-43 (Figure 7A-C). Thus, Clip34 engages the TDP-43 RRMs to prevent *and* reverse TDP-43 PS and aggregation. Our findings suggest that short, specific RNAs might be broadly applicable to antagonize aberrant phase transitions of disease-linked RBPs with PrLDs.

**Figure 7.**
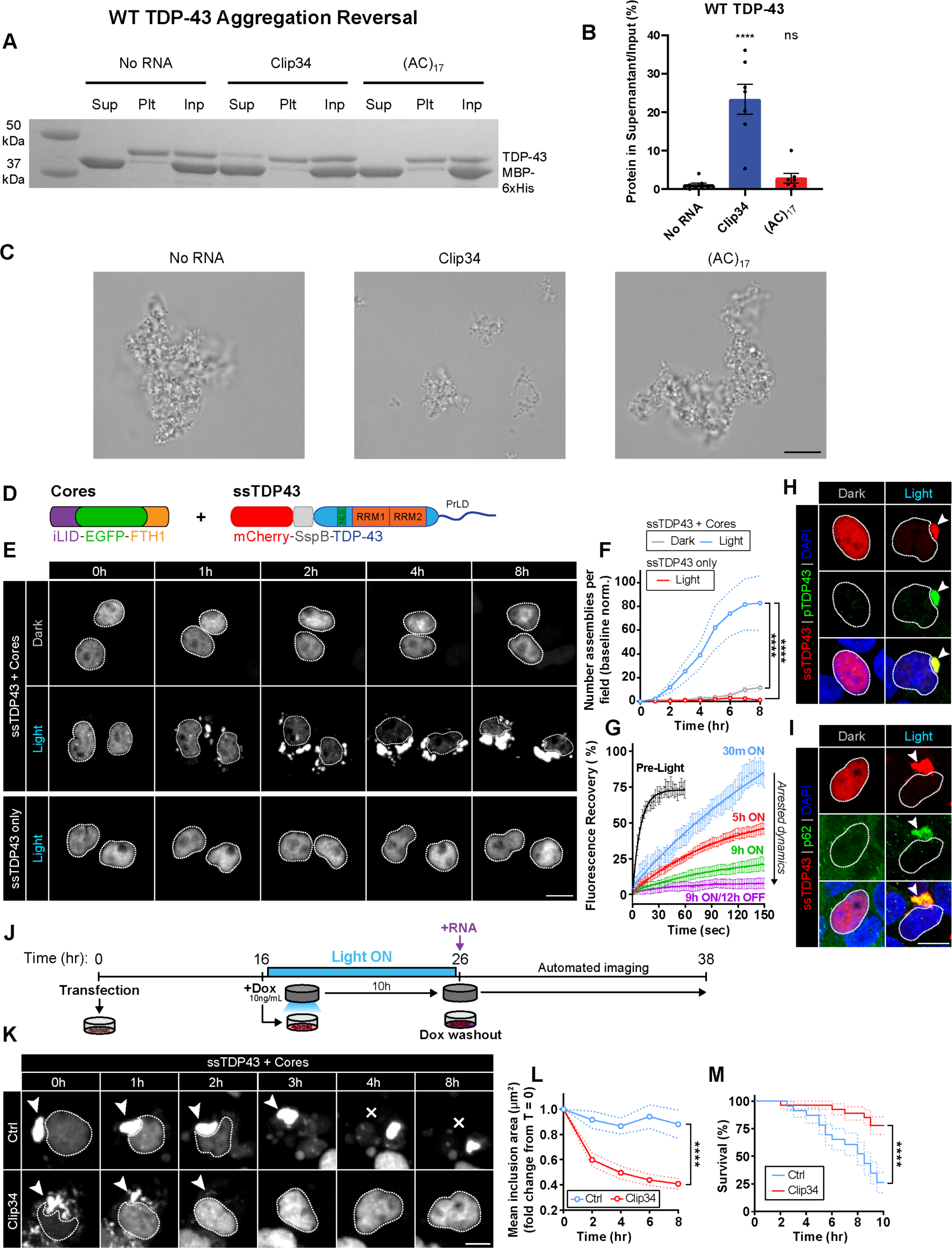
Clip34 reverses aberrant TDP-43 phase separation. **(A-C)** Preformed TDP-43 aggregates (4µM) were incubated with buffer, Clip34, or (AC)_17_ (40µM) for 16h. Reactions were processed for sedimentation analysis and the supernatant, pellet, and input (100%) were fractionated by SDS-PAGE and Coomassie stain (A). The fraction of soluble TDP-43 in the supernatant was determined by densitometry (B). Values represent means±SEM (n=7). One-way ANOVA comparing to the No RNA condition; Dunnett’s multiple comparisons test; ns: *p*>0.05, \*\*\*\**p* adjusted ≤0.0001. Alternatively (C), reactions were viewed by brightfield microscopy. Note that large and dense TDP-43 aggregates persist in buffer or after treatment with (AC)_17_ RNA, whereas Clip34 reduces aggregate size. Bar, 10µm. **(D)** Schematic of iLID cores and ssTDP-43 constructs used in (E-M). **(E)** Representative images of HEK293 cells co-expressing iLID cores and ssTDP43 (top panels) or ssTDP43 alone (bottom panel) during simultaneous live imaging and light stimulation. Bar, 10µm. Cell nuclei are circled. **(F)** Quantification of number of ssTDP43 assemblies per field-of-view during time course of live imaging. *n*=6 fields-of-view, 171-408 cells per field. Data are shown as means (solid lines) ± SEM (dashed lines). *n*=3 experiments. Two-way ANOVA with Tukey’s post hoc test was used to compare across groups; ****, *p*<0.0001. **(G)** FRAP analysis of ssTDP43 assemblies formed in HEK293 cells co-expressing iLID cores over increasing lengths of blue light stimulation (0.1-0.3mW/cm^2^, 465 nm). Values are means±SEM (n=3 experiments). **(H, I)** Immunofluorescence analysis of co-localization between ssTDP43 inclusions formed following 8h of blue light stimulation and the pathological hallmarks phospho-TDP43 (H) and p62 (I). Cell nuclei are circled. Arrows indicate light-induced inclusions. Bars, 10µm. **(J)** Schematic of light-activation paradigm used for pre-formation of ssTDP43 inclusions prior to RNA treatments and live imaging used in (L-M). **(K)** Representative live images of HEK293 cells co-expressing iLID cores and ssTDP43 pre-exposed to 10h of blue light stimulation following treatment with 2µM of RNA C2 (Ctrl) or RNA Clip34. Arrows indicate inclusions and X indicates cell death. Cell nuclei are circled. Bar, 10µm. **(L)** Quantification of mean inclusion size over time following treatment with RNA C2 (Ctrl) or RNA Clip34. Values shown are normalized to areas of individual inclusions at the onset of imaging and are presented as fold-change from T_0_. *n*=25-37 inclusions per treatment. Data are shown as means (solid lines) ± SEM (dashed lines). Two-way mixed design ANOVA with Sidak’s correction; ****, *p*<0.0001. **(M)** Survival curves of cells containing ssTDP43 inclusions at the onset of imaging treated with the indicated oligonucleotides. *n*=23-26 cells. Kaplan-Meier estimates were used to generate survival curves (dashed lines represent standard error) and Gehan-Breslow-Wilcoxon tests were used to compare across groups, ****p≤0.0001. See also **Figure S6** and **S7**.

### A short, specific RNA, Clip34, reverses aberrant TDP-43 phase separation in human cells

We next investigated whether Clip34 could reverse aberrant TDP-43 phase separation in human cells. Thus, we developed a new optogenetic model of full-length TDP-43 aggregation based upon the Corelet system.^39^ Cytoplasmic iLID-FTH1 cores were co-expressed with full-length TDP-43 that was N-terminally tagged with mCherry-SspB (ssTDP43) under the control of the doxycycline-inducible Tet3G promoter (Figure 7D). Human (HEK293) cells expressing these constructs were exposed to 10ng/mL doxycycline treatment and chronic blue light activation (∼0.3-1mW/cm^2^, 465nm) or darkness for 8h (Figure 7E, F) to induce TDP-43 condensation. Automated light activation and live-cell imaging (Figure S7A-F) revealed significant accumulation of ssTDP43 condensates in cells exposed to chronic blue light but not cells kept in the darkness (Figure 7E, F). Importantly, cells expressing ssTDP43 alone (without iLID cores) exposed to the same light activation conditions did not form ssTDP43 condensates (Figure 7E, F). Thus, ssTDP-43 condensates form due to a specific effect of light-induced Corelet association rather than by a non-specific effect of blue light exposure (Figure 7E, F).

We next determined the effect of increased light exposure on TDP-43 dynamics within the induced ssTDP43 condensates. FRAP analysis was performed on human cells expressing iLID cores and ssTDP43 both before light exposure and on ssTDP43 assemblies in response to increasing lengths of blue light activation (∼0.1-0.3mW/cm^2^, 465nm) (Figure 7G). Initial assemblies of ssTDP43 formed in response to 30 minutes of blue light displayed nearly full fluorescence recovery following bleaching, suggesting a dynamic or liquid-like state of these condensates (Figure 7G). However, a progressive decrease in recovery was observed of condensates exposed to increasing lengths of blue light activation, indicating arrested dynamics of these structures over time that remain stable for at least 12 hours following light removal (Figure 7G). Thus, much like *in vitro* reactions,^59–61^ light-induced ssTDP43 aggregate formation in a cellular context begins with an initial liquid-like stage followed by maturation of these condensates into solid-phase inclusions over time. Furthermore, the aberrant, solid TDP-43 assemblies formed after chronic (8 hour) blue light activation bore the pathological hallmarks of hyperphosphorylation (Figure 7H) and colocalization with p62 (Figure 7I). These phenotypes are commonly observed with TDP-43 inclusions in ALS/FTD postmortem patient tissue.^4^

Next, we tested whether Clip34 affects endogenous TDP-43 localization or function. Ideally, Clip34 would not perturb endogenous TDP-43 localization or splicing activity. Indeed, Clip34 treatment did not change the nuclear localization of endogenous TDP-43 (Figure S7G, H). Moreover, using a CFTR minigene assay to assess TDP-43 splicing activity, we found that the splicing function of TDP-43 was not affected by Clip34 treatment (Figure S7I-K). Thus, Clip34 does not affect endogenous TDP-43 localization or splicing activity.

To test whether Clip34 could reverse TDP-43 aggregation within human cells, we induced the formation of ssTDP43 inclusions with 10 hours of chronic blue light activation (∼0.1-0.3mW/cm^2^, 465nm) (Figure 7J). Doxycycline was then washed out to switch off ssTDP-43 expression, and cells were treated with control RNA C2 or Clip34 and imaged for 10 hours (Figure 7J). Remarkably, treatment with Clip34 resulted in a significant decrease in TDP-43 inclusion size over time when compared to control RNA C2-treated cells along with a restoration of nuclear TDP-43 (Figure 7K, L). Indeed, TDP-43 inclusions were cleared, and TDP-43 was restored to the nucleus (Figure 7K, L). Importantly, Clip34 significantly extended survival in cells containing TDP43 inclusions at the onset of imaging (Figure 7M). Thus, preformed TDP-43 and FUS inclusions can be reversed by short, specific RNAs in human cells to mitigate toxicity. Since short RNA oligonucleotides can be effectively delivered to the human brain, these agents could have therapeutic utility for ALS/FTD and related disorders.

## Discussion

An important innovation for ALS/FTD treatment will be the advent of deliverable therapeutic agents that reverse the aberrant cytoplasmic aggregation of TDP-43 and FUS, and return these proteins to native form and nuclear function.^1^ These agents would be able to counter any toxic gain of function of cytoplasmic aggregated TDP-43 or FUS conformers, as well as any toxic loss of TDP-43 or FUS function.^1^ Here, we have identified short RNAs (25-34 nts) that can prevent and, remarkably, reverse aberrant phase transitions of FUS and TDP-43 *in vitro* and in human cells. Short RNA oligonucleotides of this length can be readily delivered to the CNS.^21^ Hence, these agents could have therapeutic utility for ALS/FTD and related disorders.

Our most potent RNA for FUS is RNA S1 (Table S2), a 25mer containing GGUG and GGU FUS-binding motifs, which is derived from the 3’UTR of the *BDNF* gene.^26^ RNA S1 directly prevents and reverses condensation and fibrillization of purified FUS and ALS-linked FUS variants. RNA S1 engages the RRM to prevent and reverse FUS fibrillization. Accordingly, mutating the FUS RRM to an RNA-binding deficient form induced cytoplasmic FUS aggregation in human cells in our optogenetic model. However, RNA S1 binding to the RRM is not sufficient as RNA S1 was unable to antagonize fibrillization of FUS_371X_, which harbors the RRM but lacks the C-terminal RGG domains and ZnF. Since RNA S1 could prevent and reverse fibrillization of a FUS ZnF mutant, FUS_4C-A_, these findings suggest that RNA S1 must engage the FUS RRM and RGG regions to antagonize FUS fibrillization. Indeed, NMR revealed that RNA S1 can engage the FUS RRM and a RGG domain tightly. These binding events likely elicit a conformational change in FUS, which promotes FUS solubilization regardless of whether FUS is trapped in a liquid condensate or a solid fibril. This hypothesis is supported by our smFRET observations where another strong RNA inhibitor, RNA S2, locks FUS in a conformation that is averse to the dynamic multimerization. We suggest that these short RNAs enforce a FUS conformation that limits the multivalency that underpins PS and fibrillization.

We found several short RNAs (S1-S8; Table S2) that strongly inhibited FUS PS and fibrillization. However, RNAs S1 and S2 were unusual in their ability to prevent and reverse FUS PS and fibrillization. Moreover, not any FUS-binding RNA can antagonize FUS PS and fibrillization. We uncovered several short RNAs that engage FUS (e.g., W1; Table S2) that allow FUS PS but reduce FUS fibrillization. We also found several short RNAs (N1-N6) that had no effect on FUS PS and fibrillization. Overall, our findings suggest that RNA sequence, length, and structure encode the ability to prevent and reverse FUS PS and fibrillization. Effective RNAs engage multiple RNA-binding domains of FUS to elicit these effects.

Even though RNA S1, S2, and W1 can prevent FUS fibrillization at the pure protein level, only RNA S1 was effective in human cells and motor neurons at antagonizing aberrant FUS assembly and toxicity. We employed unmodified forms of these RNAs, which may limit their stability in cells. Nonetheless, RNA S1 was effective in cells as an unmodified RNA and was also effective *in vitro* and in cells as a 2’OMe-modified version to increase stability in cells. It will be important to determine the precise features of short RNAs that enable activity in a neuronal context. Importantly, RNA S1 prevented and reversed the formation of aberrant cytoplasmic FUS condensates in optogenetic models of FUS proteinopathy. Here, RNA S1 also promoted nuclear localization of FUS. Moreover, RNA S1 prevented cytoplasmic FUS phase separation, promoted nuclear FUS localization, and mitigated proteotoxicity in human iPSC-derived FUS^R521G^ motor neurons.

Our lead RNA for TDP-43 is Clip34 (Table S2), a 34mer that is derived from the 3’UTR of the *TARDBP* gene. We establish that Clip34 can effectively and directly prevent and reverse TDP-43 PS, even at substoichiometric concentrations. Clip34 can also effectively and directly prevent aggregation of purified TDP-43 and can even partially solubilize preformed TDP-43 aggregates. Not any short RNA can exert these effects, which requires Clip34 to specifically engage the TDP-43 RRMs. Importantly, in an optogenetic model of TDP-43 proteinopathy in human cells, Clip34 dissolves aberrant cytoplasmic TDP-43 condensates, restores nuclear TDP-43, and mitigates TDP-43 proteotoxicity.

It is interesting to note that our lead RNAs for FUS and TDP-43 emerge from 3’UTR sequences.^26,55,62^ This finding might indicate an unusual ability of specific 3’UTR sequences to influence aberrant phase separation of RBPs with PrLDs. Moreover, it appears that TDP-43 and FUS inclusions may be susceptible to dissolution by specific short RNAs, which raises the possibility that cells may even regulate TDP-43 or FUS assembly in this way. Indeed, amyloid-like forms of TDP-43 are utilized for beneficial purposes as in myogranules that promote skeletal muscle development and regeneration.^63^ It may be possible for cells to harness these stable TDP-43 structures if mechanisms are readily available to promote their dissolution, which could include specific short RNAs and nuclear-import receptors (NIRs).^8,19^

A possible concern with employing short RNAs in this way is that they might remain too stably bound to TDP-43 or FUS and thus interfere with essential RNA-processing reactions. However, we find that RNA S1 and Clip34 are not toxic to human cells in culture and do not affect the endogenous nuclear localization of FUS or TDP-43. Moreover, these short RNAs localize primarily to the cytoplasm where they would not interfere with nuclear functions of FUS and TDP-43. Indeed, Clip34 does not affect the ability of TDP-43 to function in specific pre-mRNA splicing reactions. In ALS/FTD, cytoplasmic TDP-43 inclusions are relatively devoid of RNA,^15^ which could render aggregated conformers more susceptible to targeting with short RNAs. Furthermore, once FUS or TDP-43 are solubilized by the short RNA they would then engage their cognate NIR for transport to the nucleus. When NIRs engage their RBP cargo they cause the RBP to release any RNA, such that an apo form of the RBP is transported back to the nucleus.^40,42^ Thus, the short RNA would be recycled for further rounds of RBP disaggregation in the cytoplasm and would not affect nuclear RBP function.

We suggest that these short RNAs are attractive therapeutic candidates for further development since they could mitigate gain of toxic function and loss of function toxicity in ALS/FTD connected with TDP-43 or FUS proteinopathy. Indeed, it will be of great interest to assess whether these short RNAs can mitigate neurodegeneration in mouse models of TDP-43 and FUS proteinopathy. Moreover, oligonucleotides of this size can be effectively delivered to the CNS of patients as with several therapeutic ASOs.^21,22,64^ ASOs are also being pursued against FUS, ataxin 2, and TDP-43 as potential therapeutics for ALS/FTD with promising results in model systems and progression to clinical trials.^65–67^ Nonetheless, this strategy runs the risk of promoting loss of function toxicity due to knockdown of these targets, which may be particularly problematic for TDP-43.^68^ By contrast, our short RNAs would restore RBPs to native structure and function thereby eliminating toxicity due to gain and loss of function, which could yield more powerful therapeutic effects. Our strategy could be applied broadly to other RBPs with PrLDs, including hnRNPA1, hnRNPA2, TAF15, and EWSR1, which also accumulate in cytoplasmic aggregates in ALS/FTD and related degenerative disorders,^30^ as well as other RBPs with intrinsically-disordered regions, such as tau which forms cytoplasmic fibrils in various tauopathies, including Alzheimer’s disease.^69^

## Acknowledgments

We thank Edward Barbieri, Linamarie Miller, and Miriam Linsenmeier for critical reading of the manuscript, and Frederic Allain and Fionna Loughlin for NMR assignments. L. G. was supported by Dr. Ralph and Marian Falk Medical Research Trust, Frick Foundation for ALS Research, and NIH grants R35GM138109 and RF1NS121143. J.R.M. was supported by a fellowship from the Center for Protein Conformational Diseases at the University of Pittsburgh. A.M.G. was supported by the Milton Safenowitz Postdoctoral Fellowship from the ALS Association and NIH grant T32NS086749. K.E.C. is supported by NIH grants T32GM132039 and F31NS129101. H.MO. is supported by an AstraZeneca Post-Doctoral Fellowship, an Alzheimer’s Association Research Fellowship, and the Johnson Foundation. J.L. is supported by an Alzheimer’s Association Research Fellowship and a Warren Alpert Distinguished Scholar Award. B.P. was supported by American Heart Association and BrightFocus Post-Doctoral Fellowships. A.C.M. was supported in part by NIH grant T32GM007601 and NSF graduate fellowship. N.L.F. was supported by NIH grants R01GM147677 and R01NS116176. C.J.D. is supported by NIH grants R01NS105756, R21AG064940, and R01NS127187, Target ALS, and the Robert Packard Center for ALS Research. J.S. is supported by grants from The Packard Center for ALS Research at Johns Hopkins, Target ALS, The Association for Frontotemporal Degeneration, the Amyotrophic Lateral Sclerosis Association, the Office of the Assistant Secretary of Defense for Health Affairs through the Amyotrophic Lateral Sclerosis Research Program (W81XWH-20-1-0242).

## Author Contributions

Conceptualization: L.G., J.R.M, K.E.C., H.W., J.D.R., H.M.O., J.L., B.L.L., Y.S., E.J.H., A.C., N.L.F., S.M., C.J.D., and J.S. Methodology: L.G., J.R.M, J.C.M., K.E.C., H.W., J.D.R., H.M.O., J.L., B.L.L., La.G., A.C.M., T.P., A.M.G., Z.D., A.C., N.L.F., S.M., C.J.D., and J.S. Validation: L.G., J.R.M, J.C.M., K.E.C., H.W., J.D.R., H.M.O., J.L., B.L.L., La.G., E.R., K.M.K., A.C.M., T.P., B.P., A.M.G., Z.D., J.L.C., A.S., G.P., E.L., C.E., Y.S., E.J.H., A.C., N.L.F., S.M., C.J.D., and J.S. Formal analysis: L.G., J.R.M, J.C.M., K.E.C., H.W., J.D.R., La.G., A.C.M., T.P., A.C., N.L.F., S.M., C.J.D., and J.S. Investigation: L.G., J.R.M, J.C.M., K.E.C., H.W., J.D.R., H.M.O., J.L., B.L.L., La.G., E.R., K.M.K., A.C.M., T.P., B.P., A.M.G., Z.D., J.L.C., A.S., G.P., E.L., C.E., A.C., C.J.D. Resources: L.G., J.R.M, J.C.M., K.E.C., J.D.R., H.M.O., J.L., B.L.L., La.G., A.C.M., T.P., B.P., Z.D., A.S., G.P., E.L., C.E., A.C., N.L.F., S.M., C.J.D., and J.S. Data Curation: L.G., J.R.M, J.C.M., K.E.C., J.D.R., La.G., A.C.M., T.P., A.C., N.L.F., S.M., C.J.D., and J.S. Writing – Original Draft: L.G., J.R.M, K.E.C., J.D.R., La.G., N.L.F., S.M., C.J.D., and J.S. Writing - Review & Editing: L.G., J.R.M, J.C.M., K.E.C., H.W., J.D.R., H.M.O., J.L., B.L.L., La.G., E.R., K.M.K., A.C.M., T.P., B.P., A.M.G., Z.D., J.L.C., A.S., G.P., E.L., C.E., Y.S., E.J.H., A.C., N.L.F., S.M., C.J.D., and J.S. Visualization: L.G., J.R.M, J.C.M., K.E.C., J.D.R., La.G., A.C.M., T.P., N.L.F., S.M., C.J.D., and J.S. Supervision: L.G., A.C., N.L.F., S.M., C.J.D., and J.S. Project administration: L.G., A.C., N.L.F., S.M., C.J.D., and J.S. Funding acquisition: L.G., J.R.M, K.E.C., H.M.O., J.L., A.C.M., B.P., A.M.G., N.L.F., C.J.D., and J.S.

## STAR Methods

### KEY RESOURCES TABLE

| REAGENT or RESOURCE | SOURCE | IDENTIFIER |
| --- | --- | --- |
| **Antibodies** |  |  |
| goat MAP2 | Phosphosolutions | Cat# 1099<br>RRID:AB_2752241 |
| mouse G3BP1 | Santa Cruz Biotechnology | Cat# sc-365338<br>RRID:AB_10846950 |
| rabbit FUS | Proteintech | Cat# 11570-1-AP<br>RRID:AB_2247082 |
| Alexa Fluor-647donkey anti-goat | Jackson ImmunoResearch | Cat# 705-605-003<br>RRID:AB_2340436 |
| Alexa Fluor-594 donkey anti-mouse | Jackson ImmunoResearch | Cat# 715-585-150<br>RRID:AB_2340854 |
| Alexa Fluor-488 donkey anti-rabbit | Jackson ImmunoResearch | Cat# 711-545-152<br>RRID:AB_2313584 |
| **Bacterial and Virus Strains** |  |  |
| *E. coli* BL21 DE3 (RIL) cells | Agilent | Cat# 230245 |
| *E. coli* One Shot™ BL21 Star™ (DE3) cells | Thermo Fisher | Cat# C601003 |
| **Chemicals, Peptides, and Recombinant Proteins** |  |  |
| EDTA-free Protease Inhibitor Cocktail | Roche | Cat# 11873580001 |
| Lysozyme | Sigma-Aldrich | Cat#L6876 |
| mTeSR+ | StemCell Technologies | 5825 |
| DMEM:F12 | Corning | 15090CV |
| Neurobasal | Gibco | 12348017 |
| NEAA | Gibco | 11-140-050 |
| Glutamax | Gibco | 35050061 |
| N2 | Gibco | 17502048 |
| B27 | Gibco | 17504044 |
| SB431542 | StemCell Technologies | 72234 |
| LDN-193189 | Sigma-Aldrich | SML0559 |
| Retinoic Acid (RA) | Sigma-Aldrich | R2625 |
| Smoothened-Agonist (SAG) | Cayman Chemical | 11914 |
| SU5402 | Cayman Chemical | 13182-5 |
| DAPT | Cayman Chemical | 13197 |
| TrypLE | Gibco | 12604013 |
| DNaseI | Invitrogen | 18-047-019 |
| BDNF | PeptoTech | 450-02 |
| GDNF | PeptoTech | 450-10 |
| CNTF | PeptoTech | 450-13 |
| Ascorbic acid | Sigma-Aldrich | A4403 |
| Lipofectamine RNAiMAX | Invitrogen | 13-778-075 |
| OptiMEM | Gibco | 51-985-091 |
| Sodium Arsenite | Sigma-Aldrich | 1062771000 |
| Tunicamycin | Sigma-Aldrich | T7765 |
| DMSO | Sigma-Aldrich | D4540 |
| PBS | Gibco | 10010023 |
| 32% PFA | Electron Microscopy Sciences | 15714-S |
| Donkey Serum | Jackson ImmunoResearch | 017-000-121 |
| Triton x-100 | Sigma-Aldrich | T8787 |
| Prolong Glass mounting media | Invitrogen | P36981 |
| Bovine Growth Serum | Cytiva HyClon | SH30541.03HI |
| DMEM, high glucose, pyruvate, no glutamine | Gibco | 10313039 |
| GlutaMAX™ Supplement | Gibco | 35050061 |
| Heparin sodium salt | Sigma-Aldrich | H3149-10KU |
| Animal-Free Recombinant Human EGF | PeptoTech | AF-100-15 |
| RNasin® Ribonuclease Inhibitor | Promega | N2111 |
| Recombinant Human FGF-basic (154 a.a.) | PeptoTech | 100-18B |
| GST-FUS | This paper | N/A |
| GST-FUS<sup>C428A:C433A:C444A:C447A</sup> | This paper | N/A |
| GST-FUS<sup>F305L:F341L:F359L:F368L</sup> | This paper | N/A |
| GST-FUS<sup>371X</sup> | This paper | N/A |
| GST-FUS<sup>P525L</sup> | This paper | N/A |
| GST-FUS<sup>R244C</sup> | This paper | N/A |
| GST-FUS<sup>R216C</sup> | This paper | N/A |
| pHis-TEV | This paper | N/A |
| His-FUS<sub>269-454</sub> | This paper | N/A |
| TDP-43-MBP-his | This paper | N/A |
| TDP-43<sup>5FL</sup>-MBP-his | This paper | N/A |
| **Critical Commercial Assays** |  |  |
| CellTiter-Glo kit | Promega | G7570 |
| QuikChange Site-Directed Mutagenesis Kit | Agilent | Cat# 210518 |
| NEBNext® Small RNA Library Prep Set for Illumina® | New England Biolabs | Cat# E7330S |
| **Experimental Models: Cell Lines** |  |  |
| HEK293 cells | ATCC | 293 [HEK-293]<br>CRL-1573 |
| ReNcell® VM Human Neural Progenitor Cell Line | Millipore | SCC008 |
| FUS R521G iPSC line | Cedars-Sinai RMI iPSC Core | CS37iALS-FUSn2 |
| CS06 iPSC line | Cedars-Sinai RMI iPSC Core | CS06iCTR-n2 |
| **Oligonucleotides** |  |  |
| See Table S2 for RNA oligonucleotides | Horizon Discovery Ltd or IDT | N/A |
| **Recombinant DNA** |  |  |
| GST-FUS | Sun et al.<sup>10</sup> | N/A |
| GST-FUS<sup>C428A:C433A:C444A:C447A</sup> | This Paper | N/A |
| GST-FUS<sup>F305L:F341L:F359L:F368L</sup> | This Paper | N/A |
| GST-FUS<sup>371X</sup> | This Paper | N/A |
| GST-FUS<sup>P525L</sup> | Sun et al.<sup>10</sup> | N/A |
| GST-FUS<sup>R244C</sup> | This Paper | N/A |
| GST-FUS<sup>R216C</sup> | This Paper | N/A |
| pHis-TEV | Cupo and Shorter<sup>70</sup> | N/A |
| Tet-On 3G Inducible Expression System | Takarabio | 631168 |
| ssTDP-43 | This Paper | N/A |
| optoFUS | This Paper | N/A |
| pHR:SFFVp:TDP43C:mCherry:SspB | Bracha et al.<sup>39</sup> | Addgene: 122669 |
| pHR:SFFVp:iLid:eGFP:FTHi | Bracha et al.<sup>39</sup> | Addgene: 122149 |
| pJ4M/TDP-43 plasmid | Wang et al.<sup>59</sup> | Addgene 104480 |
| **Software and Algorithms** |  |  |
| ImageJ | NIH | https://imagej.nih.gov/ij/;<br>RRID:<br>SCR_003073 |
| GraphPad Prism | GraphPad Software Inc | https://www.graphpad.com/scientific-software/prism/; RRID:<br>SCR_002798 |
| Bowtie | Johns Hopkins University | https://bowtie-bio.sourceforge.net/index.shtml |
| HOMER | UCSD | http://homer.ucsd.edu/homer/motif/ |
| Nikon Elements | Nikon | NIS-Elements |

### RESOURCE AVAILABILITY

#### Lead contact

Further information and requests for resources and reagents should be directed to and will be fulfilled by the lead contact, James Shorter.

#### Materials availability

Plasmids newly generated in this study will be made readily available to the scientific community. We will honor requests in a timely fashion. Material transfers will be made with no more restrictive terms than in the Simple Letter Agreement or the Uniform Biological Materials Transfer Agreement and without reach through requirements.

#### Data and code availability

Any additional information required to reanalyze the data reported in this paper is available from the lead contact upon request.

### EXPERIMENTAL MODEL AND SUBJECT DETAILS

#### HEK293 cell culture

HEK293 cells (female, purchased from ATCC) were maintained at 37°C and 5% CO_2_ in DMEM (high glucose, pyruvate) (Thermo Fisher Scientific) supplemented with GlutaMAX (Thermo Fisher Scientific) and 10% Bovine Growth Serum (Cytiva HyClon). Transfections were performed using Lipofectamine 3000 (Thermo Fisher Scientific) according to manufacturer’s instructions following cell seeding onto coverslips or culture plates coated with 50mg/mL collagen (Gibco) and overnight incubation at 37°C/5% CO_2_.

#### ReNcell^®^ VM human neural progenitor cell culture

ReNcell^®^ VM human neural progenitor cells (male, purchased from Millipore) were maintained at 37°C and 5% CO_2_ in DMEM/F12 (Gibco) supplemented with GlutaMAX, B27 (Gibco), 2ng/mL heparin (Sigma-Aldrich), 20ng/mL bFGF (PeproTech) and 20ng/mL hEGF (PeproTech). Neuronal differentiation was performed as previously described^15^ and differentiated neurons were maintained at 37°C and 5% CO_2_/5% O_2_ prior to lentiviral transduction.

#### Induced pluripotent stem cell (iPSC) culture

Induced Pluripotent Stem Cell lines CS37iALS-FUSn2 (female, 37 years old at time of collection) and CS06iCTR-n2 (female, 82 years old at the time of collection) were obtained from the Cedars-Sinai RMI iPSC Core, cultured in Matrigel (Corning) and mTeSR+ (StemCell Technologies) and kept in a humidified chamber with regulated levels of CO2 (5%) and temperature (37°C). All procedures for iPSC culture maintenance and differentiation were performed as described.^71–73^

#### iPSC differentiation

For differentiation, 1×10^6^ iPSCs were plated in 6 well plates. Once cells reached ∼90% confluency media was changed from mTeSR+ to N2B27 media (50% DMEM:F12, 50% Neurobasal, plus NEAA, Glutamax, N2 and B27; all from Gibco) plus 10μM SB431542 (StemCell Technologies), 100nM LDN-193189 (Sigma-Aldrich), 1μM RA (Sigma-Aldrich) and 1μM Smoothened-Agonist (SAG, Cayman Chemical). Media was changed daily for a total of 6 days. Cells were then switched to N2B27 including 1μM RA, 1μM SAG, 4μM SU5402 (Cayman Chemical) and 5μM DAPT (Cayman Chemical) and media was changed daily until day 13.

Neurons were dissociated at day 14 using TrypLE and DNAseI, plated in Matrigel-coated 24-well plates with glass coverslips for confocal imaging studies and Matrigel-coated 96-well white plates for viability assays. Cells were fed every 2 days and maintained for 13 additional days in Neurobasal media + NEAA, Glutamax, N2, B27, plus 10ng/mL BDNF, GDNF, CNTF (all from PeproTech) and 0.2μg/ml Ascorbic acid (Sigma-Aldrich).

### METHOD DETAILS

#### Cloning

QuikChange Site-Directed Mutagenesis Kit (Agilent) was used to generate mutant plasmids (i.e., _GST-FUS1-214, GST-FUSC428A:C433A:C444A:C447A, GST-FUSF305L:F341L:F359L:F368L, GST-FUS371X,_ GST-FUS^P525L^, GST-FUS^R244C^, GST-FUS^R521G^ and GST-FUS^R216C^) according to the manufacturer’s instructions. All GST-FUS constructs have a TEV cleavage site between GST and FUS as described.^10^ Mutations were verified by DNA sequencing.

pJ4M/TDP-43 encoding TDP-43-MBP-his with a TEV cleavage site between TDP-43 and MBP was from Addgene.^59^ The 5FL (F147L:F149L:F194L:F229L:F231L) mutant was generated via QuikChange Multi Site-directed Mutagenesis (Agilent) and verified via Sanger sequencing.

All doxycycline-inducible expression constructs, including FUS-SspB mutants, optoFUS and ssTDP43, were generated through Gibson Assembly (HiFi DNA Assembly Master Mix, NEB) of PCR-generated fragments inserted at the NotI/EcoRI restriction enzyme sites of a Tet3G base vector (synthesized by GeneWiz). Synthesized gBlocks (IDT) containing 4FL and 4CA point mutations were used as templates for PCR of fragments used to assembly FUS-SspB mutants.

Plasmids containing MBP-tagged FUS (Plasmid #98651, Addgene) were used as templates to generate WT FUS-SspB and optoFUS constructs. Previous-generation optoTDP43 constructs ^15^ containing TDP-43 coding sequences were used as PCR templates to generate ssTDP43 constructs. For generation of lentiviral transfer vectors, PCR-generated fragments were inserted at the BsrGI/BamHI restriction enzyme sites by Gibson Assembly of a third-generation base lentiviral vector described previously^15^ for human synapsin promoter-driven expression of target proteins.

#### Purification of TEV protease

TEV protease was purified as described.^70^

#### Purification of GST-FUS

_GST-FUS, GST-FUS1-214, GST-FUSC428A:C433A:C444A:C447A, GST-FUSF305L:F341L:F359L:F368L, GST-_FUS^371X^, GST-FUS^P525L^, GST-FUS^R244C^, and GST-FUS^R216C^ were purified as described.^10^ Briefly, N-terminally tagged GST-FUS was overexpressed in BL21(DE3)RIL *E. coli*. The *E. coli* cells were then lysed by sonication on ice in PBS and protease inhibitors (cOmplete, EDTA-free, Roche Applied Science). The protein was purified over Glutathione Sepharose 4 Fast Flow beads (GE Healthcare) and eluted from the beads using FUS assembly buffer (50mM Tris-HCl pH 8, 200mM trehalose, 1mM DTT, and 20mM reduced glutathione).

#### Purification of his-tagged FUS_269-454_ for NMR experiments

His-tagged FUS_269-454_ was expressed BL21*(DE3) (Life Technologies) in M9 minimal media with ^15^N ammonium chloride for isotopic labeling. Cultures were grown at 37°C until an OD_600_ of 0.6-1 and induced with 1mM IPTG for 4h and cells were harvested by centrifugation. FUS_269-454_ was purified as described.^33^ Briefly, bacterial pellets were resuspended in 20mM sodium phosphate, 1M NaCl, 10mM imidazole pH 7.4 with protease inhibitor tablets (Pierce A32963). The lysate was clarified by centrifugation at 20,000rpm for 1h at 4^°^C, filtered, and applied to a 5mL HisTrap column. The protein was eluted with a gradient of 10-300mM imidazole. The His-tag was cleaved by TEV protease containing a histidine tag, and the protein was dialyzed overnight into 20mM sodium phosphate, 1M NaCl, 10mM imidazole pH 7.4. The protein was filtered and applied to a 5mL HisTrap column to remove the His-tag and TEV protease.

#### Bacterial growth and recombinant protein purification for TDP-43-MBP-his utilized in PS assays

Wild-type (WT) and 5FL TDP-43-MBP-his expression plasmids were transformed into *E. Coli* One Shot^TM^ BL21 Star^TM^ (DE3) cells (ThermoFisher). Transformed *E. coli* were grown at 37°C in 1L of LB media supplemented with 0.2% dextrose and 50μg/mL kanamycin until OD_600_=0.5-0.6. Cells were then incubated at 4°C for 30-45min. Protein expression was induced with addition of 1mM IPTG and then bacterial cultures were incubated for 16h at 16°C. Cells were collected by centrifugation. Cell pellets were resuspended in 1M NaCl, 20mM TrisHCl (pH 8.0), 10mM imidazole, 10% glycerol, and 2.5mM 2-mercaptoethanol and supplemented with cOmplete, EDTA-free protease inhibitor cocktail tablets (Roche), then lysed via sonication. Cell lysates were centrifuged at 48,384rcf at 4°C for 1h. Filtered lysate was purified via FPLC using a XK 50/20 column (Cytiva) packed with Ni-NTA agarose beads (Qiagen), which were equilibrated in the resuspension buffer. Protein was recovered via a 0-80% gradient elution using 1M NaCl, 20mM TrisHCl (pH 8.0), 10mM imidazole, 10% glycerol and 2.5mM 2-mercaptoethanol as the base buffer and 1M NaCl, 20mM TrisHCl (pH 8.0), 500mM imidazole, 10% glycerol and 2.5mM 2-mercaptoethanol as the elution buffer. Eluted protein was concentrated using Amicon Ultra-15 centrifugal filters, MWCO 50kDa (Millipore), filtered and further purified with size-exclusion chromatography using a 26/600 Superdex 200 pg column (Cytiva) equilibrated with 300mM NaCl, 20mM TrisHCl (pH 8.0) and 1mM DTT. The second out of three peaks, as evaluated by absorbance at 280nm, was collected,^59^ spin concentrated as above, aliquoted, flash frozen in liquid nitrogen, and stored at −80°C until further use.

#### Bacterial growth and recombinant protein purification for TDP-43-MBP-his utilized in aggregation assays

TDP-43-MBP-his was purified as described.^13^ WT and 5FL TDP-43 expression plasmids were transformed into *E. Coli* BL21-CodonPlus (DE3)-RIL competent cells (Agilent). Transformed *E. coli* were grown in small cultures in LB with kanamycin (50µg/mL) and chloramphenicol (34µg/mL) at 37°C for approximately 4h. The cultures were then transferred to 1L of LB media supplemented with both antibiotics and glucose (0.2% w/v) and grown at 37°C until OD_600_∼0.5. Protein expression was induced with addition of 1mM IPTG and then bacterial cultures were incubated for 16h at 16°C. Cells were harvested by centrifugation, resuspended in resuspension/wash buffer (20mM Tris-HCl pH 8.0, 1M NaCl, 10mM imidazole, 10% glycerol, 1mM DTT, 5µM Pepstatin A, 100µM PMSF, and cOmplete, EDTA-free, Roche Applied Science protease inhibitors), and lysed by lysozyme (1 mg/mL) and sonication. Cell lysates were centrifuged at 30,966rcf at 4°C for 20min. The protein was purified over Ni-NTA resin (QIAGEN) and eluted from the resin using elution buffer (wash buffer except with 300mM imidazole rather than 10mM imidazole). The protein was further purified over amylose resin (NEB) and eluted with 20mM Tris-HCl pH 8.0, 1M NaCl, 10mM imidazole, 10% glycerol, 1mM DTT, 5µM Pepstatin A, 100µM PMSF, and 10mM maltose. The protein was concentrated using Amicon Ultra-15 centrifugal filters, MWCO 50kDa (Millipore), aliquoted, flash frozen in liquid nitrogen, and stored at −80°C until further use.

#### RNA-Seq

RNA that was bound to GST-FUS during protein purification was extracted by adding DNase I and then Proteinase K to the sample followed by phenol-chloroform extraction, and precipitation in 100% ethanol with 70% ethanol wash. For preparing cDNA libraries for high-throughput sequencing, we used the NEBNext® Small RNA Library Prep Set for Illumina® and followed the manufacturer’s instructions. Library quality was checked with the Agilent 2100 BioAnalyzer. The sample was sequenced on the Illumina HiSeq2000 platform. The resulting sequences were aligned to human genome and *E. coli* genome using Bowtie and the annotated peaks were analyzed by a program HOMER for motif finding.^74,75^

#### RNA oligonucleotides

RNA and fluorescein labeled RNA were purchased from Horizon Discovery Ltd or Integrated DNA Technologies (IDT) (Table S2).

#### FUS fibril assembly

_For GST-FUS, GST-FUSC428A:C433A:C444A:C447A, and GST-FUSF305L:F341L:F359L:F368L, GST-_FUS^P525L^, GST-FUS^R244C^, GST-FUS^R216C^, and GST-FUS^R521G^ fibrillization was initiated by addition of TEV protease to GST-FUS (5µM) in FUS assembly buffer (50mM Tris-HCl pH 8, 200mM trehalose, 1mM DTT, 0.2U/µL RNasin® [Promega], and 20mM glutathione) in the presence or absence of 20µM RNA.^10,50,51^ For the dose-response curves in Figure 3H-K, RNA was dosed from 0.01-1000µM. Fibrillization reactions were incubated at 25°C for 100 min without agitation. FUS^371X^ took longer to fibrillize, and its fibrillization was initiated by addition of TEV protease to GST-FUS^371X^ (10µM) in the presence or absence of 40µM RNA at 25°C for 24h with agitation at 1200rpm.

Turbidity was used to assess fibrillization by measuring absorbance at 395nm. Turbidity of FUS+buffer without TEV condition was subtracted and the resulting absorbance was then normalized to the maximum turbidity of FUS aggregation without RNA to determine the relative extent of fibrillization. For sedimentation analysis, reactions were centrifuged at 16,100g for 10min at 4°C. Supernatant and pellet fractions were then resolved by sodium dodecyl sulfate polyacrylamide gel electrophoresis (SDS–PAGE) and stained with Coomassie Brilliant Blue, and the amount in either fraction (% total) was determined by densitometry in comparison to known quantities of the RBP in question. For electron microscopy, fibrillization reactions (10μl) were absorbed onto glow-discharged 300-mesh Formvar/carbon coated copper grids (Electron Microscopy Sciences) and stained with 2% (w/v) aqueous uranyl acetate. Excess liquid was removed, and grids were allowed to air dry. Samples were viewed by a JEOL 1010 transmission electron microscope.

#### FUS fibril disassembly

Fibrils were assembled as above and used for disassembly reactions. 20µM RNA were added to _preformed GST-FUS, GST-FUSC428A:C433A:C444A:C447A, GST-FUSF305L:F341L:F359L:F368L, GST-_ FUS^P525L^, GST-FUS^R244C^, GST-FUS^R216C^, or GST-FUS^R521G^ fibrils and 40µM RNA were added to preformed GST-FUS^371X^ fibrils to disassemble fibrils. Turbidity, sedimentation analysis, and EM were used to monitor the progress of disaggregation. For turbidity, the absorbance was normalized to that of the fully assembled FUS fibrils before addition of RNA to determine the relative extent of disaggregation. Sedimentation analysis and EM were performed as above.

#### FUS droplet formation

FUS droplets were formed by incubating GST-FUS at indicated concentration in FUS assembly buffer (50mM Tris-HCl pH 8, 200mM trehalose, 1mM DTT, 0.2U/µL RNasin®, and 20mM glutathione) for 2-4h at room temperature (∼23°C±2°C). Protein samples were then spotted onto a coverslip and imaged by Differential interference contrast (DIC) microscopy.

#### Single molecule FRET

For smFRET measurements, the details of instrumentation and PEGylated slide preparation were as described.^32,76^ Briefly, the microfluidic sample chamber was created between the plasma-cleaned slide and the coverslip coated with polyethylene glycol (PEG) and biotin-PEG. Annealed RNA molecules were immobilized on the PEG-passivated surface via biotin-neutravidin interaction. All smFRET measurements were carried out in imaging buffer containing an oxygen scavenger system to stabilize fluorophores (10mM Tris–HCl, pH 7.5, 100mM KCl, 10mM trolox, 0.5% (w/v) glucose, 1mg/mL glucose oxidase and 4g/ml catalase).^76^ All smFRET assays were performed at room temperature (∼23°C±2°C). Wide-field prism-type total internal reflection fluorescence (TIRF) microscopy was used with a solid-state 532nm diode laser to generate an evanescent field of illumination to excite the fluorophores (Cy3 or Cy5) at the sample chamber. Fluorescence signals from Cy3 (donor) and Cy5 (acceptor) were simultaneously collected using a water immersion objective and sent to a charge-coupled device (CCD) camera after passing through the dichroic mirror (cut off = 630nm). Movies were recorded over different regions of the imaging surface with a time resolution of 100ms as a stream of imaging frames. FRET histograms were built by collecting FRET values from over 5000 molecules in 20 different fields of view (21 frames of 20 short movies). Long movies (1200 frames, i.e., 120s) were recorded to look through the molecular behavior using MATLAB script.

#### NMR spectroscopy methods

NMR experiments were recorded on a Bruker Avance 850 MHz ^1^H Larmor frequency spectrometer with HCN TCl z-gradient cryoprobe. All experiments were carried out at 310K. Data were processed using NMRPipe software package^77^ and then visualized using NMRFAM-Sparky.^78^ For NMR experiments, the protein was dialyzed into 20mM NaPi (pH 6.75), 150mM NaCl. Assignments were kindly provided by Frederic Allain and Fionna Loughlin.^33^ Experiments were conducted in 20mM NaPi (pH 6.75), 150mM NaCl, 10% ^2^H_2_O in the presence of 60µM FUS_269-454_ with 60µM RNA (i.e., 1:1).

#### Fluorescence anisotropy

Fluorescein-labeled RNAs (8nM) were added into GST-FUS, GST-FUS^C428A:C433A:C444A:C447A^, GST-FUS^F305L:F341L:F359L:F368L^, or GST-FUS^371X^ at indicated concentration in FUS assembly buffer (50mM Tris-HCl pH 8, 200mM trehalose, 1mM DTT, and 20mM glutathione) in the presence of RNasin®. Anisotropy (excitation 470 nm, emission 520 nm) was measured in 96-well plate using an Infinite M1000 plate reader (Tecan). The change in anisotropy was calculated by subtracting the anisotropy of 8nM fluorescein-labeled RNA and the binding curve was fitted using Prism to obtain the *K_D_*.

#### *In vitro* TDP-43 PS inhibition assay

RNA, TDP-43-MBP-his, and TEV protease were thawed on ice. TDP-43-MBP-his was centrifuged at 16,000rcf for 10min at 4°C. RNA was diluted into PS buffer (150mM NaCl, 20mM HEPES pH 7.4) and TDP-43 and TEV were diluted into PS buffer supplemented with 1mM DTT. Equal volumes of TDP-43-MBP-his and RNA were mixed and incubated at room temperature for 15min before adding an equal volume of TEV protease, for final concentrations of 4µM TDP-43, and 0.01 mg/mL TEV protease in PS buffer with 0.67mM DTT and variable amounts of RNA. An Infinite M1000 or Safire2 plate reader (Tecan) was used to scan samples in a UV-transparent half-area 96-well plate (Greiner) at 350nm, once per minute, for 2h at ∼25-30°C. Initial readings (T=0min) were subtracted from final readings (T=120min) then normalized to the “no RNA” controls.

#### *In vitro* TDP-43 PS reversal assay

Equal volumes of TDP-43-MBP-his and TEV protease, and TDP-43 and buffer (negative control) were mixed at room temperature (∼23°C±2°C) for final concentrations of 4.3µM TDP-43, and 0.011mg/mL TEV protease in PS buffer with 1mM DTT, then incubated at room temperature (∼23°C±2°C) for 1.5h to allow for TDP-43 PS. After 1.5h, this solution was transferred to wells in a UV-transparent half-area 96-well plate (Greiner) and scanned once at 350nm in the plate reader. RNAs or Buffer were added to the wells, for final concentrations of 4µM TDP-43, 0.01mg/mL TEV protease in PS buffer with 0.93mM DTT and then the samples were scanned at 350 nm, once per minute for 1h at ∼25-30°C. Background subtraction was performed by subtracting average readings for negative controls (TDP-43 with buffer, no TEV protease) from sample readings at T=0h (no RNA added) and at T=1h. Each sample reading at T=1h was normalized to its own T=0h reading, and then samples were normalized again to the “no RNA” controls.

#### *In vitro* TDP-43 aggregation inhibition assay

TDP-43-MBP-his was thawed on ice and centrifuged for 10min at 21,300rcf at 4°C. TDP-43-MBP-his was buffer exchanged into 166.66mM NaCl, 22.22mM HEPES-NaOH pH 7.0, 1.11mM DTT (Bio-Rad Micro Bio-Spin Chromatography Columns, following manufacturer’s instructions) and concentration was determined via NanoDrop, e_280_=114250 cm^-1^M^-1^. TDP-43-MBP-his and RNA (or water for controls without RNA) were added to buffer to achieve final concentrations of 5µM TDP-43, 150mM NaCl, 20mM HEPES-NaOH pH 7.0, 1mM DTT, and the indicated RNA concentration. Samples were incubated at room temperature (∼23°C±2°C) for 15min, after which TEV protease was added at a final concentration of 2.5µg/mL (TEV protease elution buffer was added for the No TEV control) to remove the MBP-his tag. An Infinite M1000 Tecan plate reader was used to assess turbidity once per minute at 395 nm in a nonbinding 96-well plate (Greiner) over 16h at ∼25-30°C. The data was standardized by subtracting out the initial reading at t=1min from each respective condition. Data was then normalized to the respective No RNA control. Area under the curve analysis was used to compare the extent of aggregation for each condition (GraphPad Prism).

#### *In vitro* TDP-43 aggregation reversal assay

TDP-43-MBP-his was thawed on ice and centrifuged for 10min at 21,300rcf at 4°C. TDP-43-MBP-his was buffer exchanged into 150mM NaCl, 20mM HEPES-NaOH pH 7.0, 1mM DTT (Bio-Rad Micro Bio-Spin Chromatography Columns, following manufacturer’s instructions) and concentration was determined via NanoDrop, e_280_=114250 cm^-1^M^-1^. TDP-43-MBP-his was diluted into buffer to achieve a final concentration of 4µM TDP-43, 150mM NaCl, 20mM HEPES-NaOH pH 7.0, 1mM DTT. TEV protease was added at a final concentration of 2.5µg/mL for WT TDP-43 to remove the MBP-his tag. Due to slower aggregation kinetics, to achieve preformed aggregates in an equivalent timeframe, TEV protease was added at a final concentration of 10µg/mL for 5FL TDP-43. An Infinite M1000 Tecan plate reader was used to assess turbidity once per minute at 395nm in a nonbinding 96 well plate (Greiner) over 6h at ∼25-30°C. After 6h, turbidity readings were paused. RNA (or water for controls without RNA) was added to samples, resulting in final concentrations of 40µM RNA (for samples with RNA), 3.648µM TDP-43, 136.8mM NaCl, 18.24mM HEPES-NaOH pH 7.0, 0.912mM DTT. Sedimentation was performed by taking samples at the end timepoint. Input samples were taken directly from the sample. Samples were centrifuged for 10min at 21,300rcf at RT. The supernatant of these centrifuged samples was taken as the supernatant sample, while the pellet was resuspended in buffer (136.8mM NaCl, 18.24mM HEPES-NaOH pH 7.0, 0.912mM DTT) for the pellet samples. 3x sample buffer with 2-mercaptoethanol was added to samples, which were boiled at 95°C for 5min. Samples were run on 4-20% Tris-HCl PAGE gels and stained with Coomassie Brilliant Blue. Quantification of bands was performed with Image Studio Lite Ver 5.2. Samples at the end timepoint were also imaged by brightfield microscopy with 100x objective (EVOS M5000)

#### iPSC neuronal culture treatment and immunostaining analyses

Oligo treatments started on day 13 after plating (DIV27) and lasted 24h. 2’OMe_RNA oligos were transfected using Lipofectamine RNAiMAX (Invitrogen) according to manufacturer’s instructions. Briefly, each oligo was diluted in OptiMEM (Gibco) and combined with 1µl Lipofectamine/well, the mixture was incubated at RT for 10min and then added dropwise to the cells to a final concentration of 500nM. Neurons were always fixed at day 14 after plating (DIV28). For SG studies, oligo treatment was started on day 13, and then sodium arsenite (Sigma-Aldrich) was added 23h later at a final concentration of 0.5mM, incubated at 37°C for 45min, fixed and stained.

For viability studies, a tunicamycin dose/response curve was performed to determine a concentration that would reduce viability significantly (reduction of >10% compared to vehicle treated) in control neurons. Tunicamycin was dissolved in DMSO (both Sigma-Aldrich), serial dilutions were made in OptiMEM, added dropwise to each well and incubated at 37°C for 24h. Cell viability was measured using the CellTiter-Glo kit (Promega). For oligo experiments, tunicamycin was used at doses of 25μM and 50μM, treatment was started 1h after oligo transfection, incubated for 24h at 37°C and cell viability measured using CellTiter-Glo. For immunofluorescence studies, cells were washed once in PBS (Gibco) and fixed in 4% PFA (Electron Microscopy Sciences) immediately after treatments ended. Cells were kept on PFA for 20min, then washed 3 times in PBS and blocked with 5% Donkey Serum (Jackson ImmunoResearch) plus 0.3% TX-100 (Sigma-Aldrich) in PBS for 30min at room temperature (∼23°C±2°C). Primary antibodies (goat MAP2 1:1000, Phosphosolutions; mouse G3BP1 1:100, Santa Cruz; rabbit FUS 1:300, Proteintech) were diluted in blocking solution and incubated overnight at 4°C. Secondary antibodies (donkey Alexa Fluor, Jackson ImmunoResearch) were used at 1:1000 dilution in blocking solution and incubated for 60min at 30min at room temperature (∼23°C±2°C). All treatments/cell lines were treated and probed simultaneously to decrease variability. Coverslips were mounted in Prolong Glass (Invitrogen).

Images were acquired (10/group) using an A1R Nikon Confocal Microscope and fields of view (FOV) were processed for analyses using Nikon NIS Elements Software. Briefly, SG signal on untreated control neurons was thresholded using a binary layer for 594nm channel (G3BP1) and settings were kept consistent across treatments. Within each FOV, total number of neurons (DAPI+/MAP2+) and SG+ neurons (neurons where G3BP1 signal met the binary thresholding requirements) were counted and percentage of cells with SGs over total number of cells was obtained per each image. SGs per cell values were obtained using the counting tool in NIS Elements only on neurons that were determined to be SG+. SG area and FUS signal intensity was obtained after the binary layer was applied to each image. iPSC image quantifications were analyzed by two-way ANOVA test with FUS genotype (WT and mutant) and Oligo treatment (vehicle, RNA C2 and RNA S1) as variables. Viability assay was analyzed by one-way ANOVA. Significance was set at 0.05 and post-hoc pairwise comparisons with the Bonferroni correction were used for analysis of specific differences in any cases where interactions were significant.

#### Detergent solubility fractionation

For assessment of relative optoFUS and ssTDP43 detergent solubility, cell lysate fractionation was performed as described^15^ with minor modifications. Briefly, cells were first lysed with RIPA buffer (25mM Tris-HCl pH 7.6, 150mM NaCl, 2mM EDTA, 1% NP-40, 1% sodium deoxycholate, 0.1% SDS) supplemented with cOmplete Protease Inhibitor Cocktail (Roche) and phosphatase inhibitor cocktails 2/3 (Sigma-Aldrich) following one wash in ice-cold PBS. After brief sonication (five 3s pulses at 30% amplitude), lysates were then centrifuged at 17,000g at 4°C for 45min and the resulting supernatant was collected as the RIPA-soluble fraction. Protein concentration was determined using the Pierce BCA assay (Thermo Fisher Scientific). Pellets were then washed in RIPA buffer prior to re-centrifugation at 17,000g at 4°C for 45min. The resulting supernatants were then discarded, and pellets were re-suspended in urea buffer (30mM TrisHCl pH 8.5, 7M urea, 2M thiourea, 4% CHAPS) supplemented with cOmplete Protease Inhibitor Cocktail (Roche) and phosphatase inhibitor cocktails 2/3 (Sigma-Aldrich) and sonicated briefly prior to centrifugation at 17,000g at room temperature (∼23°C±2°C). The resulting supernatant was then collected as the RIPA-insoluble, urea soluble fraction and samples were separated by SDS-PAGE prior to western blot analysis.

#### SDS-PAGE/Western blotting

Prior to SDS-PAGE, samples were first diluted in 4X Laemmli sample buffer (Bio-Rad) supplemented with 2-mercaptoethanol (Bio-Rad) and heated at 70°C for 10-15min. Samples were then loaded into 12% or 4-20% Mini-PROTEAN TGX Precast gels (Bio-Rad) and separated by SDS-PAGE. Separated samples were next transferred to PVDF membranes (Bio-Rad) prior to washing (TBS) and blocking with Odyssey Blocking Buffer (Li-Cor). Membranes were then incubated with primary antibodies diluted in Odyssey Blocking Buffer supplemented with 0.2% Tween-20 overnight at 4°C. Primary antibody dilutions consisted of: mouse anti-mCherry (Novus Biologicals, 1:1000), rabbit anti-mCherry (Abcam, 1:1000), rabbit anti-FUS (Proteintech, 1:1000), rabbit anti-TDP43 (Proteintech, 1:1000), mouse anti-α-tubulin (Sigma, 1:10000). The next day, membranes were washed with TBS-T (0.1% Tween-20) and incubated with secondary antibodies (Li-Cor, IRDye 680/800, 1:10000) for 1h at room temperature (∼23°C±2°C) prior to TBS-T washes and imaging (Odyssey CLx imaging system).

#### Immunofluorescence

For immunofluorescent characterization of optoFUS and ssTDP43 inclusions, cells seeded onto collagen-coated coverslips (Thermo Fisher, 50μg/mL) were first fixed for 15min at room temperature (∼23°C±2°C) in 4% PFA following one PBS wash. Three additional PBS washes were then performed prior to a 1h incubation in blocking buffer (0.3% TX-100/5% NDS in PBS) at room temperature (∼23°C±2°C). Cells were then incubated overnight at 4°C with primary antibodies diluted in blocking buffer at the following concentrations: rabbit anti-TAF15/TAFII68 (Bethyl Labs, 1:500), mouse anti-EWSR1 (Santa Cruz, 1:200), rat anti-methylated TLS/FUS (Clone 9G6, Sigma-Aldrich, 1:100), guinea pig anti-MAP2 (Synaptic Systems, 1:1000), rabbit anti-G3BP1 (Proteintech, 1:500), rat anti-phospho-TDP43 (S409/410) (Clone 1D3, Biolegend, 1:200), rabbit anti-SQSTM1/p62 (Abcam, 1:500). The following day, primary antibodies were removed, and cells were exposed to three PBS washes prior to a 1h incubation with secondary antibodies (AlexaFluor 488/594/647, 1:1000) diluted in blocking buffer at room temperature (∼23°C±2°C). Three additional PBS washes were then performed prior to mounting coverslips onto slides using ProLong Diamond Antifade Mountant with DAPI (Invitrogen). Slides were allowed to cure overnight prior to visualization by confocal microscopy.

#### Live-cell imaging

Live-cell imaging experiments were performed on a Nikon Eclipse Ti2 inverted microscope equipped with an X-Light V2 (CrestOptics) spinning disk unit using CFI Plan Apo Lambda 40X dry or CFI Plan Apo VC 60X water immersion objectives (Nikon) and a Prime 95B CMOS camera (Photometrics). Cells were maintained at 37°C and 5% CO_2_ in a Tokai HIT STX stagetop incubator throughout the imaging process. For chronic stimulation paradigms, wells were illuminated (∼0.1-0.3mW, 465nm) using custom-built 6-well, 24-well, 96-well LED panels designed to sit atop the plates in between image acquisition periods using a 5V analog output from a Texas Instruments BNC-2110 triggering device as described.^15^ For acute LIPS experiments, cells expressing iLID cores along with FUS-mCh-SspB mutants were first imaged using only the 594nm laser line to establish baseline FUS-SspB fluorescence intensity and spontaneous condensate assembly. Acute activation sequences (30s or less) were then achieved through dual-channel imaging with the 594nm and 488nm (75% power) laser lines, followed by post-activation image sequences for up to 10min acquired using only 594nm lasers to avoid further activation.

#### Fluorescence recovery after photo-bleaching (FRAP) imaging and analysis

For FRAP analysis of optoFUS and ssTDP43 assemblies, cells expressing these constructs were first imaged prior to light activation of optogenetic proteins to acquire baseline fluorescence recovery rates due to diffusion. Cells were then exposed to light activation for the indicated times and relative dynamics of light-induced condensates/inclusions were determined by FRAP. All imaging was performed on a Nikon A1 laser-scanning confocal microscope utilizing a 60X oil immersion objective (Nikon, CFI Plan Apo Lambda 60X Oil) and Tokai HIT stagetop incubator to maintain cells at 37°C and 5% CO_2_. In brief, 2µm diameter bleaching regions-of-interest (ROIs) were drawn within nuclear compartments (for dark or pre-activation conditions) or around light-induced assemblies. 2-3 baseline images were then acquired prior to photo-bleaching within bleaching ROIs using the 488nm laser line (500ms, 50% power). Post-bleaching image sequences were then acquired for up to five minutes and fluorescence recovery within bleaching ROIs was measured over time. Fluorescence intensity values were normalized to intensities within reference ROIs of the same size drawn in non-bleached cells to control for fluorescence loss resulting from post-bleach imaging. These values were then normalized to each ROI’s minimum and maximum intensities and were plotted as mean recovery rates per condition.

#### Automated image analysis

All automated image analysis was performed in NIS-Elements Advanced Research software (Nikon) using built-in analysis packages. For analysis of FUS-SspB condensate formation following acute light activation protocols, individual ROIs were first drawn around all cells expressing both iLID cores and FUS-SspB mutant constructs in each field-of-view. Baseline FUS-SspB fluorescence intensity was determined in frames prior to light activation. Automated Spot Detection was then used to identify and quantify the number of FUS-SspB droplets within each ROI during and following light activation sequences and the Time Measurement tool was used to export the number of objects per cell over time to Microsoft Excel. Object number values were then normalized to baseline values (prior to light activation), weighted based on baseline FUS-SspB fluorescence intensity (compared to population mean), and plotted over time. For graphs comparing threshold FUS-SspB concentrations required for LIPS, baseline fluorescence values were plotted against the maximum number of objects observed in each individual cell over the time-course of the experiment. C_thresh_ values were determined by calculating the mean baseline fluorescence intensity of the five lowest-expressing cells in each mutant condition that underwent LIPS (defined as the formation of >10 condensates in response to light activation). For quantification of condensate dissociation kinetics, the number of objects identified in each individual cell in the first frame following light removal (T_0_) was set at 100% and values in each successive frame were normalized as a percentage of initial T_0_ values and mean dissociation values were plotted over time. One-phase exponential decay curves were then fit and T½ values for each FUS-SspB mutant were determined using Graphpad Prism 8 software.

For automated analysis of optoFUS normalized aggregation area, individual z-stacks were acquired in 9-16 randomized fields-of-view and maximum intensity projections were generated for analysis. First, binaries for cell nuclei and optoFUS inclusions were generated through fluorescence intensity thresholding of DAPI and mCherry signals respectively (Figure S7A-H). Binary subtraction operations were then performed to generate a new binary layer consisting of mCherry signal with nuclear signal subtracted to remove confounding nuclear optoFUS signal from analysis. The total area of this resulting binary layer (optoFUS inclusions only) was then calculated and normalized to total optoFUS cell area (determined by cell masks based upon mCherry fluorescence) and was presented as normalized aggregation area. Mean aggregation area values were then determined across fields-of-view and plotted as fold-change from control.

For automated quantification of light-induced formation of ssTDP43 inclusions, maximum intensity projections were first generated from z-stacks acquired over at least 6 individual fields-of-view per condition. Automated Spot Detection was then utilized to identify and quantify the number of light-induced condensates per field-of-view over time. These values were then normalized to baseline (prior to light activation) values and plotted as mean increase from baseline over the course of light stimulation. For quantification of ssTDP43 and optoFUS inclusion disassembly, individual inclusions from 6-8 fields-of-view were identified and tracked over time. Here, baseline inclusion area was first determined through automatically or manually drawn ROIs in the first frame acquired following RNA treatments (T_0_). ROI areas were then determined for subsequent frames every 2h for up to 10h, normalized to baseline values and presented as fold change from T_0_ over time. Survival of these inclusion-bearing cells was also manually tracked, and Graphpad Prism 8 was used to generate and compare Kaplan–Meier survival curves between treatment groups. All above analyses were performed blinded.

#### Minigene and splicing assays

For monitoring of TDP-43 splicing function, the CFTR exon 9 minigene assay was performed as previously described^79^ with minor modifications. In brief, HEK293 cells transfected with the CFTR minigene plasmid were treated with siRNA (25nM) or RNA inhibitor oligonucleotides (2.5μM) for 72h prior to cell lysis and RNA extraction using the miRNA Easy Kit (Qiagen). The iScript cDNA Synthesis Kit (Bio-Rad) was then used to generate cDNA from RNA samples and PCR reactions were then performed using cDNA templates and primers flanking exon 9 of the CFTR minigene^79^ prior to separation on a 1% agarose gel. Primer sequences are as follows: Fwd: 5’-CAACTTCAAGCTCGTAAGCCACTGC-3’; Rev: 5’-TAGGATCCGGTCACCAGGAAGTTGGTTAAATCA-3’. Bands were then visualized and imaged using the Chemidoc MP Imaging System (Bio-Rad).

#### QUANTIFICATION AND STATISTICAL ANALYSIS

Quantification is as described in the figure legends. Statistical analyses were performed using the GraphPad Prism (GraphPad Software, Inc.; La Jolla, CA, USA) as described in figure legends.

---

**Figure S1.**
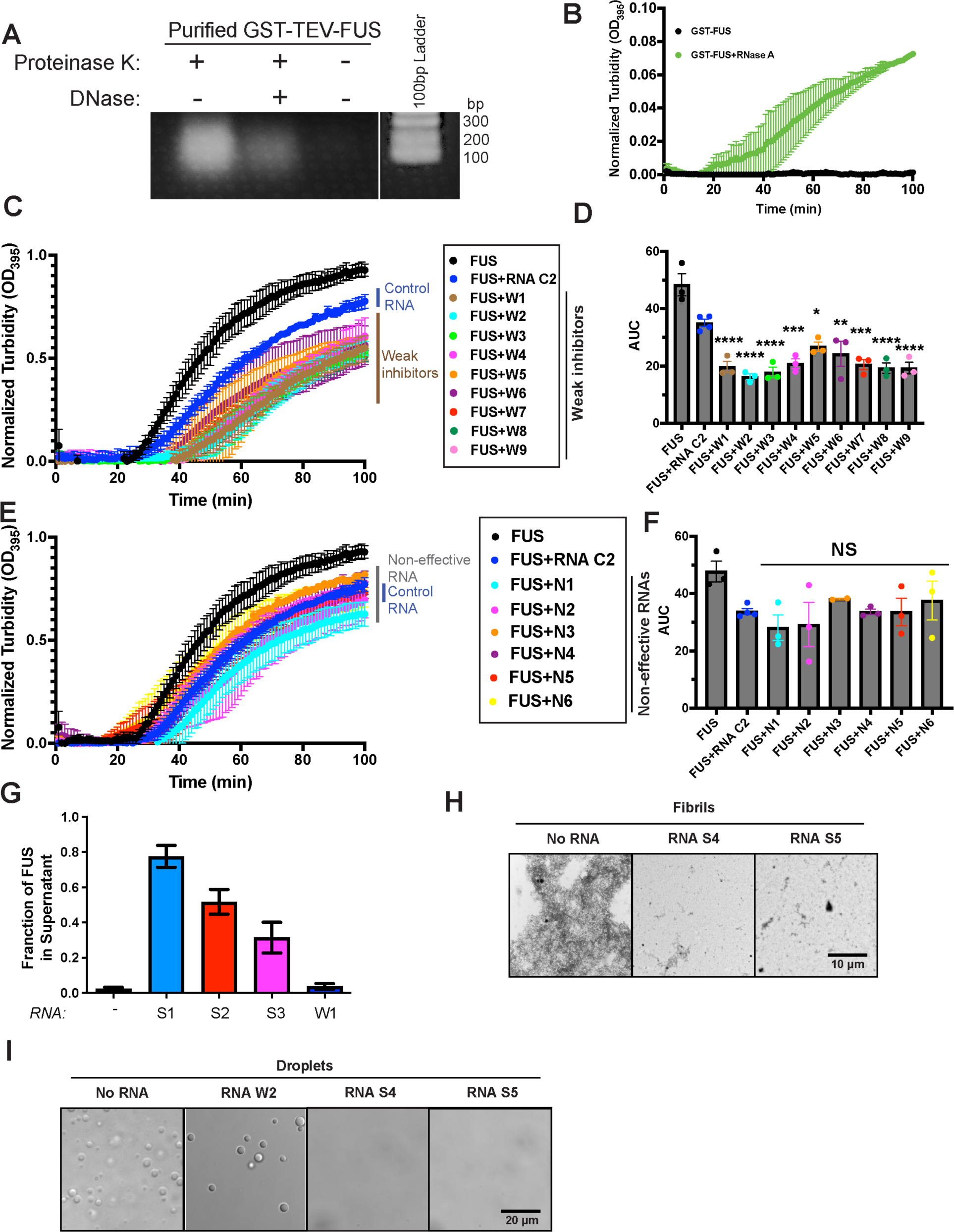
Weak RNA inhibitors inhibit FUS fibrillization but not FUS PS. **(A)** Agarose gel reveals RNA in GST-FUS purified from *E. Coli.* GST-FUS purified from *E. Coli* was treated at 37°C for one hour with proteinase K, proteinase K and DNase, or left untreated. Samples were then analyzed by 1% agarose gel and stained with ethidium bromide. **(B)** GST-FUS (5µM) was incubated without TEV protease in the presence or absence of RNase A. FUS assembly was monitored by turbidity. Values represent means±SEM (n = 2). **(C)** GST-FUS (5µM) was incubated with TEV protease in the presence or absence of weak RNA inhibitor (RNA W1-W9; 20µM) for 0–100min. Turbidity measurements were taken every minute to assess the extent of FUS assembly. The FUS only curve and FUS+RNA C2 curve were plotted from the same data set as in Figure 1B, since experiments in these two figures were run at the same time. Values represent means±SEM (n = 3). **(D)** Area under the curve calculated using Prism for each replicate summarized in (C) quantifies the extent of FUS assembly. The FUS only curve and FUS+RNA C2 curve were plotted from the same data set as in Figure 1B, since experiments in these two figures were run at the same time. Values represent means±SEM (n = 3). One-way ANOVA and Dunnett’s test were used to compare to the RNA C2 condition; ns: p>0.05, *p ≤0.05, **p≤0.01, ***p≤0.001, and ****p≤0.0001. **(E)** GST-FUS (5µM) was incubated with TEV protease in the presence or absence of indicated RNA (20µM) for 0–100min. Turbidity measurements were taken every minute to assess the extent of FUS assembly. The FUS only curve and FUS+RNA C2 curve were plotted from the same data set as in Figure 1B, since experiments in these two figures were run at the same time. Values represent means±SEM (n = 3). **(F)** Area under the curve calculated using Prism for each replicate summarized in (E) quantifies the extent of aggregation. Values represent means±SEM (n = 3). Non-effective RNAs do not show statistical difference compared to RNA C2 in inhibiting FUS assembly. One-way ANOVA and Dunnett’s test were used to compare to the RNA C2 condition; ns: p>0.05. **(G)** GST-FUS (5µM) was incubated with TEV protease in the presence or absence of the indicated RNA (20µM) for 0– 90min. At 90min, reactions were processed for sedimentation analysis. Supernatant fractions were resolved by SDS-PAGE and stained with Coomassie Brilliant Blue. The amount of FUS in the supernatant fraction was determined by densitometry in comparison to known quantities of FUS. Values represent means±SEM (n=3). **(H)** GST-FUS (5µM) was incubated with TEV protease in the presence or absence of strong RNA inhibitor (RNA S4, RNA S5; 20µM) for 100min. Samples were processed for EM at the end of the reaction. Bar, 10μm. **(I)** GST-FUS (10µM) was incubated for 4h in the presence or absence of the indicated RNA (40µM). Droplet formation was assessed by DIC microscopy. Bar, 20μm. Related to Figure 1.

**Figure S2.**
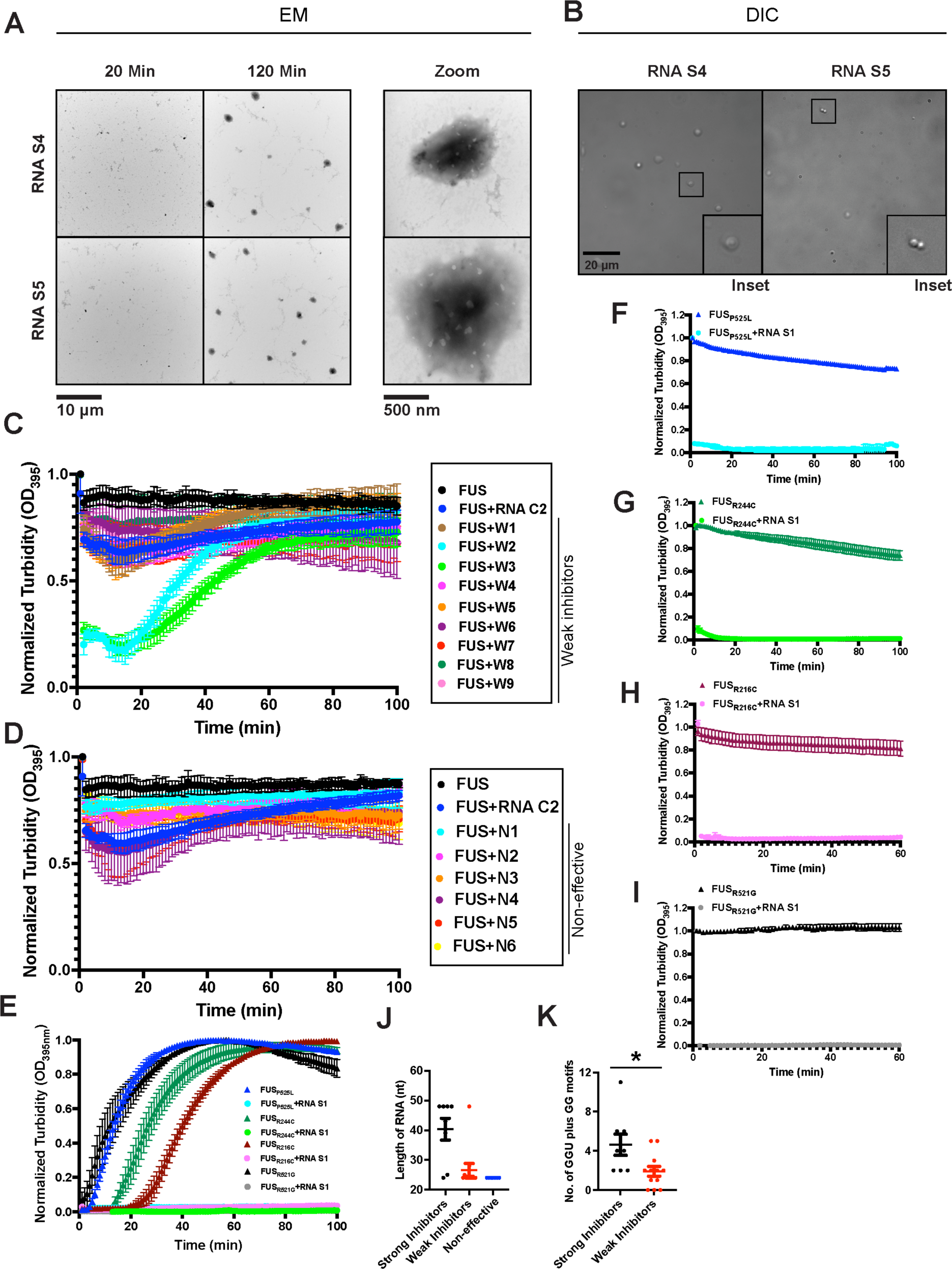
Strong RNA inhibitors prevent and reverse fibrillization of ALS-linked FUS variants. **(A)** GST-FUS (5µM) was incubated with TEV protease for 100min to assemble fibrils. At the end of the reaction, RNA S4 or S5 (20µM) were added to the reaction. Samples were taken after 20min and 120min to visualize the disaggregation products via EM. Bar, 10µm. The right images show higher magnification of the dense protein phase observed in the middle panel. Note the porous structure indicative of hydrogel formation. Bar, 500nm. **(B)** DIC images of the hydrogel sample observed in (A) indicating they are small solid-like drops that do not fuse. Bar, 20μm. **(C)** GST-FUS (5µM) was incubated with TEV protease for 100min to assemble fibrils. At the end of the reaction, water or indicated weak RNA inhibitor (20µM) was added to the reaction. Turbidity measurements were taken every minute to assess the extent of disaggregation. The FUS only curve and FUS+RNA C2 curve were plotted from the same data set as in Figure 1F, since experiments in these two figures were run at the same time. Values represent means±SEM (n = 3-4). **(D)** GST-FUS (5µM) was incubated with TEV protease for 100min to assemble fibrils. At the end of the reaction, water or indicated non-effective RNA (20µM) was added to the reaction. Turbidity measurements were taken every minute to assess the extent of disaggregation. The FUS only curve and FUS+RNA C2 curve were plotted from the same data set as in Figure 1F, since experiments in these two figures were run at the same time. Values represent means±SEM (n = 3-4). **(E)** GST-FUS^P525L^, GST-FUS^R244C^, GST-FUS^R216C^ or GST-FUS^R521G^ (5µM) was incubated with TEV protease in the presence or absence of RNA S1 (20µM) for 0–100min. Fibrillization was assessed by turbidity. Values represent means±SEM (n = 3). **(F-I)** GST-FUS^P525L^ (F), GST-FUS^R244C^ (G), GST-FUS^R216C^ (H), or GST-FUS^R521G^ (I) (5µM) was incubated with TEV protease for 100min to form fibrils. At this time, water, or RNA S1 (20µM) was added to the reaction. Turbidity measurements were taken every minute to assess the extent of disaggregation. Values represent mean ±SEM (n = 3). **(J)** Length distribution of strong RNA inhibitors (n=8), weak RNA inhibitors (n=15), and non-effective RNAs (n=8). Bars represent means±SEM. **(K)** Distribution of the number of GGU motifs plus GG motifs in strong inhibitors (n=8) and weak inhibitors (n=15). Bars represent means±SEM. Unpaired Student’s t-tests were used to compare between groups. \**p*≤0.05. Related to Figure 1.

**Figure S3.**
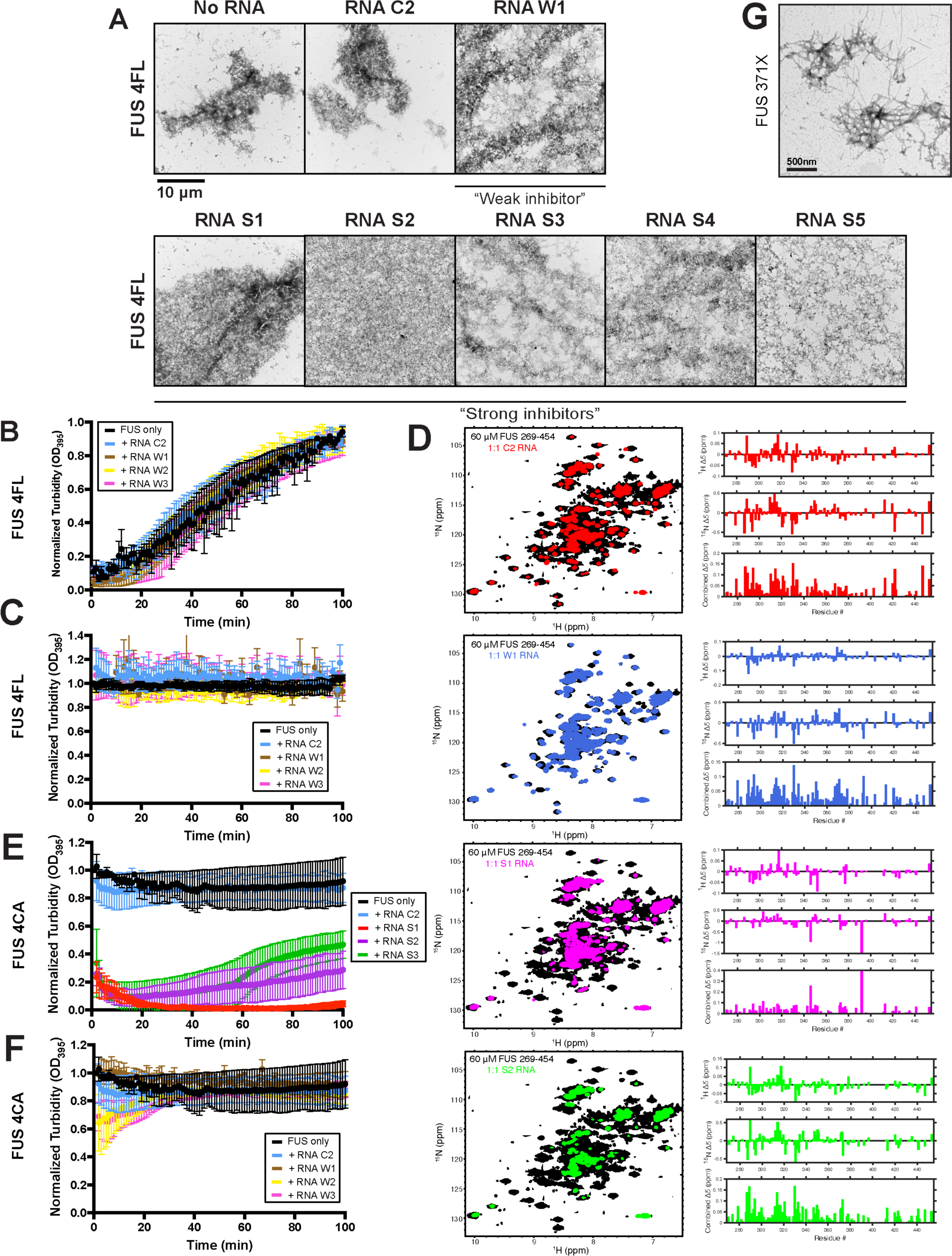
Strong and weak RNA inhibitors engage multiple RNA-binding domains of FUS to antagonize FUS fibrillization. **(A)** GST-FUS_4F-L_ (5µM) was incubated with TEV protease in the presence or absence of the indicated RNA inhibitors or the control C2 RNA (20µM) for 0– 100min. At the end of the fibrillization reactions, samples were processed for EM. Bar, 10μm. **(B)** GST-FUS_4F-L_ (5µM) was incubated with TEV protease in the presence or absence of weak RNA inhibitors W1, W2, or W3 or the control C2 RNA (20µM) for 0–100min. Fibrillization was assessed via turbidity. Values represent means±SEM (n = 3). **(C)** FUS_4F-L_ fibrils (5µM monomer) were treated with water or the indicated RNA (20µM). Disaggregation was assessed by turbidity. Values represent means±SEM (n = 2-3). **(D)** NMR spectra (left) of FUS_269-454_ without and with the addition of the indicated RNA show significantly more line broadening by addition of S1 and S2 RNA inhibitors than C1 or W1, consistent with tighter binding for S1 and S2 compared to C1 and W1. Chemical shift perturbations (right) quantified for these spectra as a function of residue number show perturbations across the entire FUS sequence for all RNAs, even for C1 and W1 RNAs, suggesting RNA binding across multiple FUS domains. **(E)** FUS_4C-A_ fibrils (5µM monomer) were treated with water or the indicated RNA (20µM). Disaggregation was assessed by turbidity. Values represent means±SEM (n = 3). **(F)** FUS_4C-A_ fibrils (5µM monomer) were treated with water or the indicated RNA (20µM). Disaggregation was assessed by turbidity. The FUS only curve and FUS+RNA C2 curve were plotted from the same data set as in (E), since experiments in these two panels were run at the same time. Values represent means±SEM (n = 3). **(G)** GST-FUS_371X_ (10µM) was incubated with TEV protease at 25°C for 24h with agitation at 1200rpm. At the end of the fibrillization reaction, sample was processed for EM. FUS_371X_ forms fibrils like WT FUS, although the kinetics are much slower. Bar, 500nm. Related to Figure 3.

**Figure S4.**
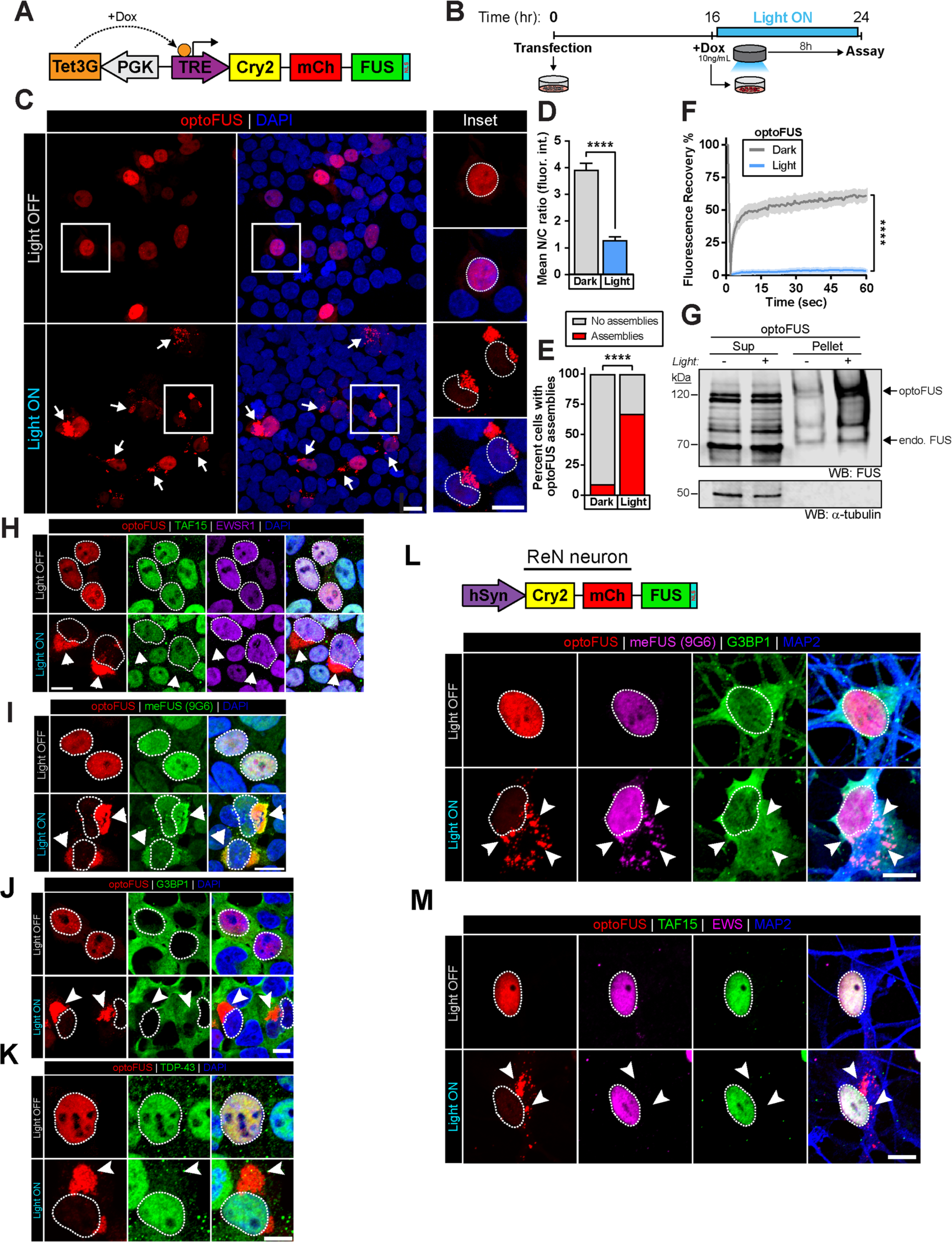
An optogenetic model of FUS-ALS pathology. **(A)** Schematic of the optoFUS construct used in these experiments, in which an N-terminal Cry2olig-mCherry fusion to the full-length FUS protein is expressed under the control of the doxycycline-inducible pTRE3G promoter. **(B)** Light-induction paradigm used to induce optoFUS inclusion formation. **(C)** Representative images of cells optoFUS-expressing cells exposed to 8h of darkness or light. Cell nuclei are circled. Bar, 10µm. **(D)** Immunofluorescence analysis of optoFUS nuclear/cytoplasmic signal following light induction protocol. Values represent means±SEM. *n=*45 cells per group. Unpaired Student’s t-tests were used to compare across groups, ****p<0.0001. **(E)** Quantification of the percentage of cells containing cytoplasmic optoFUS inclusions following 8h of darkness or light. *n=*128-147 cells per group. Unpaired Student’s t-tests were used to compare across groups. **** *p*<0.0001. **(F)** Fluorescence recovery after photobleaching (FRAP) analysis of light-induced inclusions or nuclear optoFUS signal in cells kept in darkness. Values represent means (solid line) ± SEM (shaded area). *n=*15-23 cells. Two-way ANOVA with Sidak post-hoc analysis, **** *p*<0.0001. **(G)** Detergent-solubility fractionation of optoFUS cell lysates collected following 16h of darkness or light. **(H-I)** Immunofluorescence analysis of optoFUS inclusions for co-localization with (H) FTLD-FUS pathological hallmarks TAF15 (green) and EWSR1 (purple) or (I) the ALS-FUS-associated methylated FUS antibody 9G6 (green). Cell nuclei are circled. Bar, 10µm. **(J, K)** HEK293 cells expressing optoFUS were exposed to 8h of blue light stimulation prior to fixation and immunofluorescence analysis of stress granule marker G3BP1 and ALS-related protein TDP-43. Arrows indicate optoFUS inclusions. Cell nuclei are circled. Bars, 10µm. **(L, M)** Human ReN neurons expressing optoFUS under the control of the human synapsin promoter (hSyn) were exposed to 72h of blue light stimulation prior to immunofluorescence analysis of FUS pathological hallmarks. Similar to inclusions formed in HEK293 cells, optoFUS inclusions in human neurons are positive for methylated FUS (9G6), negative for stress granule protein G3BP1 and negative for fellow FET family proteins TAF15 and EWSR1, suggesting a closer resemblance to ALS-FUS than FTD-FUS pathology. Arrows indicate optoFUS inclusions. Cell nuclei are circled. Bars, 10µm. Related to Figure 5.

**Figure S5.**
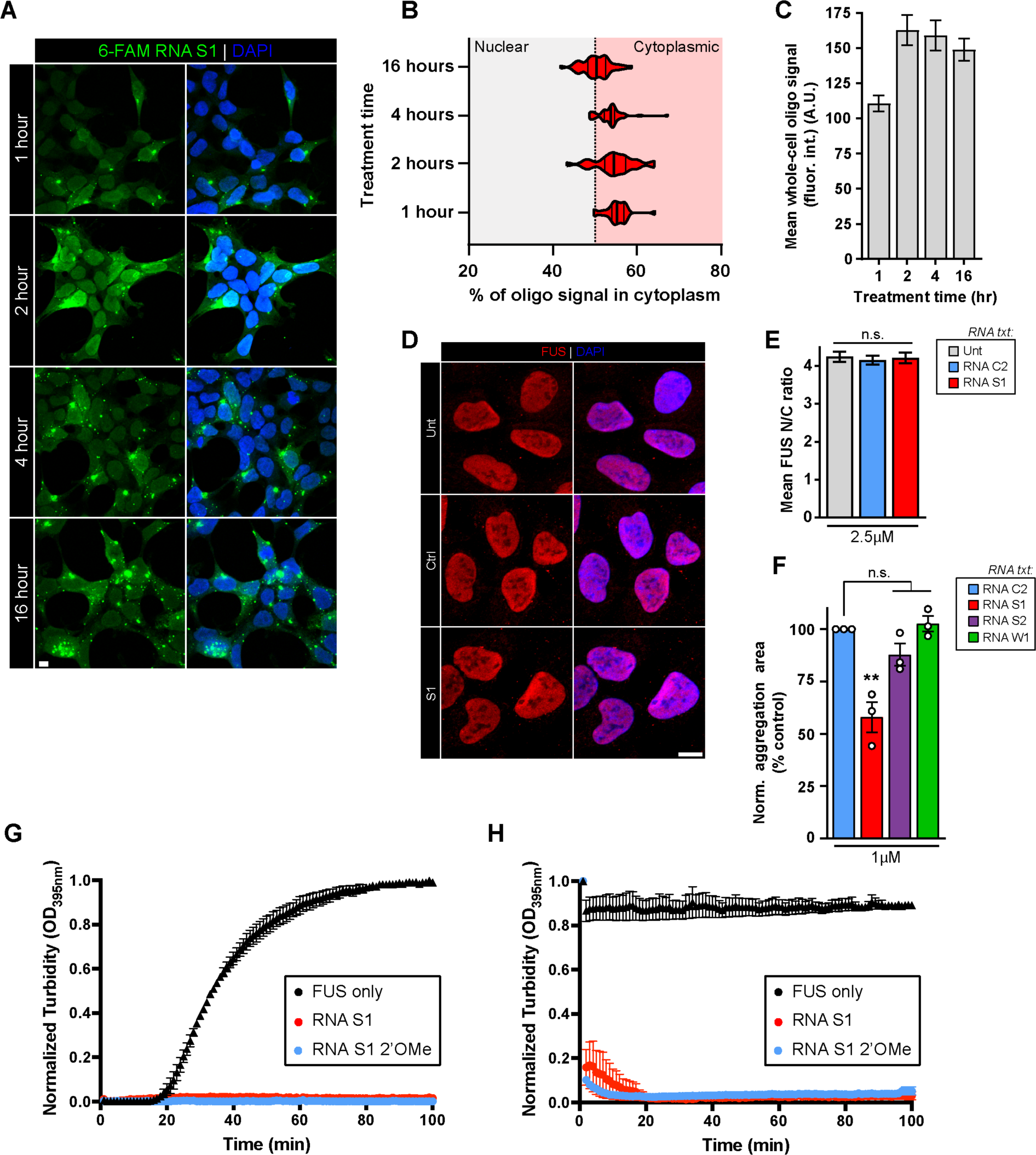
RNA S1 prevents and reverses aberrant phase transitions of FUS in human cells. **(A)** Representative images of HEK293 cells treated with 2.5µM of a 6-FAM-labeled RNA S1 for the indicated time periods. Bar, 10µm. **(B)** Quantification of percentage of 6-FAM-labeled RNA S1 signal present in the cytoplasm of cells treated for the indicated time periods. *n*=50-86 cells per treatment time. **(C)** Quantification of mean whole-cell fluorescence intensity of 6-FAM-labeled RNA S1 present within cells treated for the indicated time periods. Values represent means±SEM. *n*=34-47 cells per treatment time. **(D)** HEK293 cells were either untreated or treated with 2.5µM of RNA C2 (Ctrl) or RNA S1 for 24h prior to immunofluorescence analysis of endogenous FUS localization. Bar, 10µm. **(E)** Mean nuclear/cytoplasmic ratios of FUS fluorescence intensity in cells treated with the indicated oligonucleotides. Values represent means±SEM. *n*=41-66 cells per group. One-way ANOVA with Tukey’s post hoc test was used to compare across groups. **(F)** Normalized aggregation area of optoFUS-expressing HEK293 cells pre-treated with 1µM of RNA C2 (Ctrl), RNA S1, RNA S2, or RNA W1 for two hours prior to a 6-hour light activation period. Bars represent means±SEM. Data points represent individual experiments. *n* = 3 individual experiments, 1236-2835 cells across 9 randomized fields-of-view per experiment. One-way ANOVA with Tukey’s post hoc test was used to compare across groups; **, *p*<0.01. **(G)** GST-FUS (5µM) was incubated with TEV protease in the presence or absence of RNA S1 or 2’OMe-modifed RNA S1 analogue (20µM) for 0–100min. Fibrillization was assessed via turbidity. Values represent means±SEM (n = 3). **(H)** Fibrillization reactions were performed as in (G) for GST-FUS and at the end of the reaction, water, strong inhibitors S1 or S1 analogue (20µM) were added to the reaction. Turbidity measurements were taken every minute to assess the extent of disaggregation. Values represent means±SEM (n = 3). Related to Figure 5.

**Figure S6.**
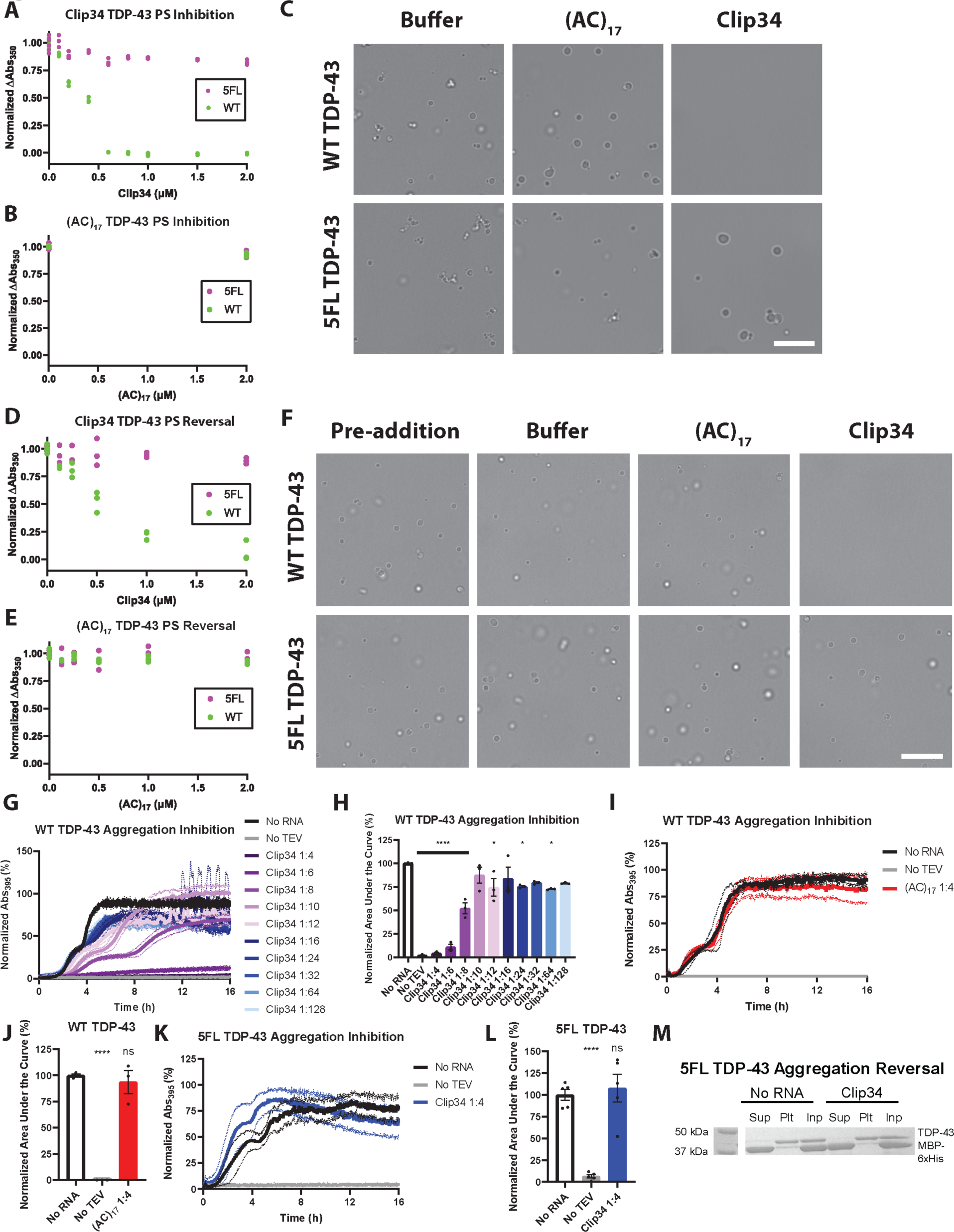
Clip34 directly prevents and reverses aberrant TDP-43 PS. (**A, B**) TDP-43 or TDP-43^5FL^ (4µM) were incubated with (A) Clip34 (0-2µM) or (B) (AC)_17_ (0-2µM) for 2h and PS was assessed via turbidity. Individual data points for 3-6 independent trials are plotted for each RNA concentration. **(C)** TDP-43 or TDP-43^5FL^ (4µM) were incubated with buffer, (AC)_17_ or Clip34 (2µM) for 2h and PS was assessed via brightfield microscopy. Bar, 10µm. **(D, E)** Preformed TDP-43 or TDP-43^5FL^ (4µM) condensates were incubated with (C) Clip34 (0-2µM) or (D) (AC)_17_ (0-2µM) for 1h and condensate integrity was assessed via turbidity. Individual data points for 3 independent trials are plotted for each RNA concentration. **(F)** Preformed TDP-43 or TDP-43^5FL^ condensates (4µM) were incubated with buffer, (AC)_17_ or Clip34 (2µM) for 1h and condensate integrity was assessed via brightfield microscopy. Bar, 10µm. **(G)** TDP-43 (5µM) was incubated in the presence of the indicated Clip34 concentration as molar ratio RNA:TDP-43. No TEV protease serves as a negative control. Fibrillization was tracked by turbidity. Values represent means. Dotted lines of corresponding colors represent ±SEM (n=3). **(H)** Area under the curve data for each replicate quantifies the extent of TDP-43 aggregation in the presence of Clip34, normalized to the no RNA condition. Values represent means±SEM (n=3). One-way ANOVA comparing to the No RNA condition; Dunnett’s multiple comparisons test; ns: p>0.05, *p adjusted ≤0.05, and ****p≤0.0001. **(I)** TDP-43 (5µM) was incubated in the presence of the indicated (AC)_17_ concentration as molar ratio RNA:TDP-43. No TEV protease serves as a negative control. Fibrillization was tracked by turbidity. Values represent means. Dotted lines of corresponding colors represent ±SEM (n=3). **(J)** Area under the curve data for each replicate summarized in (I) quantifies the extent of TDP-43 aggregation. Values represent means±SEM (n=3). One-way ANOVA comparing to the No RNA condition; Dunnett’s multiple comparisons test; ns: p>0.05, *p adjusted ≤0.05, and ****p≤0.0001). **(K)** TDP-43^5FL^ (5µM) was incubated in the presence of the indicated Clip34 concentration as molar ratio RNA:TDP-43. No TEV protease serves as a negative control. Fibrillization was tracked by turbidity. Values represent means. Dotted lines of corresponding colors represent ±SEM (n=5). **(L)** Area under the curve data for each replicate summarized in (K) quantifies the extent of TDP-43 aggregation. Values represent means±SEM (n=5). One-way ANOVA comparing to the No RNA condition; Dunnett’s multiple comparisons test; ns: p>0.05, *p adjusted≤0.05, and ****p≤0.0001. **(M)** Preformed TDP-43^5FL^ aggregates (4µM) were incubated with buffer or Clip34 (40µM) for 16h. Reactions were processed for sedimentation analysis and the supernatant fraction, pellet fraction, and input (100%) were fractionated by SDS-PAGE and Coomassie stain. Note that Clip34 is unable to return TDP-43^5FL^ to the supernatant fraction. Related to Figure 7.

**Figure S7.**
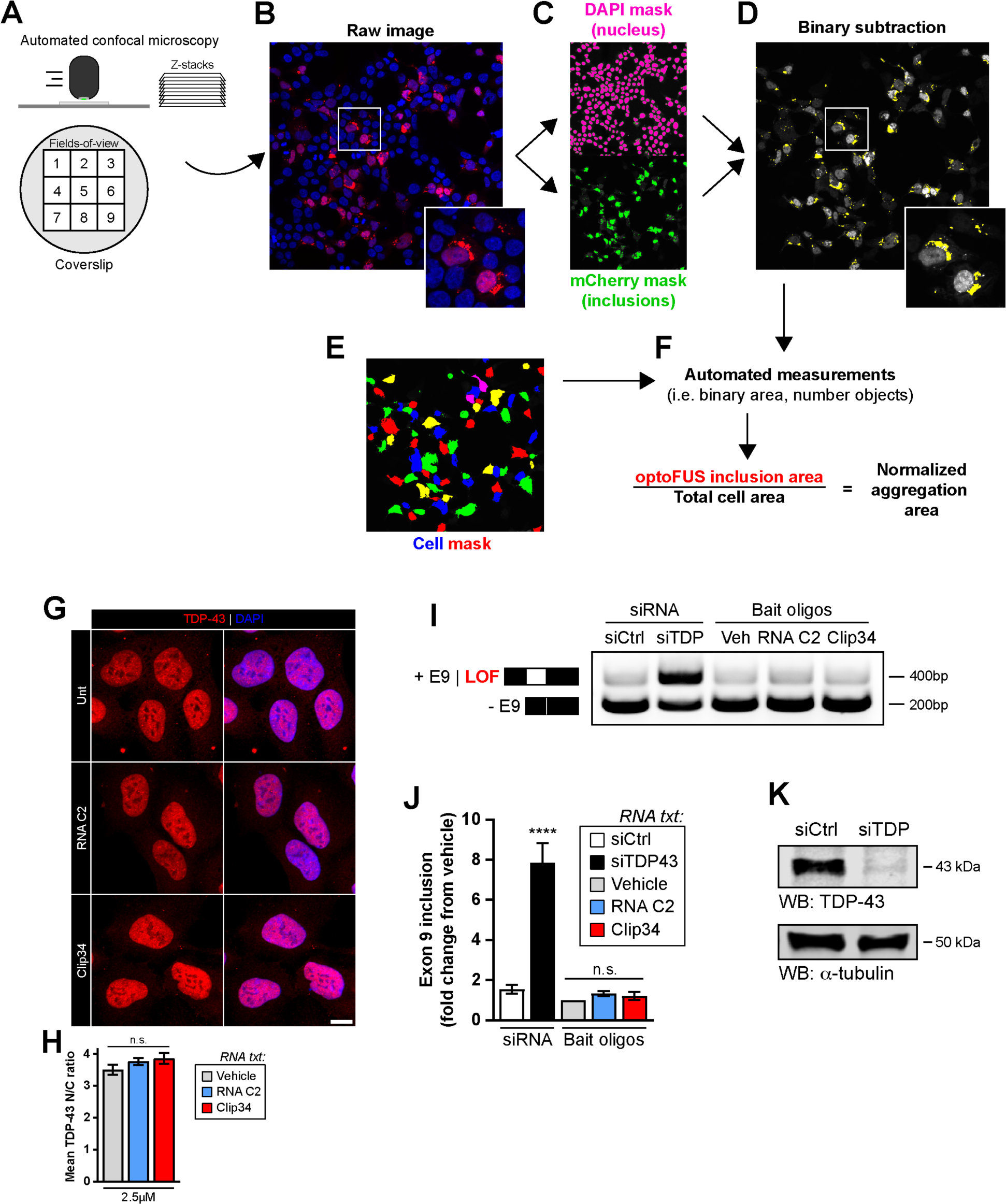
Clip34 does not affect endogenous TDP-43 localization and splicing function. **(A-F)** Automatic aggregation analysis workflow. **(G)** HEK293 cells were left untreated (Unt) or treated with 2.5µM of the indicated oligonucleotides (RNA C2 or Clip34) for 24h prior to immunofluorescence analysis of endogenous TDP-43 localization. Bar, 10µm. **(H)** Mean nuclear/cytoplasmic ratios of TDP-43 fluorescence intensity in cells treated with the indicated oligonucleotides. Values represent means±SEM. *n*=25-39 cells per group. One-way ANOVA with Tukey’s post-hoc test. **(I)** A CFTR minigene assay was used to assess endogenous TDP-43 splicing function in cells treated with the indicated siRNA (25nM) or RNA oligonucleotides (2.5µM) for 72h. Top bands indicate loss of TDP-43 splicing function (exon 9 inclusion). TDP-43 knockdown (siTDP43) was used as a positive control in these assays. **(J)** Quantification of (I). Values represent means±SEM (n=2-3). One-way ANOVA with Tukey’s post-hoc test; ****p≤0.0001. **(K)** Western blot analysis of HEK293 cells treated with 25nM of non-targeting (siCtrl) or TDP-43-targeting (siTDP) siRNA to confirm efficient TDP-43 knockdown at the time points of these experiments. Related to Figure 7.

**Table S1.**
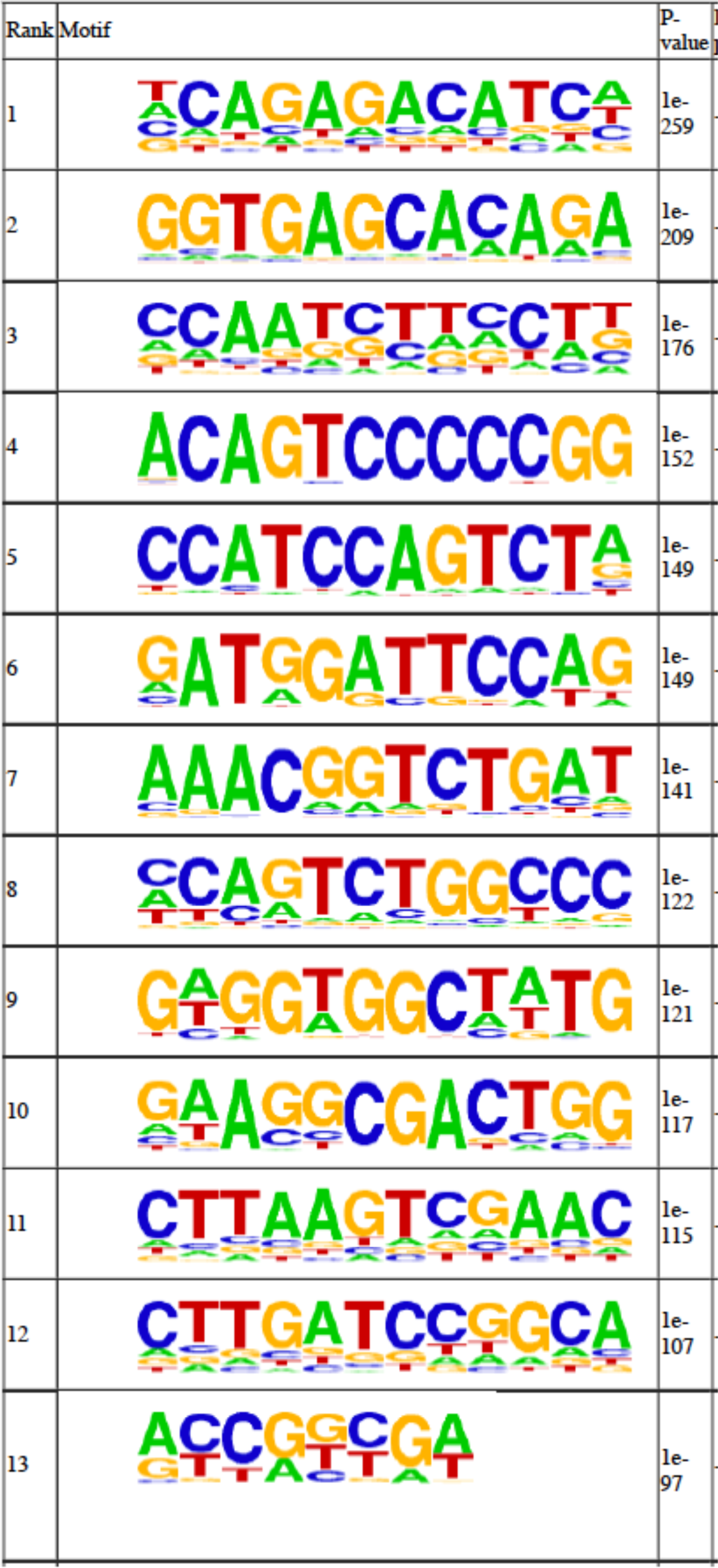
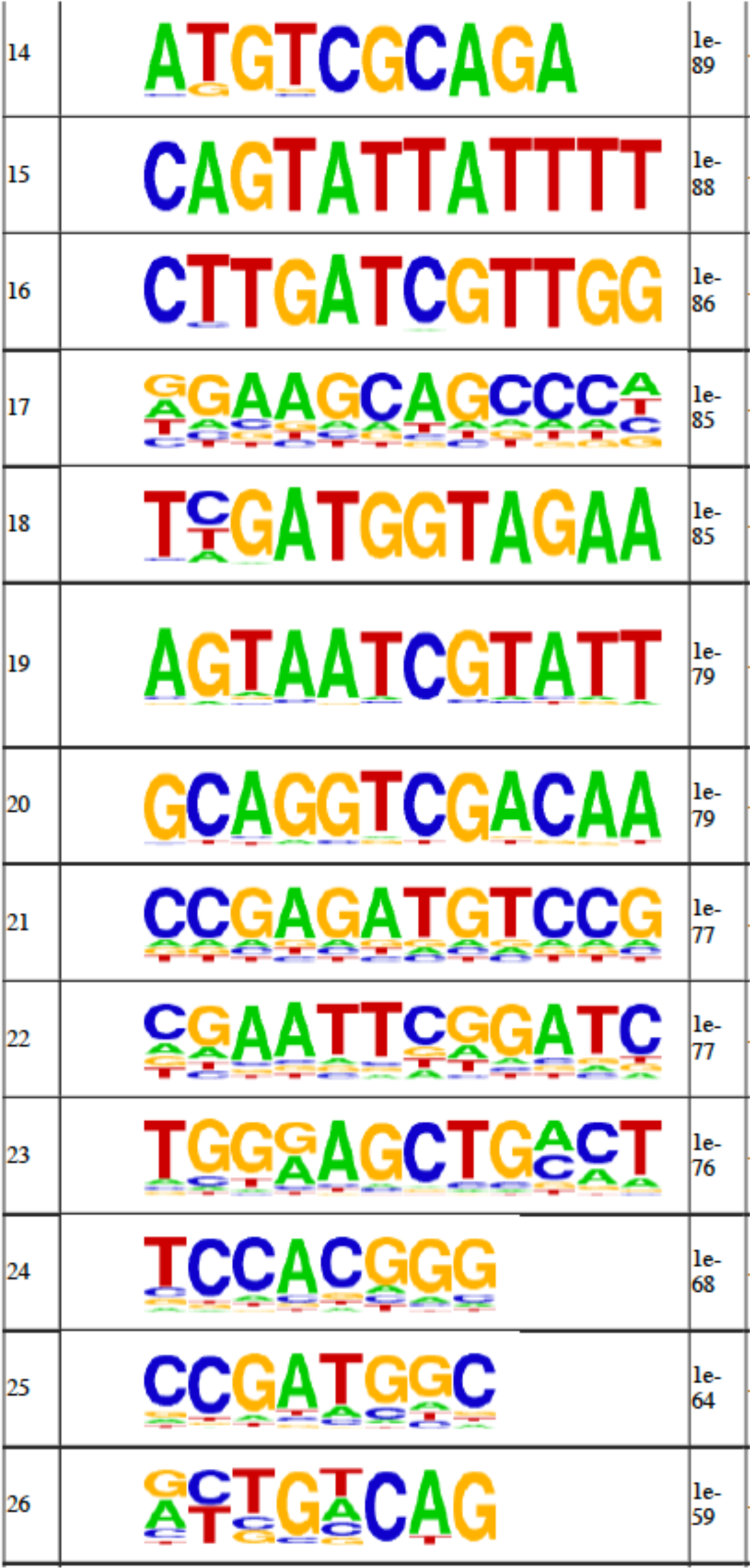
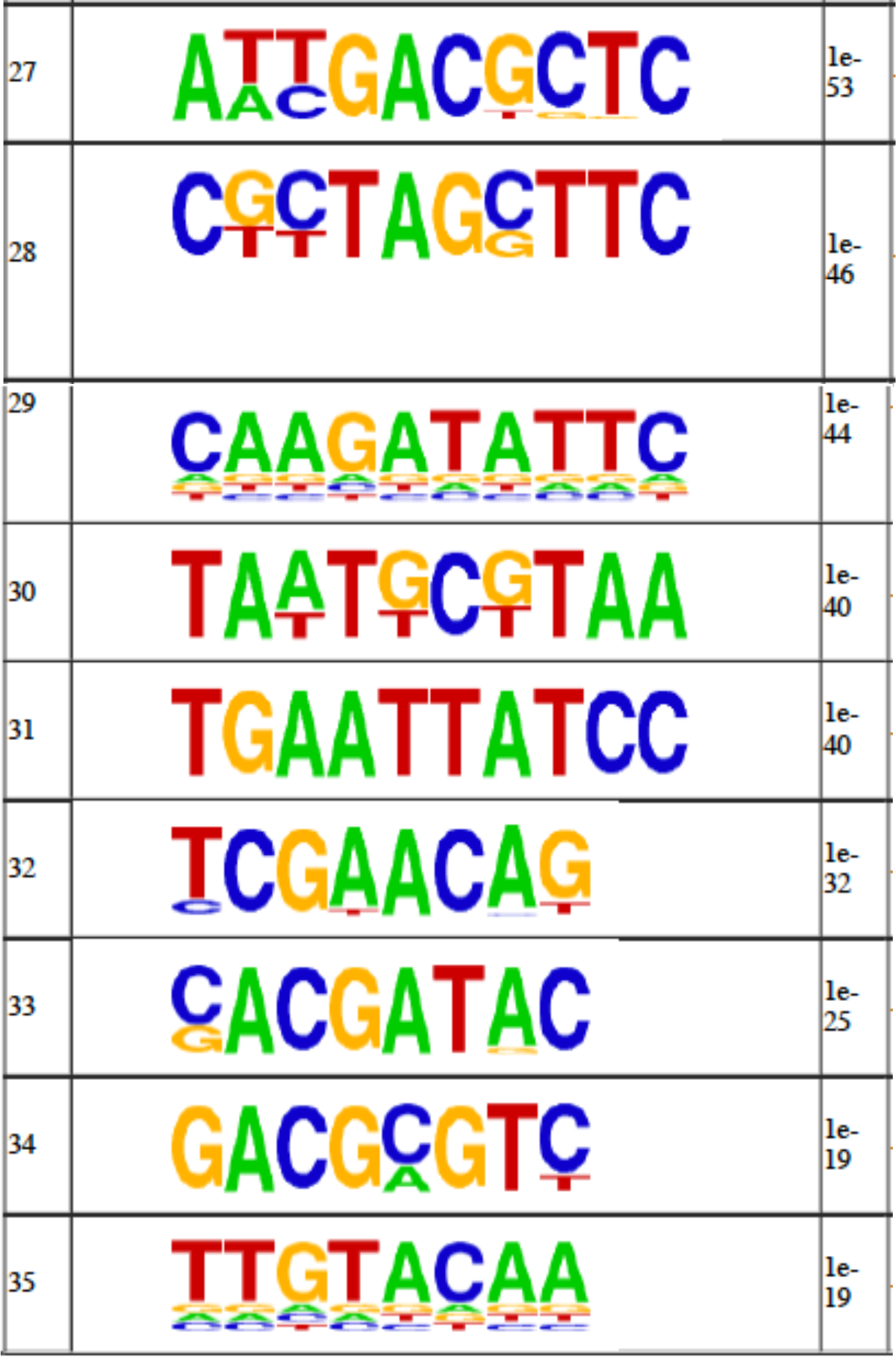
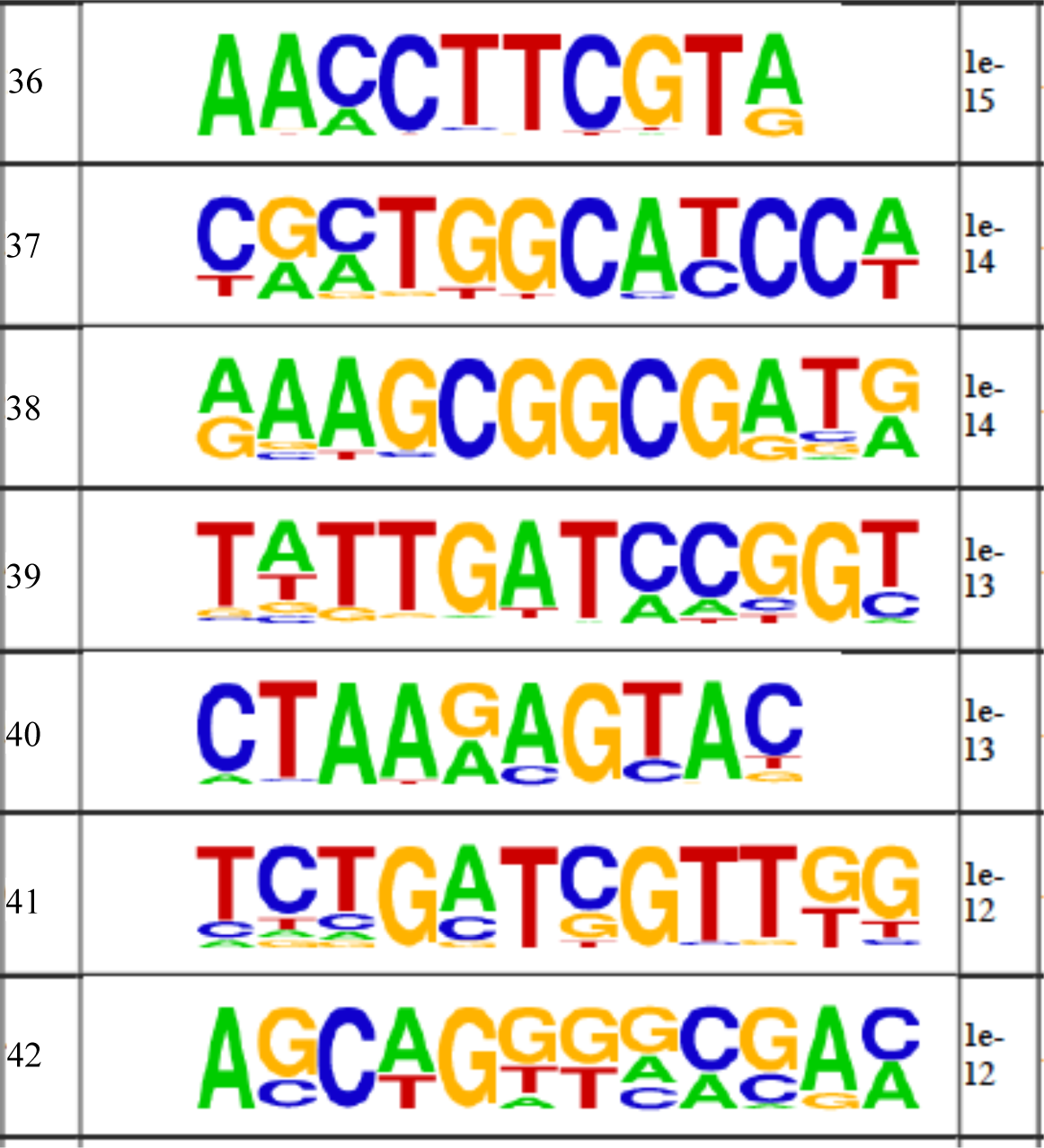
Motif analysis by Homer shows enriched sequence motifs in the FUS-binding RNA library.

**Table S2:**
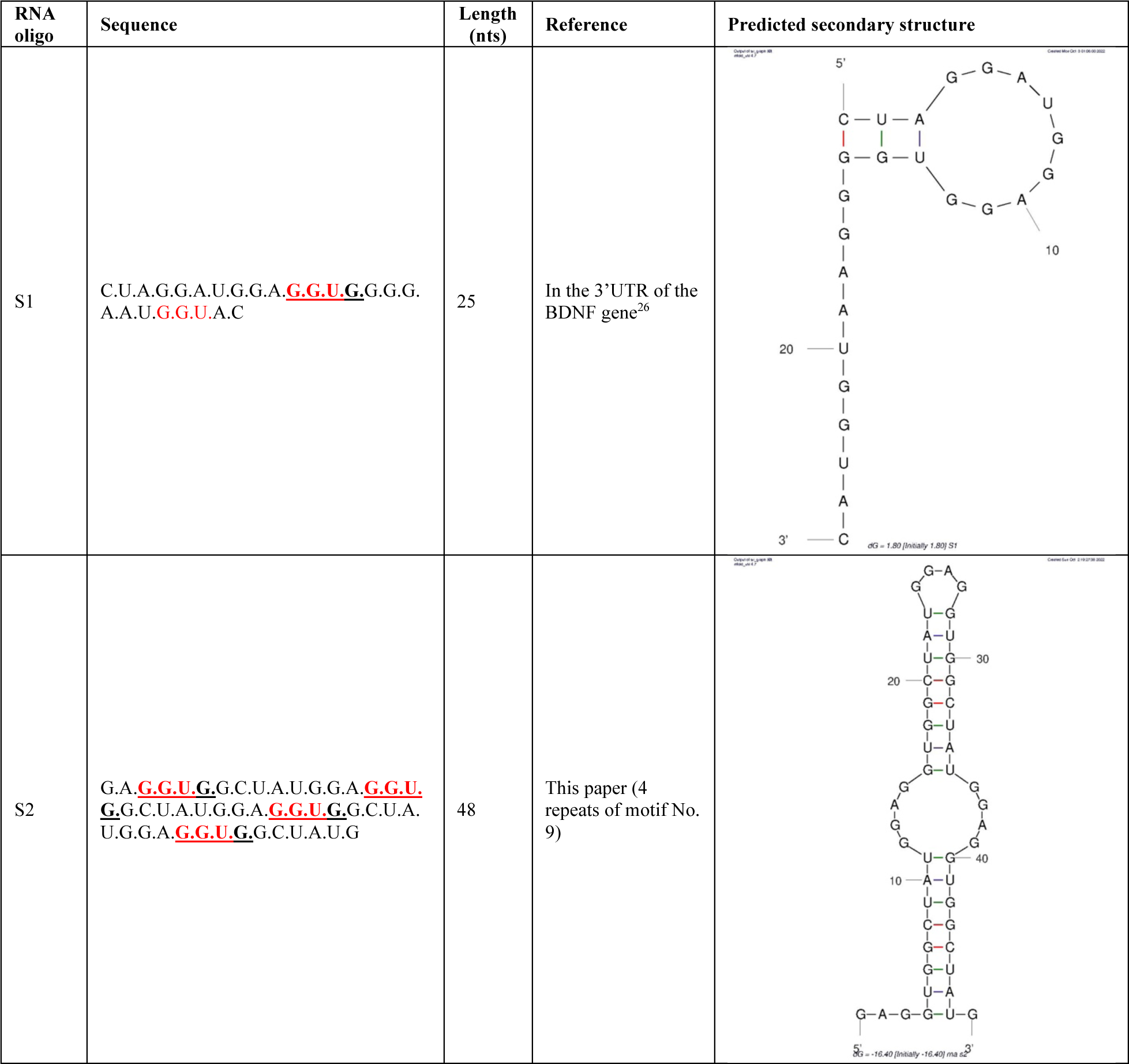
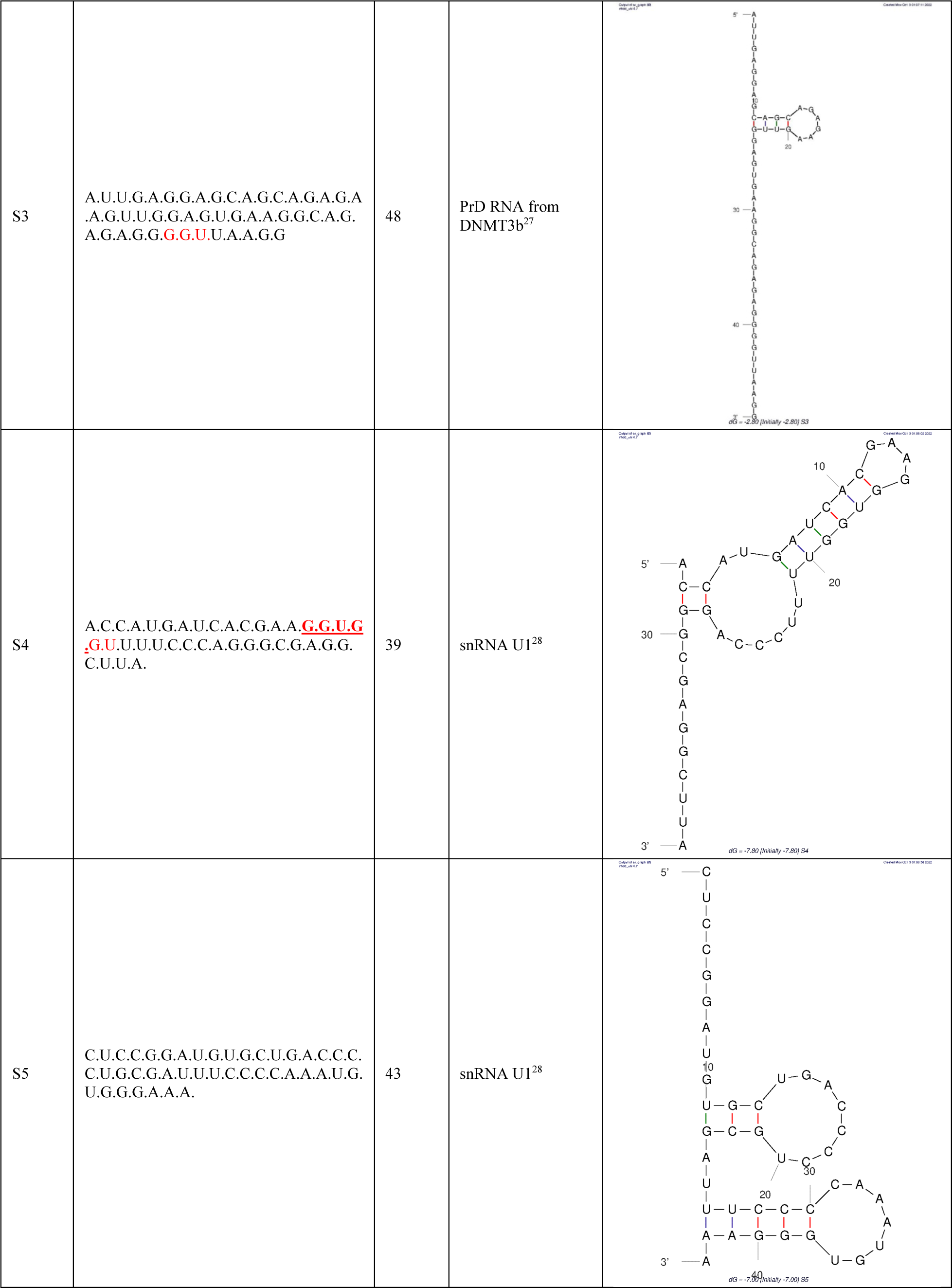
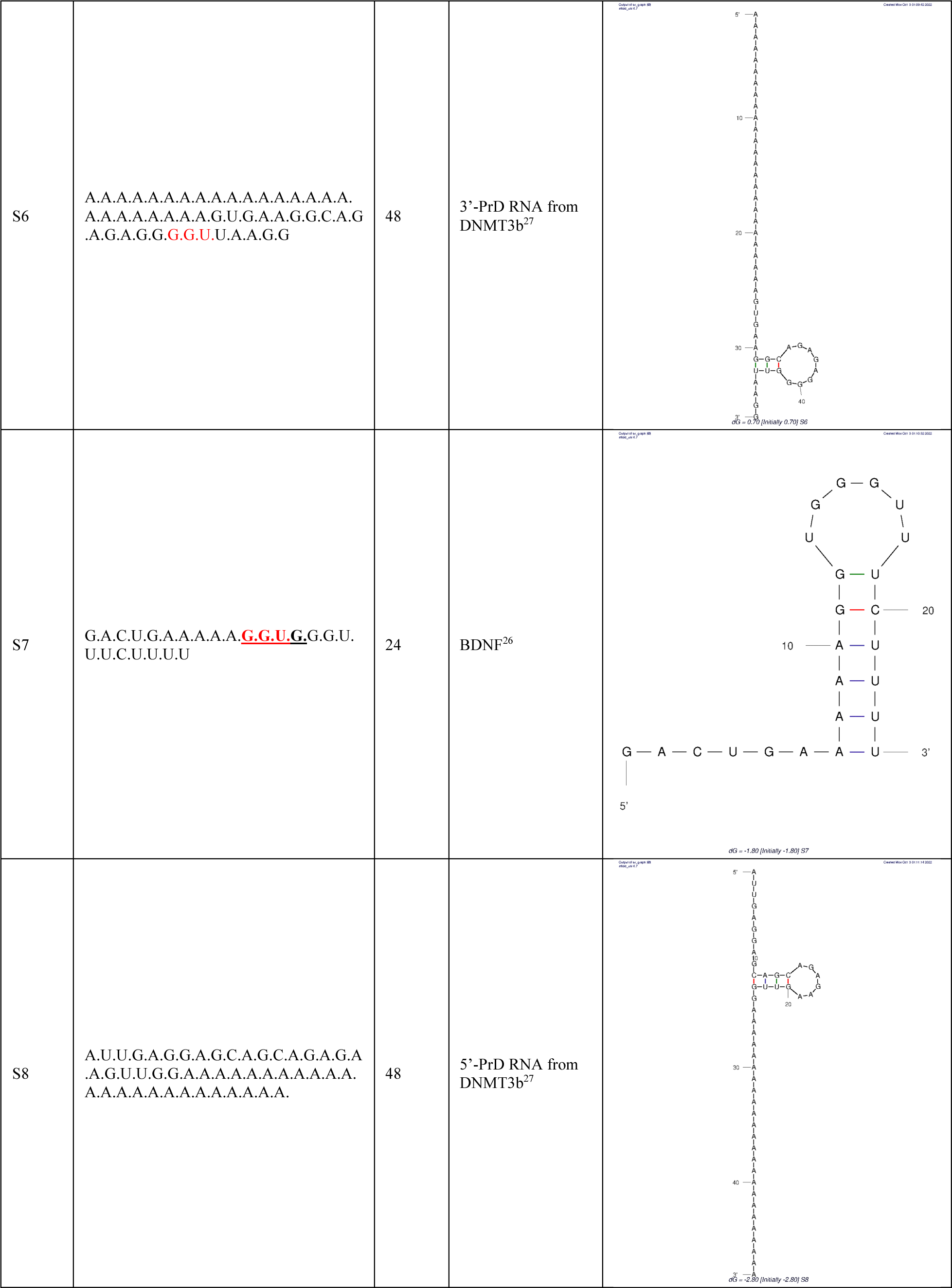
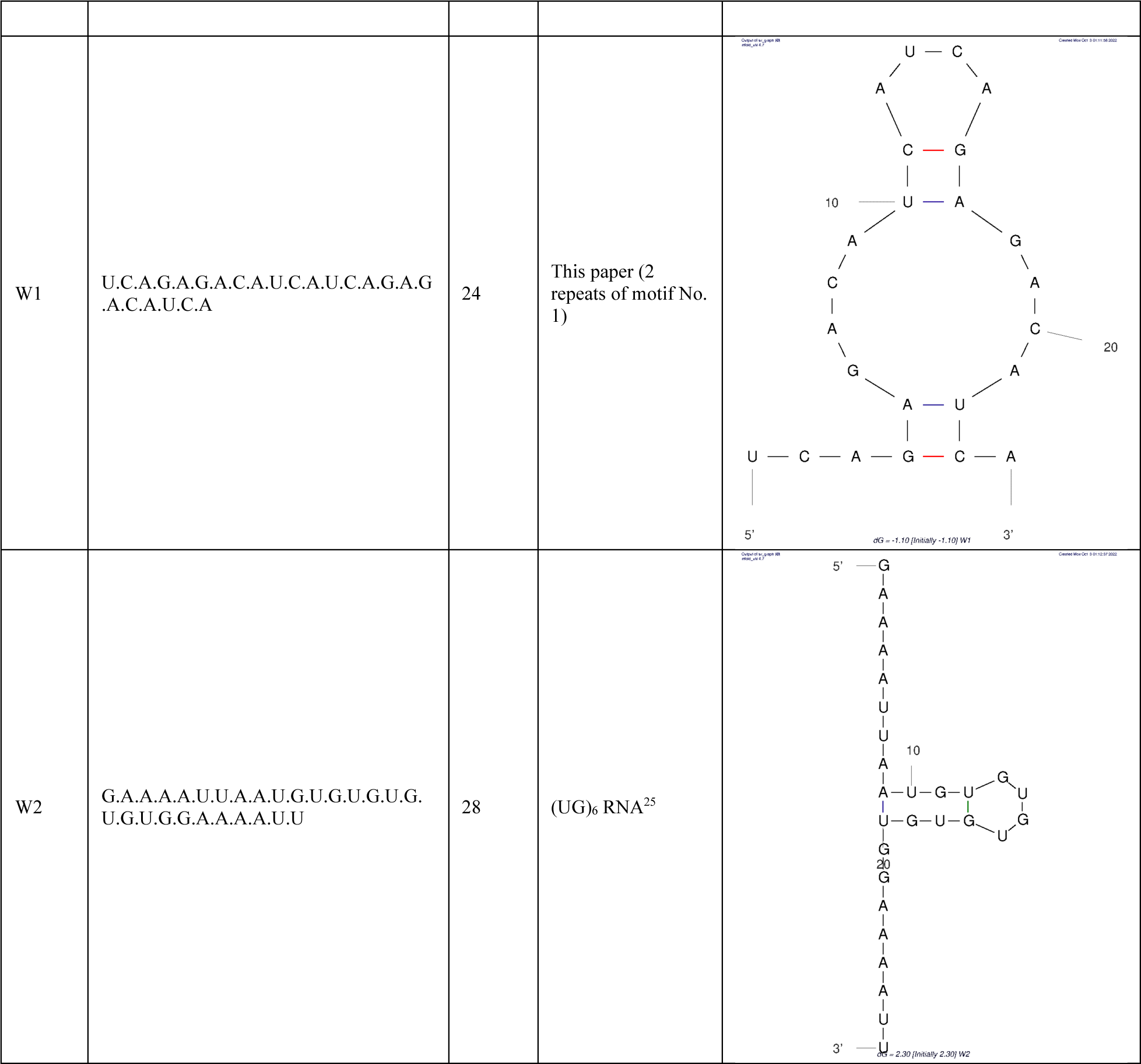
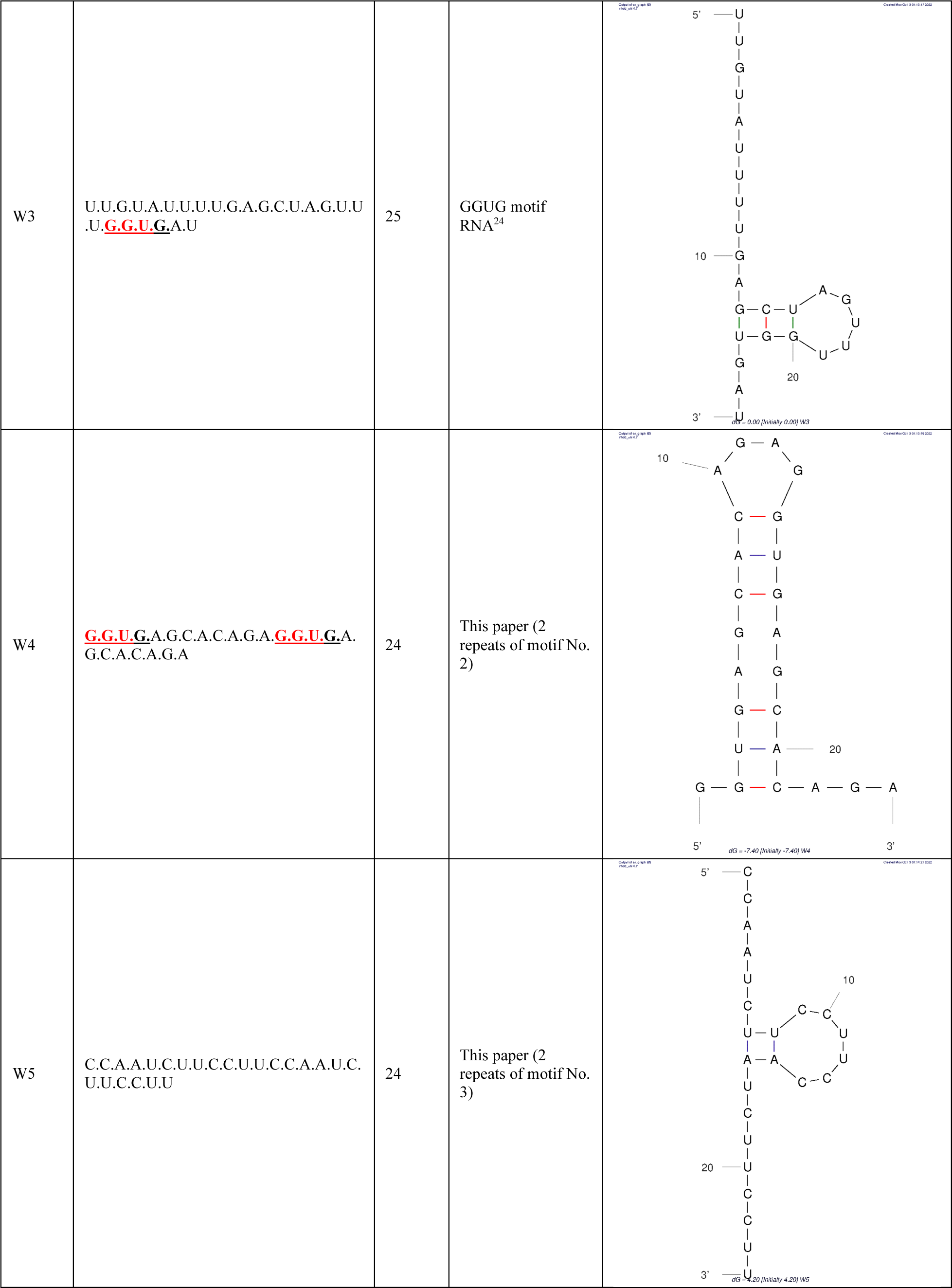
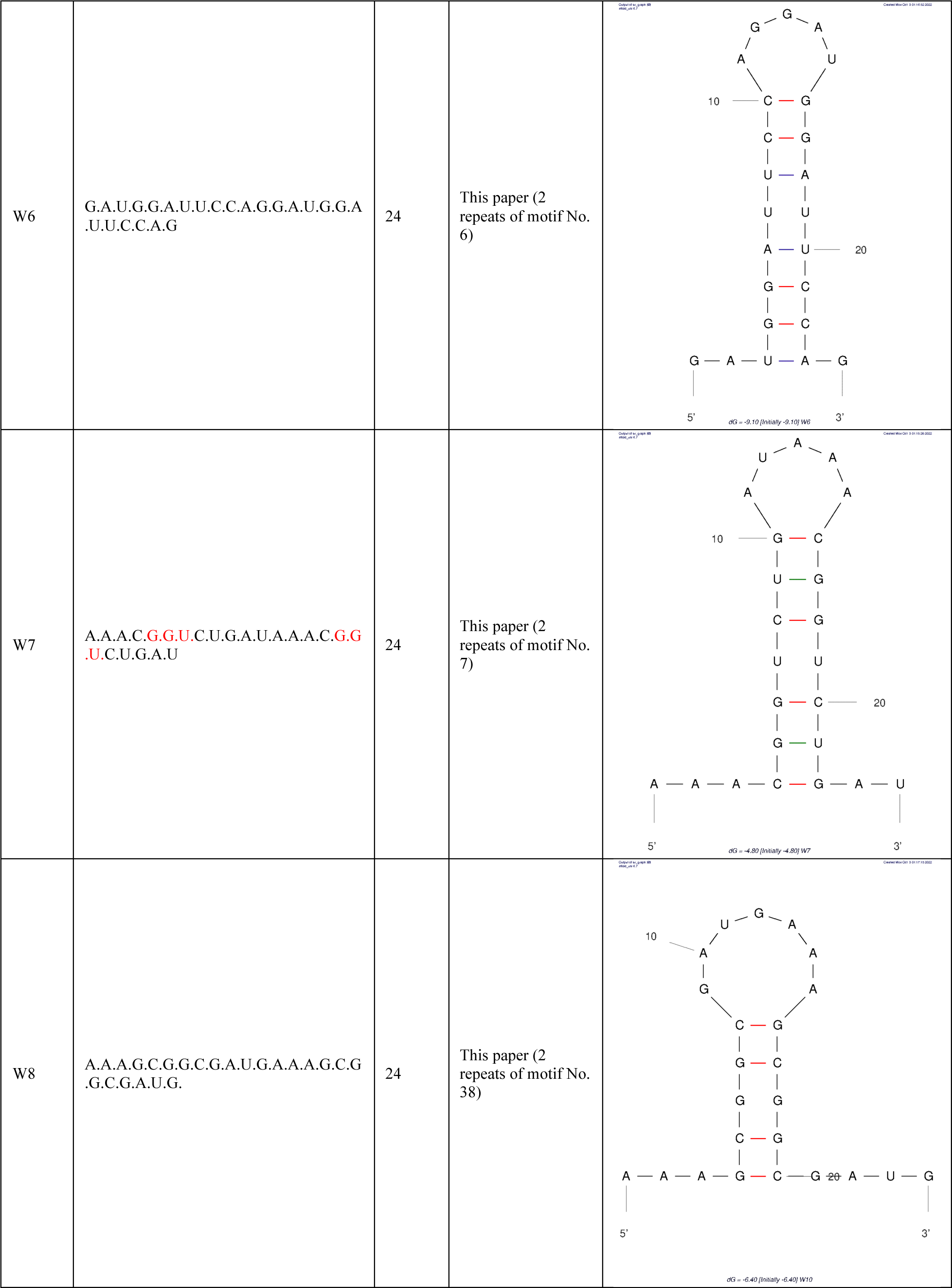
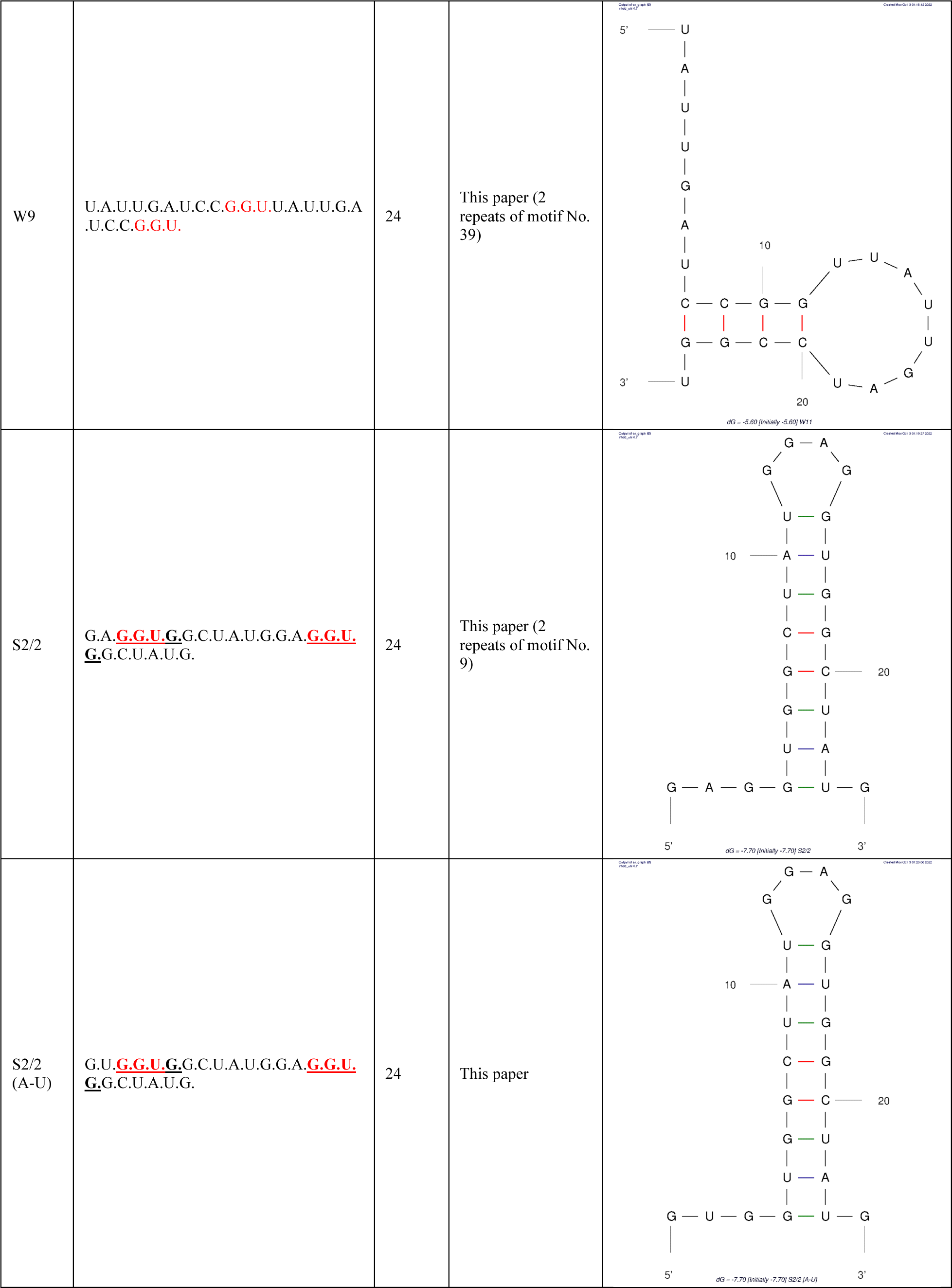
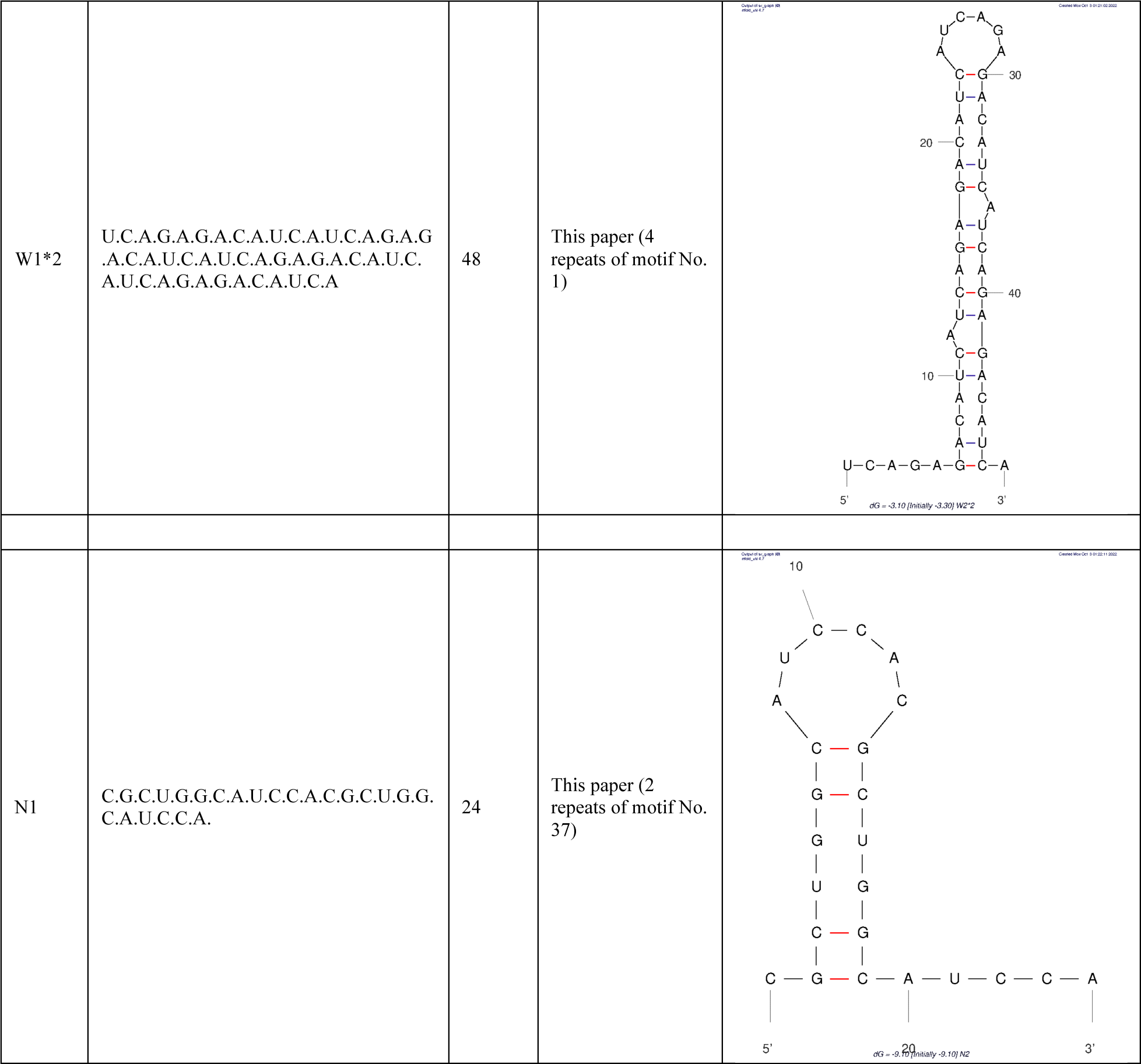
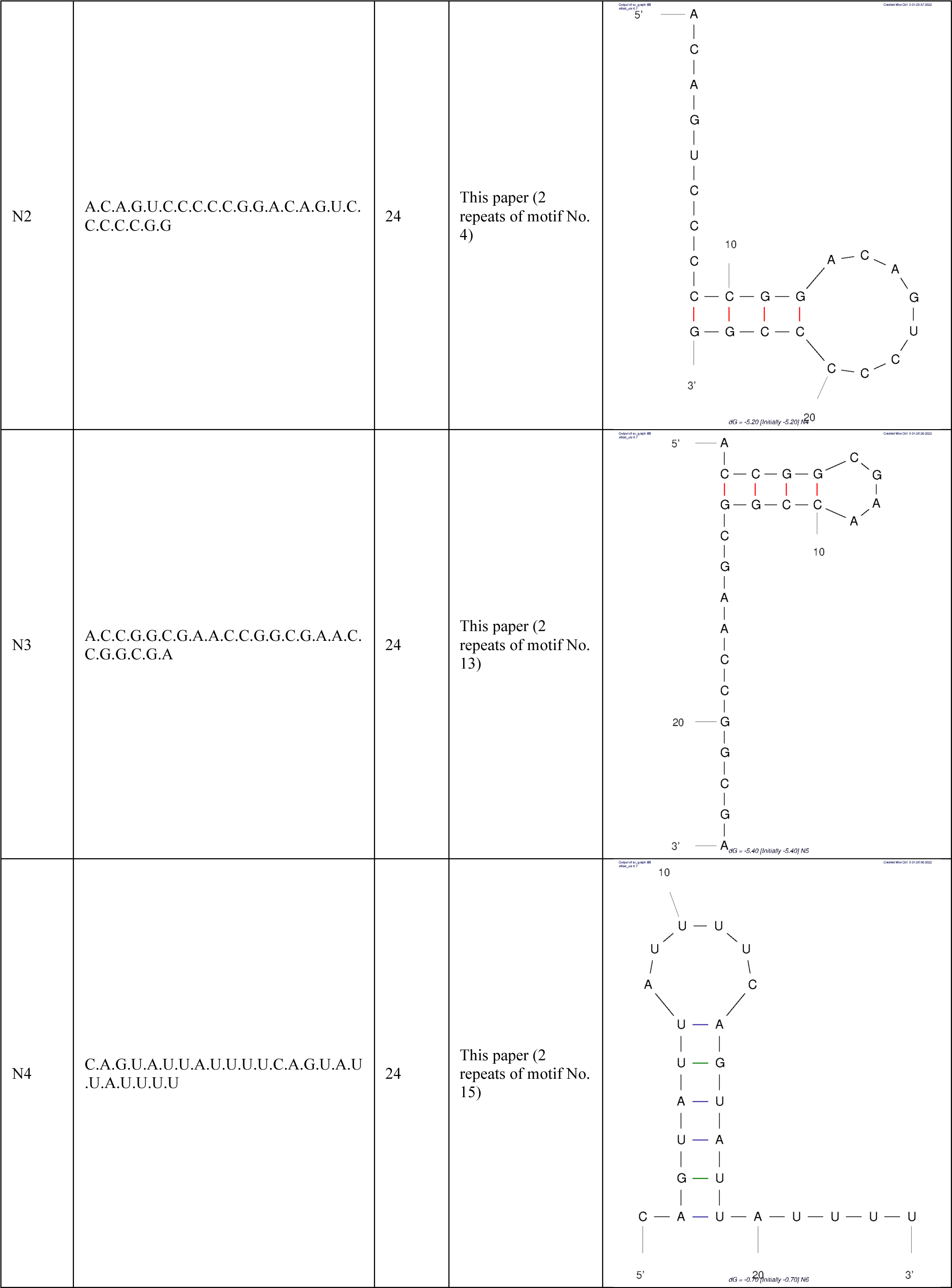
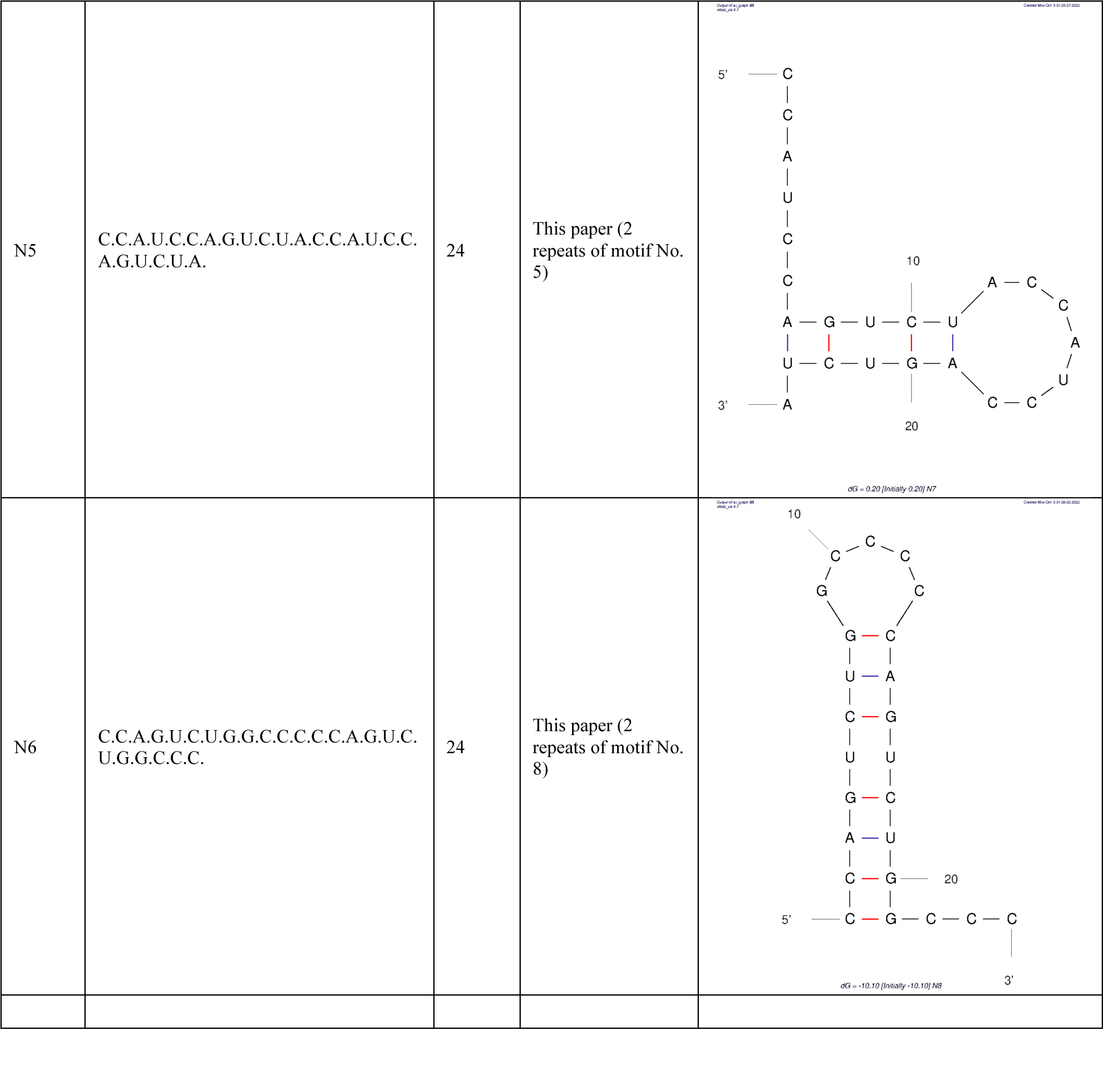
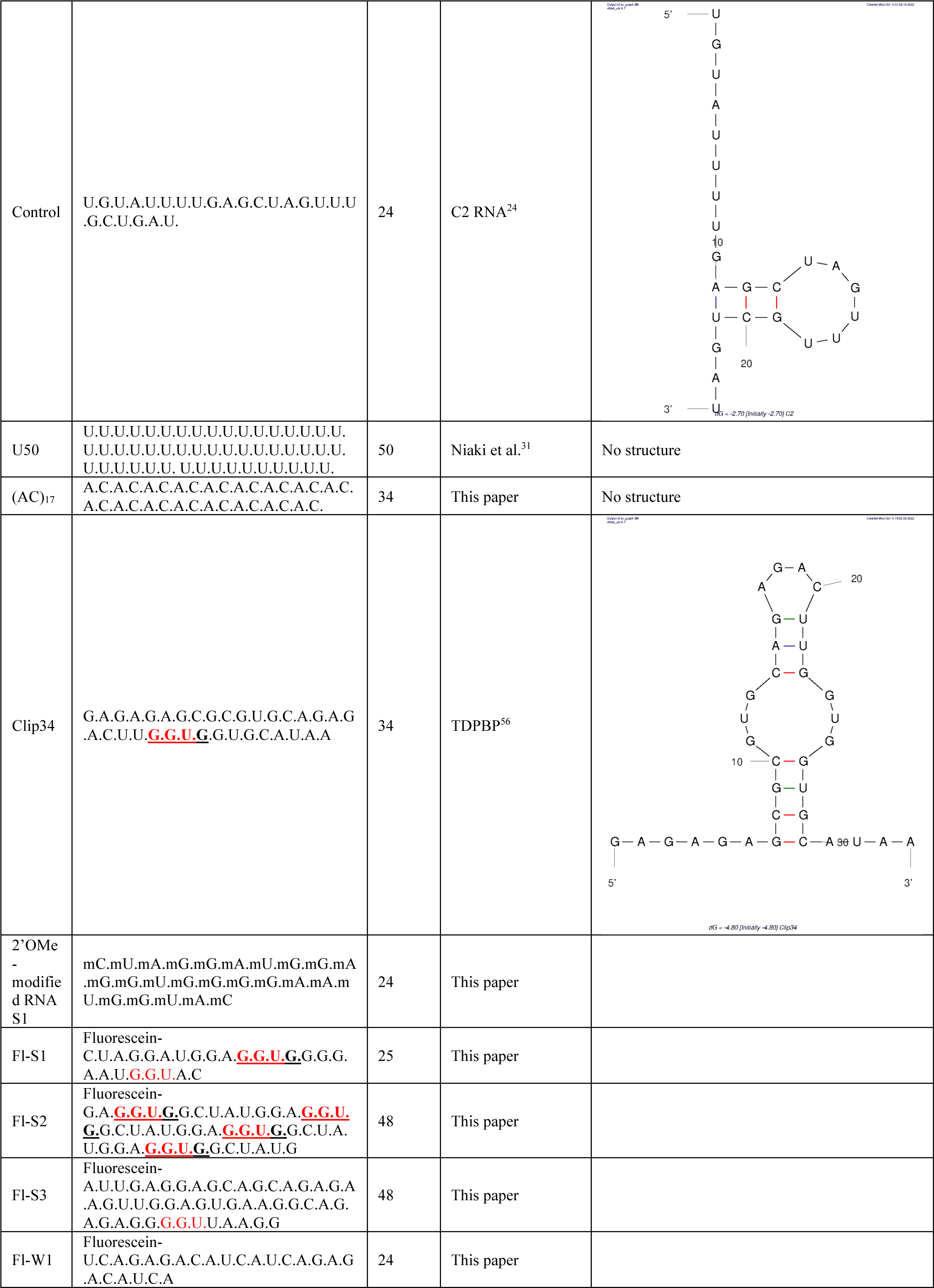
List of RNAs used in this study divided into strong inhibitors, weak inhibitors, and RNAs with no activity. Known FUS-binding motifs are highlighted as following: GGUG motifs are bolded and marked by underline and GGU motifs are highlighted in red color. Secondary structure of the RNA is predicted using M-fold ^81^. Lowest free energy secondary structure of each RNA is shown.

